# Discovery of SARS-CoV-2 M^pro^ Peptide Inhibitors from Modelling Substrate and Ligand Binding

**DOI:** 10.1101/2021.06.18.446355

**Authors:** H. T. Henry Chan, Marc A. Moesser, Rebecca K. Walters, Tika R. Malla, Rebecca M. Twidale, Tobias John, Helen M. Deeks, Tristan Johnston-Wood, Victor Mikhailov, Richard B. Sessions, William Dawson, Eidarus Salah, Petra Lukacik, Claire Strain-Damerell, C. David Owen, Takahito Nakajima, Katarzyna Świderek, Alessio Lodola, Vicent Moliner, David R. Glowacki, Martin A. Walsh, Christopher J. Schofield, Luigi Genovese, Deborah K. Shoemark, Adrian J. Mulholland, Fernanda Duarte, Garrett M. Morris

**Affiliations:** Chemistry Research Laboratory, Department of Chemistry and the Ineos Oxford Institute for Antimicrobial Research, 12 Mansfield Road, Oxford, OX1 3TA, U.K.; Department of Statistics, University of Oxford, 24-29 St Giles’, Oxford, OX1 3LB, U.K.; Centre for Computational Chemistry, School of Chemistry, University of Bristol, Cantock’s Close, Bristol, BS8 1TS, U.K.; Intangible Realities Laboratory, School of Chemistry, University of Bristol, Cantock’s Close, Bristol, BS8 1TS, U.K.; School of Biochemistry, University of Bristol, Medical Sciences Building, University Walk, Bristol, BS8 1TD, U.K.; RIKEN Center for Computational Science, 7-1-26 Minatojima-minami-machi, Chuo-ku, Kobe, Hyogo 650-0047, Japan; Diamond Light Source Ltd., Harwell Science and Innovation Campus, Didcot, OX11 0QX, U.K.; Research Complex at Harwell, Harwell Science and Innovation Campus, Didcot, OX11 0FA, U.K.; Biocomp Group, Institute of Advanced Materials (INAM), Universitat Jaume I, 12071 Castello, Spain; Food and Drug Department, University of Parma, Parco Area delle Scienze, 27/A, 43124 Parma, Italy; Univ. Grenoble Alpes, CEA, IRIG-MEM-L_Sim, 38000 Grenoble, France

## Abstract

The main protease (M^pro^) of SARS-CoV-2 is central to its viral lifecycle and is a promising drug target, but little is known concerning structural aspects of how it binds to its 11 natural cleavage sites. We used biophysical and crystallographic data and an array of classical molecular mechanics and quantum mechanical techniques, including automated docking, molecular dynamics (MD) simulations, linear-scaling DFT, QM/MM, and interactive MD in virtual reality, to investigate the molecular features underlying recognition of the natural M^pro^ substrates. Analyses of the subsite interactions of modelled 11-residue cleavage site peptides, ligands from high-throughput crystallography, and designed covalently binding inhibitors were performed. Modelling studies reveal remarkable conservation of hydrogen bonding patterns of the natural M^pro^ substrates, particularly on the N-terminal side of the scissile bond. They highlight the critical role of interactions beyond the immediate active site in recognition and catalysis, in particular at the P2/S2 sites. The binding modes of the natural substrates, together with extensive interaction analyses of inhibitor and fragment binding to M^pro^, reveal new opportunities for inhibition. Building on our initial M^pro^-substrate models, computational mutagenesis scanning was employed to design peptides with improved affinity and which inhibit M^pro^ competitively. The combined results provide new insight useful for the development of M^pro^ inhibitors.

## 1. Introduction

Severe acute respiratory syndrome coronavirus 2 (SARS-CoV-2) is the etiological agent of coronavirus disease 19 (COVID-19) that caused the World Health Organization to declare a global pandemic in March 2020. At the time of writing, >177 million cases of COVID-19 have been reported with >3.8 million deaths.^1^ A key step in maturation of SARS-CoV-2, a single-stranded positive-sense RNA virus, is the hydrolysis of its polyproteins pp1a and pp1ab. The majority of these cleavage events—at 11 sites—are performed by the SARS-CoV-2 main protease (M^pro^; also known as the 3 chymotrypsin-like or 3CL proteinase, 3C-like protease, 3CL^pro^; or non-structural protein 5, Nsp5).

M^pro^ is a homodimeric nucleophilic cysteine protease, with each protomer consisting of three domains and having a cysteine-histidine catalytic dyad (Cys-145/His-41) located near its dimeric interface.^2^ SARS-CoV-2 M^pro^ displays 96% sequence identity with SARS-CoV M^pro^ from a closely related coronavirus, which causes SARS.^3^ Dimerisation of M^pro^ is proposed to be a prerequisite for catalysis; the N-terminal “N-finger” of one protomer contributes part of the active site of the other.^4^ Indeed, the monomeric form of SARS-CoV M^pro^ is reported to be inactive.^5^ Evidence from non-denaturing mass spectrometry (MS) based assays indicates that M^pro^ monomers are not only inactive (at least with tested substrates) but also do not bind 11-mer substrate fragments with high affinity.^6^

M^pro^ and SARS-CoV M^pro^ have similar substrate specificities, both recognizing the general sequence [P4:Small] [P3:X] [P2:Leu/Phe/Val/Met] [P1:Gln] ↓ [P1’:Gly/Ala/Ser/Asn], where “Small” denotes a small residue (Ala, Val, Pro or Thr), “X” denotes any residue, and “↓” indicates the scissile amide (**Figure 1**).^7,8^ These positions in the substrates are referred to as P4-P1’, and are recognised by the corresponding subsites S4-S1’ on M^pro^. In part because these sequences are not known to be recognised by a human protease, M^pro^ is an attractive drug target.^4^ Although no clinically approved M^pro^ drugs are available, several small molecule inhibitors and peptidomimetics have been designed to inhibit SARS-CoV M^pro^ and more recently SARS-CoV-2 M^pro^.^9,10^ Indeed, a covalently-reacting M^pro^ inhibitor from Pfizer has recently entered clinical trials.^11,12^

**Figure 1:**
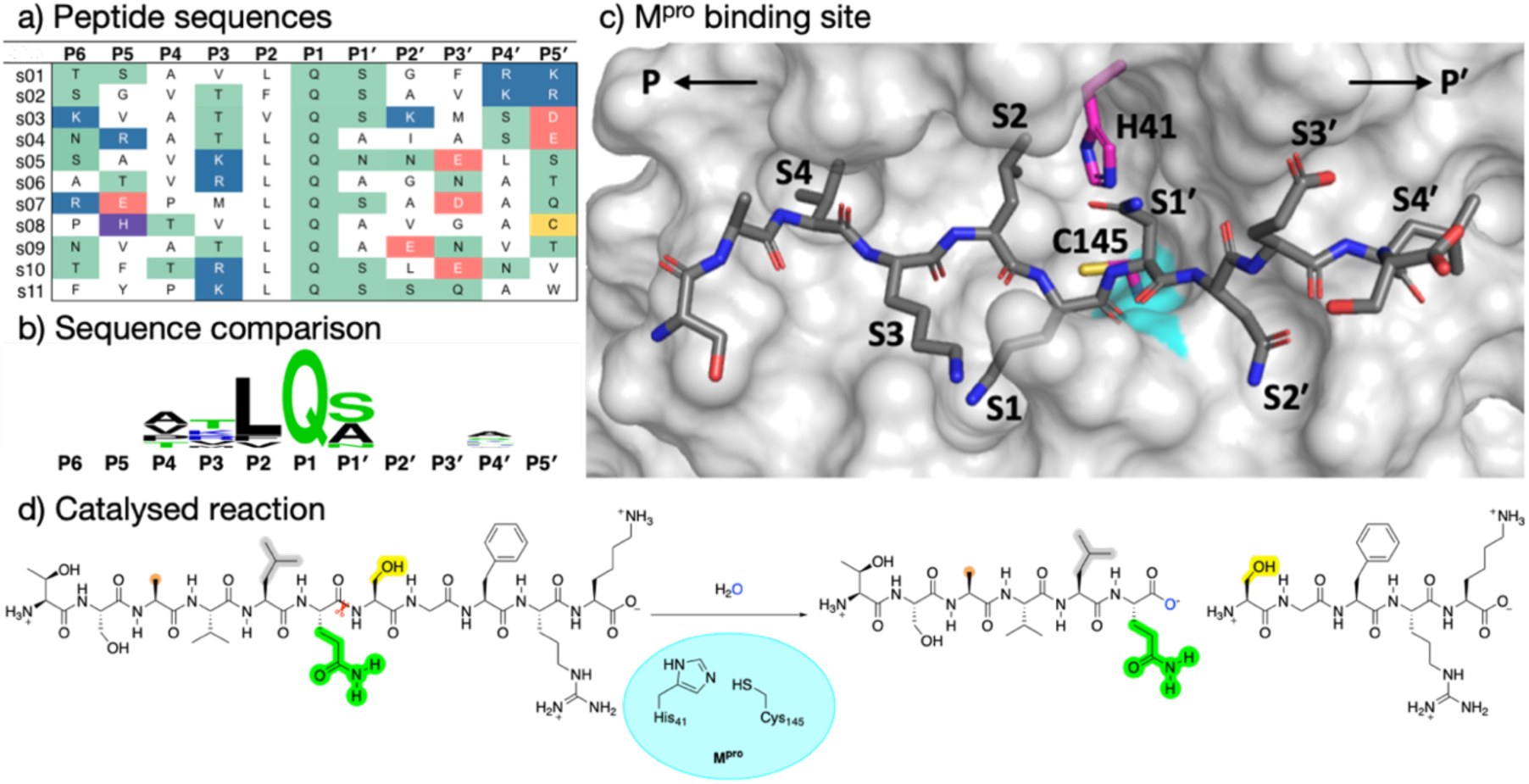
Substrates processed by SARS-CoV-2 M^pro^. a) The 11 SARS-CoV-2 M^pro^ cleavage sites, and the corresponding 11-residue fragment peptides, s01-s11, with positively/negatively charged residues in blue/red, respectively; histidine is in purple; residues with polar sidechains are in green; and cysteine is in yellow. b) Comparison between the 11 substrate sequences (generated by WebLogo)^36^ highlighting the conserved Gln at P1 and the Leu predominant at P2. c) View of an energy minimised model, built using *apo* M^pro^ (PDB: 6yb7, light grey surface)^37^, of M^pro^ complexed with s05 (dark grey sticks); subsites S4-S4’ are labelled. The oxyanion hole formed by the M^pro^ backbone NHs of Gly-143, Ser-144 and Cys-145 is in cyan. d) The reaction catalysed by M^pro^ exemplified by s01. Substrate residues important in recognition (see main text) are highlighted.

Multiple crystallographic and computational modelling studies concerning the M^pro^ mechanism^13-16^ and inhibition are available.^17-22^ (For a list of modelling studies on M^pro^ and related coronaviruses, see the CORD-19 database^23^). It is proposed that during M^pro^ catalysis, His-41 deprotonates the Cys-145 thiol, which then reacts with the carbonyl of the scissile amide to give an acyl-enzyme intermediate stabilised by a hydrogen bond network that includes the scissile amide carbonyl in an ‘oxyanion hole’. The C-terminal part of the product likely leaves the active site at this stage. The acyl-enzyme intermediate is subsequently hydrolysed with loss of the N-terminal product regenerating active M^pro^. Computational and mechanistic studies on SARS-CoV M^pro 24-27^ and SARS-CoV-2 M^pro 28-30^ suggest that His-41 and Cys-145 are most likely neutral in the active site and that the protonation states of other histidines nearby (*e*.*g*. His-163, 164, and 172) affect the structure of the catalytic machinery — although it has been suggested that the protonation state of the catalytic dyad may change in the presence of an inhibitor or substrate in SARS-CoV M^pro^.^31^ However, a different picture has been obtained from neutron crystallographic analysis of M^pro^ in the absence of a substrate or inhibitor, which indicates that the ion pair form of the dyad is favoured at pH 6.6.^32^ While neutron crystallography, in principle, enables the direct determination of hydrogen atom positions, questions remain about how pH and the presence of active site-bound ligands influence the precise and likely dynamic protonation state of the dyad.

Key questions remain regarding M^pro^ catalysis, including to what extent the active site protonation state, solvent accessibility, and substrate sequence influence activity. The lack of this knowledge makes it difficult to carry out effective computational studies on M^pro^ catalysis and inhibition.

With the aim of helping to combat COVID-19, in April 2020 we embarked on a collaborative effort, involving weekly virtual meetings, to investigate the relationship between M^pro^ substrate selectivity and activity. We employed an array of classical molecular mechanics (MM) and quantum mechanical (QM) techniques, including non-covalent and covalent automated docking, molecular dynamics (MD) simulations, density functional theory (DFT) and combined quantum mechanics/molecular mechanics (QM/MM) modelling and interactive MD in virtual reality (iMD-VR). The results provide consensus atomic-level insights into the interactions of M^pro^ with 11-residue peptides derived from the 11 natural cleavage sites (named “s01” to “s11”, in order of occurrence in the polyprotein, **Figure 1a**). The identification of key interactions between M^pro^ and its substrates, together with analysis of fragment/inhibitor structures,^33^ led to design of peptides proposed to bind more tightly than the natural substrates, several of which inhibit M^pro^. The results are freely available via GitHub (https://github.com/gmm/SARS-CoV-2-Modelling).

## 2. Results and Discussion – Understanding substrate binding and recognition

### 2.1 Protonation state of the catalytic dyad

#### 2.1.1 QM/MM studies of proton transfer in the catalytic dyad

It is proposed that M^pro^ has a neutral Cys-His catalytic dyad embedded in a chymotrypsin-like fold.^34^ To investigate the protonation state of the dyad following substrate binding, we studied proton transfer (PT) in the dyad with s01-bound (**Figure 1a**) M^pro^ using QM/MM umbrella sampling simulations at the DFTB3/MM and ωB97X-D/6-31G(d)/MM levels of theory. A 2.5 Å resolution structure of the H41A inactivated mutant of SARS-CoV M^pro^ complexed with the peptide TSAVLQ↓SGFRK (PDB entry 2q6g)^35^ was used to model SARS-CoV-2 M^pro^ bound to the same peptide, s01 (**Figure 1a**). His-41 was treated as N*δ*-protonated in the neutral state of the dyad, as reported by Pavlova *et al*. to be preferred for both *apo* and N3 inhibitor-bound M^pro^ based on MD studies.^28^

Cys-145 was treated as neutral, with its Sγ protonated. We investigated the effect of varying the protonation state of the dyad-neighbouring residue His-163, which interacts with Tyr-161, Phe-140 and the sidechain of the substrate P1 Gln. Three His-163 protonation states were considered: (i) N*δ*-protonated neutral, or “HID”; (ii) N*ε*-protonated neutral, or “HIE”; and (iii) both N*δ* and N*ε*-protonated and positively charged, or “HIP”, using AMBER forcefield naming.^38^ Both the forwards and backwards PT processes were simulated.

DFTB3/MM free energy profiles showed significant hysteresis, with the neutral (N) dyad preferred in the forward direction (from neutral to zwitterionic state), and the zwitterionic state being preferred in the backward direction (**Figure S2.1**). This hysteresis likely arises from different conformations adopted by the charged Cys-145 thiolate in the zwitterionic (or ion pair, IP) state, and interactions between the thiolate and surrounding residues (**Figure 2a, b**), in particular His-163. For all three His-163 protonation states, the forward trajectories (N to IP) showed the anionic Cys-145 to be stabilised solely by interaction with positively charged His-41 (**Figures 2a** and **S2.2A-C**). In this arrangement, the thiolate is rotated away from the scissile carbonyl during PT resulting in a poor orientation for nucleophilic attack. This suggests that such a zwitterionic state is transient, with a concerted proton transfer and simultaneous nucleophilic attack of the thiolate onto the scissile amide carbon being more likely than a stepwise mechanism.^13^ By contrast, the backwards PT trajectories (from IP to N) showed stabilisation of the Cys-145 thiolate in addition to that provided by His-41. In the case of HID-163 and HIP-163, the thiolate was stabilised by interactions with the His-163 N*δ* proton and the P1 Gln backbone N-H (**Figure S2.2E, F**). For HIE-163, additional thiolate stabilisation came from interactions with the backbone N-H and the hydroxyl of the P1’ Ser, and an additional water that diffused into the active site (**Figures 2b** and **S2.2D**). Such hysteresis effects are common in simulations of proton/charge transfer,^39^ and reflect the fact that the simulation is effectively biased towards the starting structure, or possibly limited exploration of distinct free energy basins.

**Figure 2:**
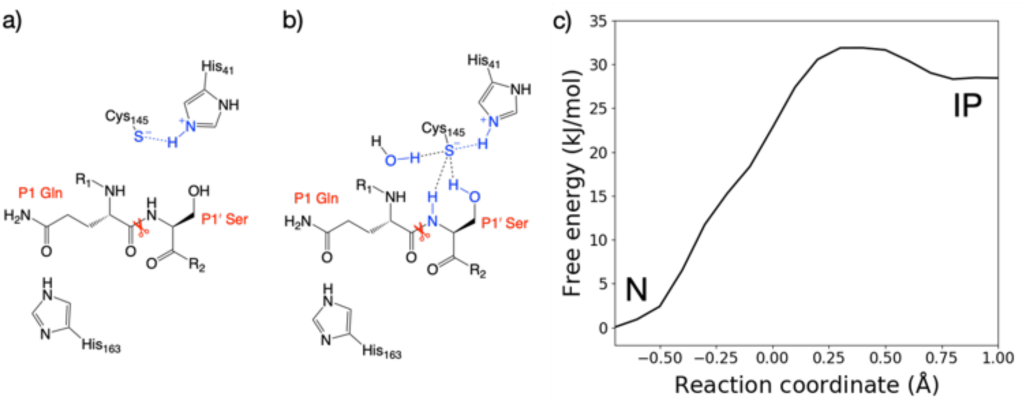
QM/MM umbrella sampling of the proton transfer in the catalytic dyad. Representation of the interactions of the Cys-145 thiolate in the ion pair (IP) state from the a) forwards and b) backwards simulations. c) Free energy profile for interconversion of the neutral (N) and ion pair states of the dyad in the HIE-163 system, from the combined forwards and backwards QM/MM umbrella sampling MD simulations, corrected to the ωB97X-D/6-31G(d)/MM level of theory.

The zwitterionic state with HID-163 was less stable with respect to the neutral state than the zwitterionic states with HIE-163 and HIP-163 (**Figure S2.1C**). This was due to a His-163 sidechain rearrangement in the backwards PT trajectory that results in breaking the N*δ*-H hydrogen bond with Tyr-161 and its π-π stacking interaction with Phe-140, suggesting that a N*δ*-protonated His-163 is unlikely. Double protonation of His-163 results in a loss of both interactions in the forwards and backwards PT trajectories. Despite both HIP-163 and HIE-163 giving similar PT free energy profiles, the loss of these interactions suggests HIP-163 is unfavourable for productive catalysis. These QM/MM results therefore suggest that an N*ε*-protonated neutral His-163 is most likely. Along with conserving interactions with Tyr-161 and Phe-140, an N*ε*-protonated His-163 also formed a hydrogen bond with the P1 Gln side chain (**Figures 2a, b**), an interaction not observed in PT trajectories with HID-163 and HIP-163.

It is known that DFTB3 overestimates the proton affinity of methylimidazole (similar to a histidine side chain) by 30.1 kJ mol^-1^.^40^ Therefore, the DFTB3/MM method will over-stabilise the zwitterionic state relative to the neutral state. To account for this error, the backwards PT reaction with a N*δ*-protonated His-163 was modelled at the ωB97X-D/6-31G(d)/MM level of theory. This showed the zwitterionic state was 24.3 kJ mol^-1^ above the neutral state, an increase of 26.4 kJ mol^-1^ compared to DFTB3/MM (**Figure S2.3**). Applying the free energy difference between ωB97X-D/6-31G(d) and DFTB3 at each reaction coordinate value as a correction to the combined QM/MM free energy profile in the case of HIE-163, the neutral catalytic dyad is preferred, with the ion pair being 28.5 kJ mol^-1^ higher in energy than the neutral state (**Figure 2c**).

#### 2.1.2 Tautomeric and conformational states of other histidine residues

With the neutral form of the dyad and an N*ε*-protonated neutral His-163 established, we used the highest resolution contemporary structure of M^pro^ available (PDB entry 6yb7, 1.25 Å resolution),^37^ for simulations of M^pro^ in complex with its 11 peptide substrates. The protonation state assignments of other histidines are given in **Table S2.1**. Our assignments agree with those of Pavlova *et al*. for the N3 inhibitor-bound complex,^28^ and the fluorescent tag-containing polypeptide-enzyme complex reported by Swiderek and Moliner.^13^

His-41 and His-164 are of particular interest. Analysis of several M^pro^ structures (both *apo* and inhibitor-bound) indicates a conserved crystallographically observed water located between His-41, His-164 and Asp-187 (**Figure S2.4**);^2,4,34,37^ this water (HOH-644 in PDB 6yb7) has been suggested to play a role in SARS-CoV M^pro^ catalysis.^27^ It has been noted that in PDB entry 6yb7, His-41 is in a distinct conformation compared to other M^pro^ structures (PDB entries 6wqf, 6y2g, and 7bqy).^28^ To facilitate equilibration of our simulations into a productive conformation, a rotation was applied to the imidazole of His-41 (N*δ*-protonated). Our preliminary MM MD simulations suggest that this rotamer, along with protonation of His-164 at N*ε* and retention of HOH-644, results in lower and less fluctuating root mean square positional deviation (RMSD) values for the His-41 and His-164 sidechains (**Figure S2.5**).

### 2.2 Models of SARS-CoV-2 M^pro^-substrate peptide complexes

To identify the key interactions with its substrates and to assess their relative binding affinities, we constructed models of SARS-CoV-2 M^pro^ complexed with its 11 cleavage site-derived substrates as 11-amino acid peptides, from P6 to P5’, by comparative modelling using the H41A mutant of SARS-CoV M^pro^ bound to an N-terminal substrate^35^ and the SARS-CoV-2 sequence from GenBank entry MN908947.3^41^ (see **Section S1.2** in **Methods** and **Figures 1a** and **S2.6**). We refer to these peptides as ‘substrates’ as their hydrolytic sites are all cleaved by M^pro^ (*vide infra*). The substrates were modelled in crystallographic chain A of the M^pro^ dimer with a neutral dyad; unless otherwise stated, all M^pro^ residue numbers and names in the following discussions refer to chain A. Initial models were subjected to three independent MD simulations each of 200 ns.

Three of the 11 cleavage site-derived peptides (s01, s02, and s05) were also modelled using interactive MD using virtual reality (iMD-VR), as an alternative to comparative modelling and traditional MD. iMD-VR provides an immersive 3D environment for users to interact with physically rigorous MD simulations. Since the simulation responds in real-time to manipulation, iMD-VR is a useful tool for guiding generation of docked structures based on chemical knowledge; *e*.*g*. if a user tries to bring two negatively charged groups together, it will endeavour to minimise the energy of the system by moving them apart.^42^ The three substrates were chosen because s01 and s02 have the highest relative efficiencies (of SARS-CoV M^pro^) of all substrates; while s05 has the second-lowest catalytic efficiency but the same P2 and P1 residues as s01.^43^ We hypothesised that an iMD-VR user may perceive a difference when docking these substrates, and that this perception might relate to catalytic efficiency. Indeed, the iMD-VR user found both s01 and s05 were easier to dock than s02, as s02 required manipulation of residues after initial docking to eliminate clashes between M^pro^ and the substrate. Three 200 ns replicates of implicit solvent MD simulation were performed on each docked structure of s01, s02, and s05. It should be noted that iMD-VR has successfully produced accurate docked structures of various drug-viral enzyme complexes, including oligopeptide- and inhibitor-M^pro^ complexes.^44,45^

Throughout both the explicit-solvent MD of all 11 substrates and the implicit-solvent MD simulations of iMD-VR structures of s01, s02, and s05, all substrates remained tightly bound in the active site (see backbone RMSD, and root mean square fluctuation or RMSF analyses in **Figures S2.7-10**). Backbone stability is maintained especially in the central region of the substrates, with only the N- and C-terminal residues showing substantial flexibility. Local sidechain fluctuations are present, notably at the P3 residue which occupies the solvent-facing S3 subsite (**Figure 1c**; RMSF plot in **Figure S2.8**). Another clear trend is that the C-terminal P’-side residues consistently fluctuate more than the N-terminal P-side residues, likely in part as a result of fewer protein-substrate hydrogen bonds on the P’ side (*vide infra*).

#### 2.2.1 Conserved hydrogen bond interactions

Crystallographic studies on SARS-CoV M^pro^ reveal the importance of hydrogen bonds in binding of substrate s01,^35^ which has the same cleavage site sequence in SARS-CoV-2 (**Figure 1a**). To investigate whether this is the case for SARS-CoV-2 M^pro^ in complex with its substrates, we analysed the prevalence of hydrogen bond (HB) interactions at each subsite for all 11 substrates, in both explicit-solvent MD and implicit-solvent MD simulations of iMD-VR docked structures. Twelve key HBs were identified (**Figure 3**) and monitored, with distance and angle distributions shown in **Figure S2.11**.

**Figure 3:**
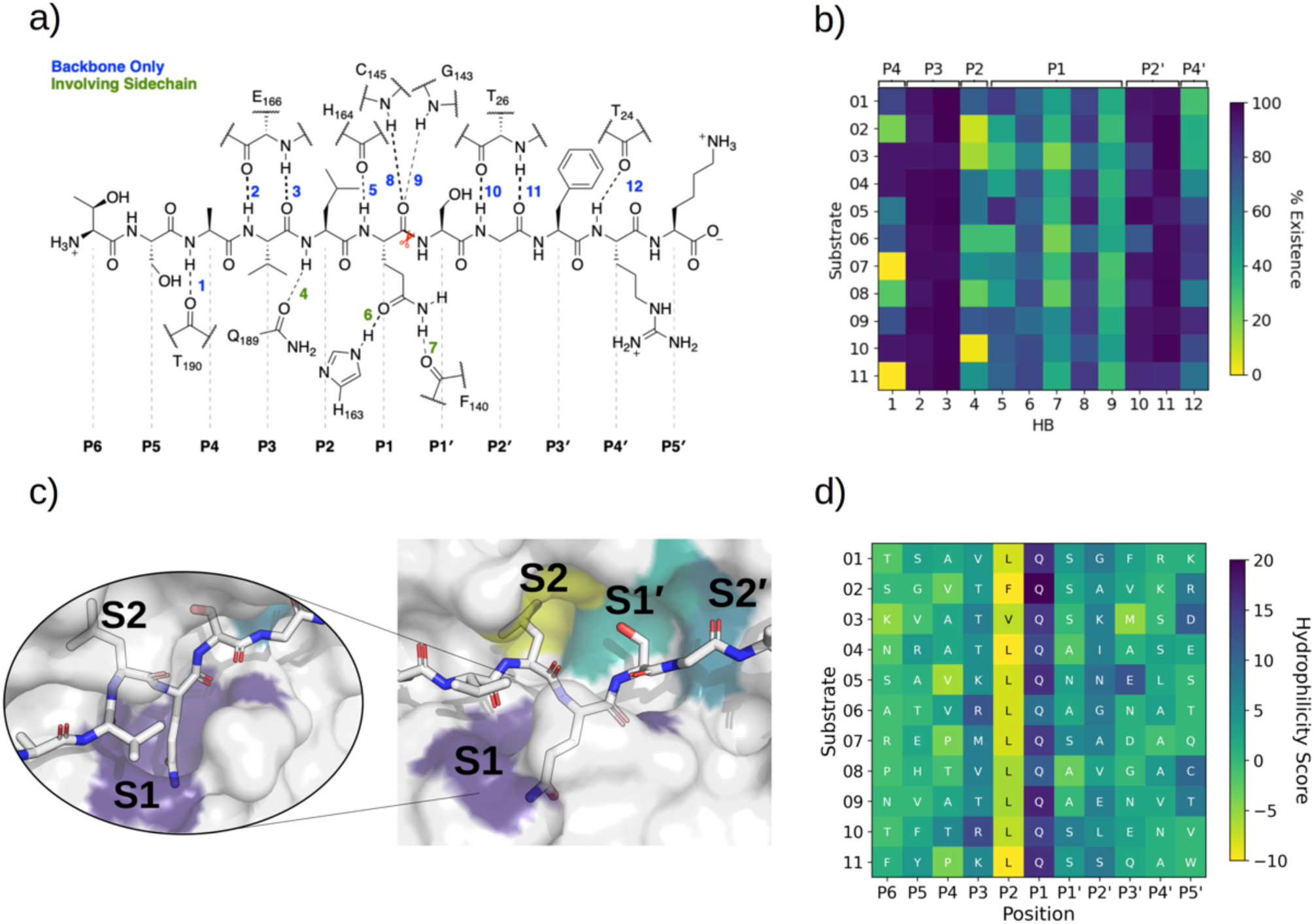
Interactions between SARS-CoV-2 M^pro^ and its substrates. a) M^pro^-substrate hydrogen bonds (HBs), using substrate s01 as an example. b) An annotated heat-map displaying the frequency of each HB observed, with blue indicating highest frequency. Frames were extracted every ns from 600 ns of cumulative explicit-solvent MD conducted per system. c) Close-up of the MD-generated binding mode of SARS-CoV-2 M^pro^-substrate s01 with subsites S1, S2, S1’ and S2’ labelled. Two different angles of the S1 subsite are shown, emphasizing the deep S1 pocket that accommodates the P1 Gln sidechain. Subsite surface colour corresponds to the hydrophilicity score, with hydrophobic subsites shown in yellow, hydrophilic subsites in dark blue, and neutral subsites in turquoise. d) Hydrophilicity map for the 11 substrates calculated as the sum of hydrophilic interactions subtracted from the sum of hydrophobic interactions. Interactions were obtained through analysis of substrate MD poses by Arpeggio^46^ (see **Methods Section S1.5**).

In both the explicit-solvent MD and iMD-VR simulations, all 11 substrates are primarily held in place by stable backbone-backbone HBs (numbered 2, 3, 10 and 11 in **Figure 3**), involving M^pro^ Glu-166 at S3 (2, 3) and Thr-26 at S2’ (10, 11). Both residues consistently form significant contact interactions and energy contributions (*vide infra*). The backbone HBs further away from the cleavage site, *i*.*e*. HBs 1 and 12, show greater variation in explicit water, and HB 1 is completely absent in s07 and s11 where P4 is Pro. By contrast, both of them are well conserved in the iMD-VR-generated structures. In both approaches, HB 4, which involves the flexible Gln-189 side chain, is formed only intermittently.

The P1 Gln is conserved in all 11 M^pro^ cleavage sites (**Figure 1a, b**): while individual HBs formed in the S1 site (HBs 5-9) are observed less frequently than HBs 2, 3, 10 and 11, they outnumber other sites. In explicit-solvent MD simulations, HBs 6 and 8 predominate (**Figure 3b**). In the iMD-VR simulations of s01, s02 and s05, HBs 5, 6, and 7 show variation, while HBs 8 and 9 are well conserved in all three substrates. The consistency in HB 8 formation suggests that this interaction could play a fundamental role in catalysis.

The Cys-145 backbone amide forms part of the oxyanion hole, which stabilises the tetrahedral intermediate formed upon nucleophilic attack of Cys-145 Sγ on the scissile amide carbonyl. M^pro^’s exquisite specificity for Gln at P1 is likely based on the formation of HB 6 with His-163, and to a lesser extent HB 7, along with the narrowness of the S1 pocket accommodating the Gln sidechain in an extended conformation.

#### 2.2.2 Hydrophobicity analysis

To obtain more detailed information on the nature of the interactions in each subsite, we generated a hydrophilicity map (**Figure 3d**). As expected, the S1 subsite has a substantial hydrophilic character consistent throughout all substrates. Although only two highly conserved contact residues were identified between the S2 subsite (Met-49 and Met-165) and all 11 substrates, the character of S2 is nonetheless consistently hydrophobic. In accord with the HB analysis (**Figure 3b**), the conservation of the subsites decreases with increasing distance from the cleavage site. Although subsites S3 and S2’ show a slight bias towards hydrophilic interactions, none of the other subsites show a consistent pattern across the different substrates.

#### 2.2.3 Other contact interactions

Beyond HBs, several other interaction types are conserved across the substrates; these were identified by running Arpeggio^46^ on snapshots extracted from the explicit-solvent MD simulations. As seen in **Figure 4**, 6 out of the 8 most common P1 interactions are conserved across most substrates. This includes the previously described HB pattern between the P1 backbone oxygen and the backbone NHs of Cys-145 (HB 8) and Gly-143 (HB 9) that constitute the oxyanion hole, as well as HB 6 (with His-163) and HB 7 (with Phe-140) that stabilise the P1 Gln sidechain. In addition, interactions with residues Ser-144, His-163, His-164, Glu-166 and, to a lesser extent, Phe-140 were highly conserved at subsite S1. None of the other subsites show this level of consistency in residue-level contacts, although some individual interactions such as hydrophobic interactions at Met-49 and Met-165 were always present in subsite S2. Furthermore, important stabilising backbone HBs (HBs 2-3 between Glu-166 and P3; and HBs 10-11 between Thr-26 and P2’) were conserved with all substrates. Finally, this residue-level contact analysis reveals that the P’ contacts are in general preserved less often than the P side (**Figure 4**). The same trend was found when docking s01, s02, and s05 using iMD-VR, where P’-side residues tended to be more flexible than P-side residues.

**Figure 4:**
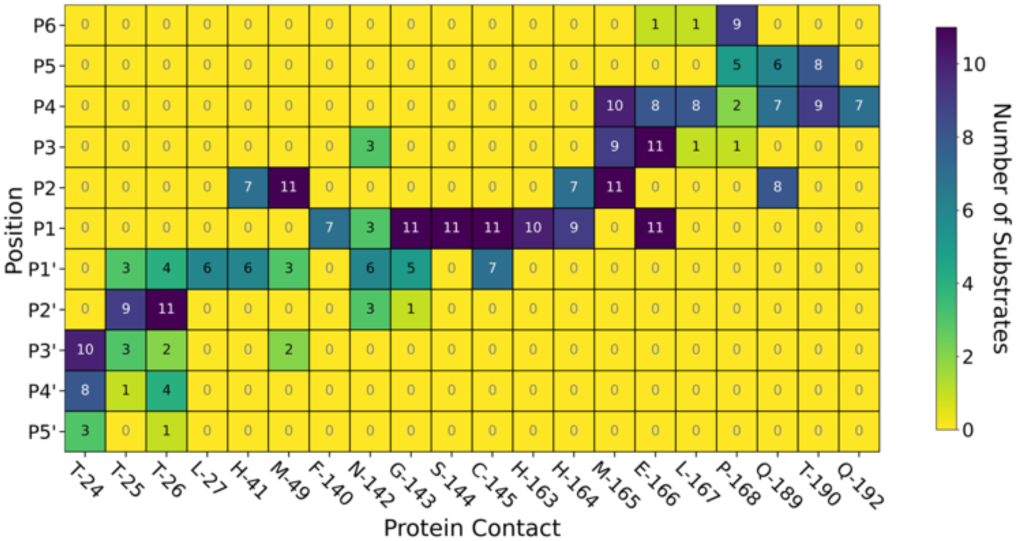
Key M^pro^-substrate contacts. Map of key interactions including HBs and other non-covalent interactions between the 11 substrates and M^pro^ from Arpeggio analysis of the most representative pose for each substrate generated by MD. Dark blue indicates the interaction is observed across all 11 substrates at that substrate position, while yellow indicates no substrates form this interaction.

#### 2.2.4 MM-GBSA analysis

We analysed van der Waals and electrostatic contributions to protein-substrate interactions by employing the molecular mechanics-generalised Born surface area (MM-GBSA) method.^47,48^ This approach has been used for identifying protein hotspot residues that contribute significantly to ligand binding, with a crude estimation of the effects of solvation.^49,50^

The ten M^pro^ residues contributing most to the binding energy were identified for each of the 11 substrate complexes. These hotspot residues were largely conserved across substrates (**Figure S2.12**), with the most frequently-identified residues (and their occurrences across the 11 systems) being: Glu-166 (11), Thr-26 (11), Met-165 (11), His-41 (11), His-163 (10), Cys-145 (10), Pro-168 (9), Thr-25 (8), Gln-189 (8), Thr-24 (6), Thr-190 (6). These residues are all within proximity to the complexed substrate (**Figure 5**) and were also identified by Arpeggio as conserved contacts (**Figure 4**).

**Figure 5:**
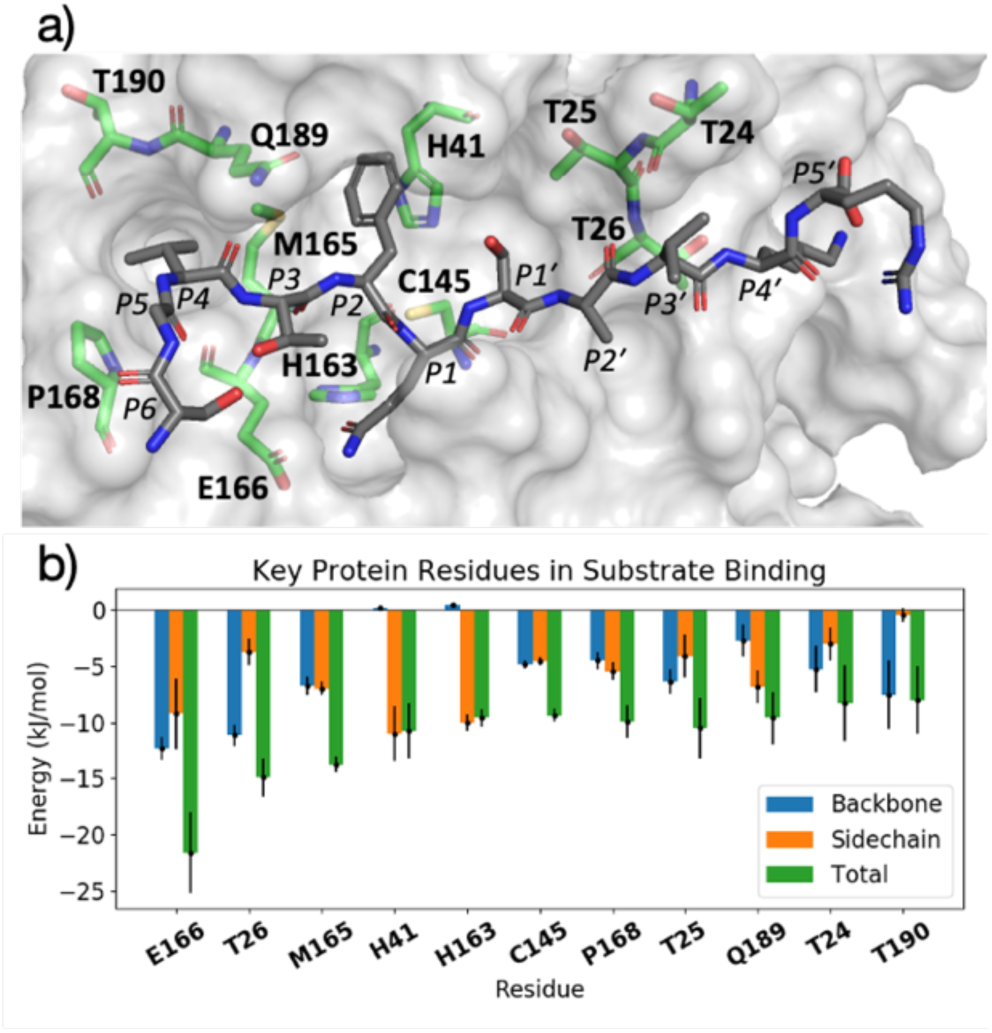
Key M^pro^ residues involved in substrate binding identified by MM-GBSA. a) View of the energy minimised structure of M^pro^ (PDB: 6yb7, light grey surface)^37^ in complex with s02 (dark grey sticks), with hotspot residues as identified by MM-GBSA per-residue decomposition shown (green sticks). b) Contributions to the MM-GBSA binding energy by each hotspot residue (average ± standard deviation across 11 systems), in descending order of consistency across the 11 substrates.

As anticipated, residues that form the most stable HBs (HBs 2, 3, 10, 11), namely Glu-166 and Thr-26, display large favourable contributions (**Figure 5**). Further insight could be gained from decomposition into their respective backbone and sidechain components. Interestingly, while the contributions from their backbones are comparable, as expected from their similar modes of HB formation, Glu-166 has a larger total contribution due to its sidechain, which is adjacent to the conserved P1 Gln sidechain. Other HB-forming residues (backbones of Cys-145, Thr-190, and Thr-24; and sidechains of His-163 and Gln-189) are also identified as hotspots. For the remaining key residues, contributions to binding are dominated by van der Waals interactions (**Figure S2.13**), as exemplified by Met-165 and His-41, both of which engage in non-polar contacts with the hydrophobic P2 residue. Optimising interactions with these hotspot residues could help guide the design of optimal M^pro^ inhibitors.

Similar per-residue decomposition of binding energy contributions was performed at each substrate position. Given the variation in sequences, the contribution from each position varies across substrates (**Figure S2.14**). Nevertheless, binding contributions from the P4, P3, P1 and P2’ backbones are observed with all substrates, in agreement with the observation of HBs between these residues and M^pro^ residues (**Figure 3**).

Contributions from the P2 sidechain are significant (**Figure S2.15**). The S2 site appears to tolerate Phe (s02) well, in addition to the more common Leu that is found in nine of the eleven substrates, while Val (s03) is less favourable. The possibility of filling the S2 pocket with a larger, aromatic moiety is of interest in designing inhibitors.

#### 2.2.5 Density functional theory analysis of the interaction network

We performed linear-scaling DFT calculations on the entire solvated M^pro^-substrate complexes by applying the BigDFT code.^51^ This formalism enables the automatic decomposition of large molecular systems into coarse-grained subsets of atoms, or ‘fragments’, in an unbiased manner.^52^ With such a reduced-complexity description, quantities like inter-fragment bond analysis and interaction strengths can be easily quantified.

Initial comparison between the interaction energies computed from BigDFT and those obtained using classical force fields shows good correlation (**Figure S2.16**), providing further support for the MD results. It also suggests that charge-polarisation, which DFT partially captures but which is absent in force-field based MD, does not play a major role in the studied substrate-enzyme interactions with electrostatic interactions being more important (**Figure S2.17**).

Using representative snapshots extracted from explicitly solvated MD trajectories of M^pro^ complexed with s01, s02, and s05 as inputs for BigDFT calculations, we generated a QM interaction (contact) map to identify interactions between substrates and M^pro^. The strength of the interaction was quantified by the contact interaction energy, *E*_cont_, which is defined as the partial trace of the system’s Hamiltonian between fragments (**Figure 6a**). By computing this quantity between dimeric M^pro^ and s01, with either one or both sites occupied, we were able to evaluate cooperativity effects on binding. The obtained values are within error (*E*_cont_ = −736 ± 119 and −699 ± 95.0 kJ mol^-1^ for one or two substrates bound to the dimer, respectively), suggesting that no significant cooperativity operates between the two active sites in the M^pro^ dimer (at least with investigated substrates). Moreover, no preference for binding in a particular site was observed.

**Figure 6:**
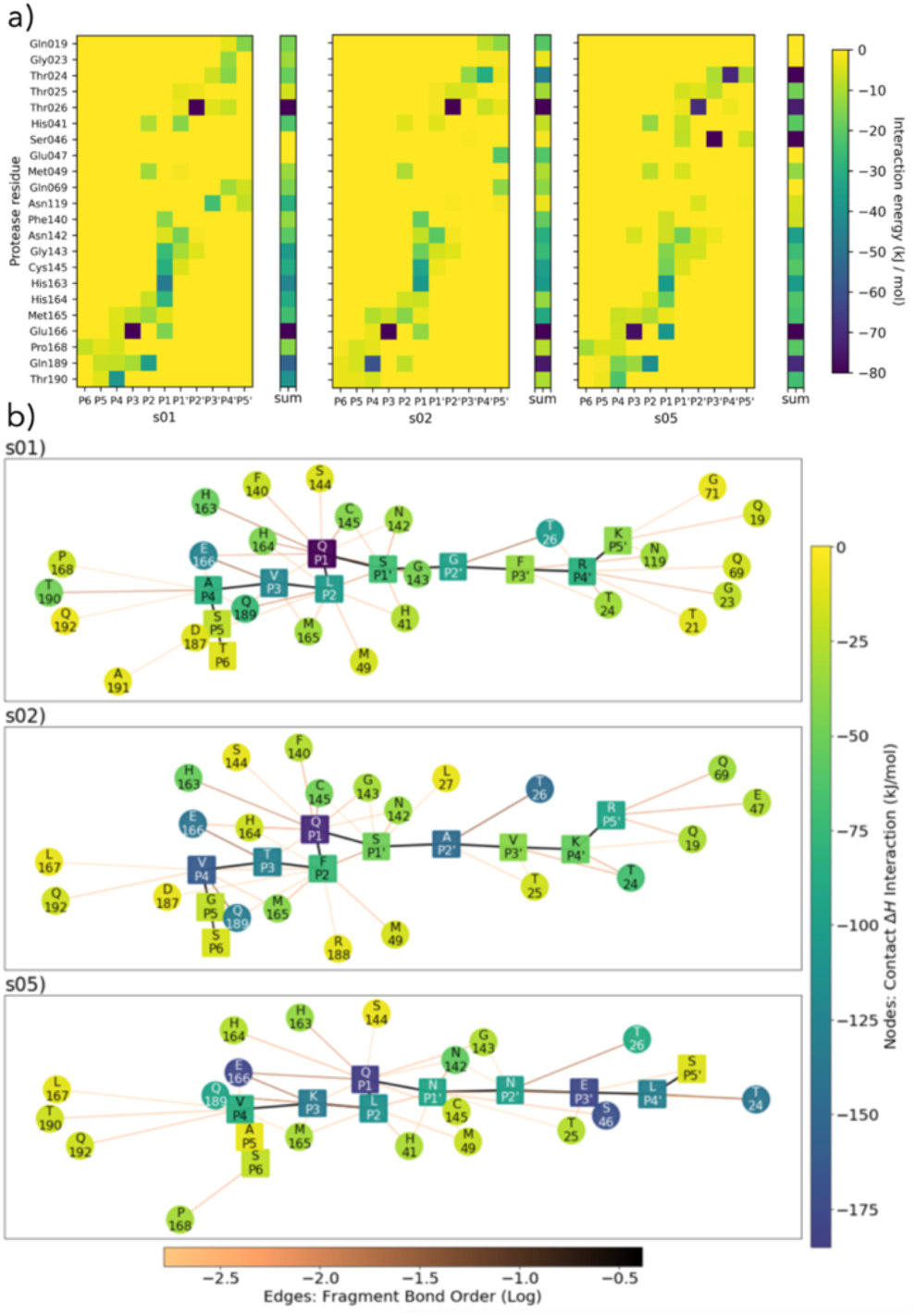
BigDFT analysis of M^pro^-substrate interactions. a) QM contact interaction energies and graphs between 22 selected residues of M^pro^ and s01, s02 and s05. The nature of these interactions can be seen in the M^pro^-substrate HB map in **Figure 3** and 3D representation of key residues for s02 in **Figure 5**. b) Interaction networks where the colour of the nodes refers to the interaction strength, ranging from dark blue for the strongest through green to yellow for the weakest interactions. Square nodes belong to substrates, while circular nodes belong to M^pro^. The width and colour of the edges indicate the fragment bond order between residues, which is a unitless measure associated with bond strength (analogous to an atomic bond order), ranging from black for the strongest to orange for the weakest.^52^ Interaction energies and maps were computed using ensemble-averaged results of MD snapshots using the BigDFT code.

This analysis also recapitulated the key roles of Glu-166 and Thr-26, with interactions observed in all three peptides s01, s02, and s05. This is consistent with the HB analysis described earlier (**Figure 3**). It shows that Gln-189 consistently forms an HB with P2 (HB 4) in s01 and s05, but rarely with s02. The P2 residue in s01 and s05 is Leu, while in s02, P2 is Phe, the bulk of which may weaken the HB. Finally, a strong interaction between Ser-46 and the P3’-Glu sidechain is observed for s05 (P3’=Glu), but not s01 or s02 (P3’=Phe and Val, respectively), suggesting the P3’-Glu carboxylate is important in forming a strong HB with Ser-46.

From this analysis, a graph-like view of substrate-enzyme interactions can be obtained, enabling identification of the most relevant (MD-averaged) interaction networks (**Figure 6b**). Note the interactions are predominantly on the non-prime side of all three substrates.

A conserved contact is present in the three substrates between Cys-145 and both P1 and P1’ residues. Interactions between His-41 and P2/P1’ are observed for s01 and s05, and between Glu-166 and P1/P3 (and, to some extent, to P4) for all three substrates. This analysis singles out the character of s02, which is dominated by the bulky character of its P2 Phe. Substitutions in this region may have a substantial effect on the interaction network close to the catalytic site. While the P side exhibits an interconnected character, the network on the P’ side has a more linear character, once again indicating that hot-spot residues responsible for binding are present on the P side. The relevant interactors on the P side range from P1 to P4, as the interaction graphs show relatively intricate patterns in this region of the substrates. Distributions around the average values are shown in **Figure S2.18**.

The following trends emerge from our studies on models of M^pro^ in complex with its 11 substrates: (i) Binding stability is partly conferred by a series of HBs ranging from the P4 to P4’ positions, in particular those between the backbones of M^pro^ Glu-166 and Thr-26 and substrate positions P3 and P2’ respectively, as well as HBs involving the conserved P1 Gln sidechain; (ii) substrate residues N-terminal of the cleavage site (P side) form a larger number of, and more consistent, contact interactions with M^pro^ compared to the P’ side, with interactions contributed by M^pro^ residues Met-49, Gly-143, Ser-144, Cys-145, His-163, His-164, Met-165 and Glu-166 being the most conserved. We conclude that the S1 and S2 pockets are prime targets for active site substrate-competing inhibitor design due to their well-defined hydrophilic character, large energy contribution to substrate binding and vital conserved hydrogen bonds in S1 for substrate recognition.

#### 2.2.6 Conformational plasticity in M^pro^ crystal structures

While others have compared the dynamics of ligand binding sites across SARS-CoV-2, SARS-CoV and MERS-CoV M^pro^,^53^ we investigated the conformational plasticity of the SARS-CoV-2 M^pro^ active site exhibited in 333 M^pro^:ligand co-crystal structures obtained from Fragalysis^54^ by comparing to a reference *apo* structure of M^pro^ (PDB entry 6yb7),^37^ calculating per-residue heavy atom RMSD between the active site residues of the *apo* and protein-ligand bound structure (**Figure 7**). A high degree of plasticity was observed at residues Thr-24, Thr-25, His-41, Thr-45, Ser-46, Met-49, Asn-142, Met-165, Glu-166, Arg-188, Gln-189 and Ala-191. By contrast, the S1 subsite is particularly rigid, with almost no change in residue conformations across all 333 crystal structures.

**Figure 7:**
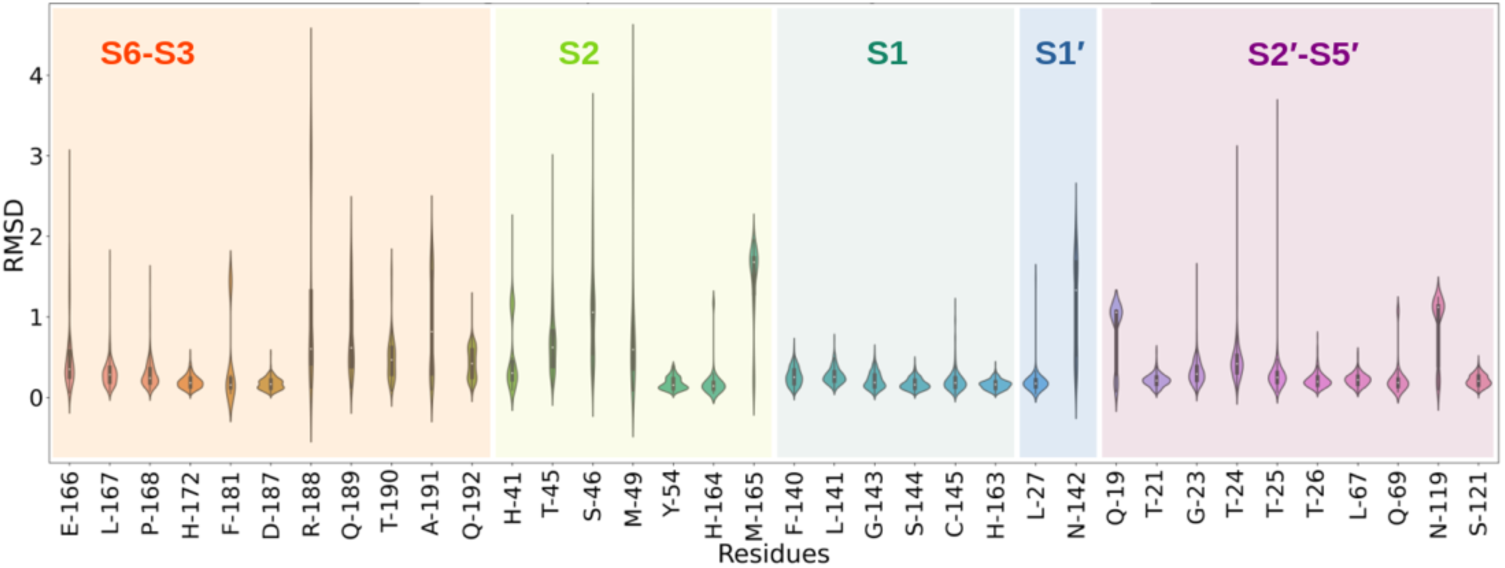
Analysis of the active site plasticity of 333 M^pro^ co-crystal structures. Active site residues (residues 19, 21, 23-27, 41, 45, 46, 49, 54, 67, 69, 119, 121, 140-145, 163-168, 172, 181, 187-192) were chosen based on the MD analysis of the 11 substrate-M^pro^ models and correspond to all M^pro^ residues that contact any substrate. The violin plots show the distributions of the individual heavy atom RMSD values between the 333 M^pro^-ligand co-crystal structures and a reference *apo* structure (PDB 6yb7). Residues corresponding to each subsite are colour-coded.

This is not surprising considering that the P1 Gln is conserved in all 11 substrates. In all cases, Gln recognition is mainly driven by protein-substrate interactions with residues Gly-143, Ser-144, Cys-145 (oxyanion hole) as well as His-163 and Phe-140. On the other hand, the S2 subsite is highly flexible, especially at residues Thr-45, Ser-46 and Met-49. Although the P2 residue in all 11 substrates is conserved in terms of hydrophobicity (Leu, Phe, Val), the S2 pocket is highly flexible and can adapt to accommodate smaller functional groups such as aliphatic carbons in leucine, but also larger substituted aromatic groups found in many of the analysed co-crystal ligands. The outer regions of the active site corresponding to subsites S3-S6 and S2’-S5’ vary in flexibility, echoing our observations from our MD simulations. Active site binding ligands in the 333 co-crystal structures are most likely to bind around S2-S1’ and should only affect the conformation of some of the outer subsite residues. A 3D overlay of all 333 co-crystal structures (**Figure S4.4**) highlights the plasticity at S2.

### 2.3 Monitoring of substrate sequence hydrolysis by mass spectrometry

To rank the SARS-CoV-2 M^pro^ preferences for hydrolysis of the 11 assigned SARS-CoV-2 cleavage sites, we monitored turnover of 11-mer peptides by solid-phase extraction (SPE) coupled to mass spectrometry (MS). Interestingly, after the N-terminal autocleavage site s01, s11 was found to be the next preferred substrate for catalysis (**Figure S2.20**). Peptides s06, s02, and s10 were also hydrolysed, but less efficiently than s11. Slow turnover was observed for s07 and s09. Evidence for low turnover of s05 was obtained after prolonged incubation with M^pro^ (9.56%) (**Figure S2.21**). Under our standard conditions, no evidence for cleavage was observed for s03, s04, and s08.

We then examined turnover under non-denaturing MS conditions using ammonium acetate buffer (**Figure 8a**). Peptides s01, s06, s08, s10 and s11 demonstrated evidence for fast turnover. The level of substrate ion depletion was >70% after 1-min and >90% after 6-min incubation. Peptides s02, s04 and s09 showed substrate ion depletion from 35 to 45% after 1-min incubation, >70% depletion after 6 minutes, and >90% depletion after 12 minutes; while peptides s03, s05 and s07 demonstrated slow turnover with <20% depletion after 1-min incubation and <50% turnover after 6-min incubation that also remained below 50% after 12-min incubation.

**Figure 8:**
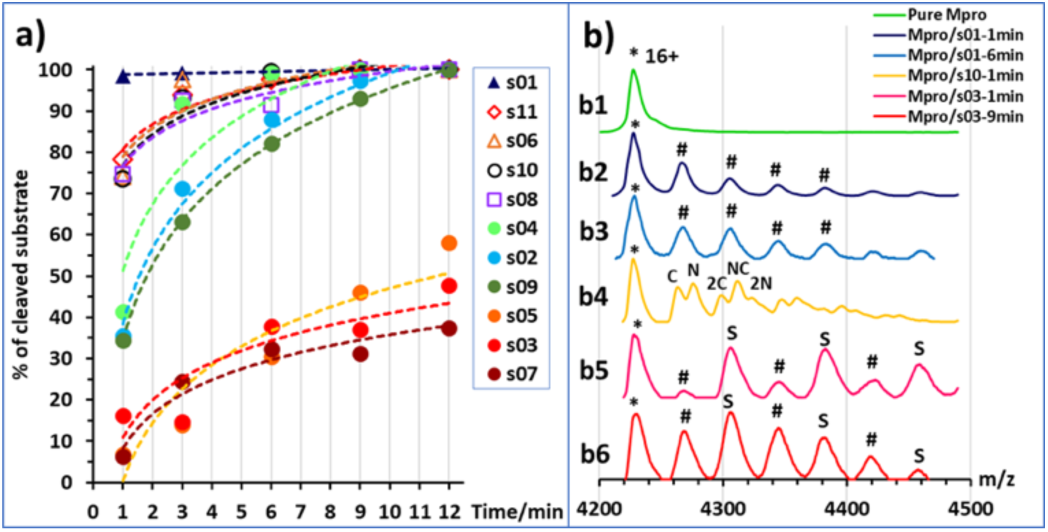
Non-denaturing mass spectrometry of M^pro^ substrate turnover. a) Substrate turnover *versus* incubation time as measured by non-denaturing MS. Trend lines are given for visual guidance only. b) Examples of mass spectra in the *m/z* region around the 16+ charge state of M^pro^ dimer: b1) pure M^pro^ solution (5 μM), asterisk (*) indicates the mass peak of the 16+ charge state of the dimer; b2) M^pro^ and s01 solution after 1-min incubation, hashes (#) indicate the mass peaks corresponding to the s01 cleaved fragments sequentially attached to the M^pro^ dimer; note: the resolution is not sufficient to distinguish between the N- or C-terminal fragments (mass shifts of 617 and 593 Da, respectively); b3) same solution as b2) after 6-min incubation; b4) M^pro^ and s10 solution after 1-min incubation, ‘N’ labels N-terminal fragment(s) attached (765 Da), ‘C’ labels C-terminal fragment(s) attached (560 Da); b5) M^pro^ and s03 solution after 3-min incubation, ‘S’ labels intact substrate(s) attached, hashes (#) label attached substrate fragments, but the N- and C-terminal fragments cannot be distinguished (mass shift 644 and 566 Da, respectively); b6) same solution as b5) after 9-min incubation.

In the protein region of the mass spectra, complexes between M^pro^ dimers and the cleavage products were observed starting from 1 min of incubation for the fast-turnover substrates s01, s06, s08, s10 and s11, and also s02, s04 and s09 (**Figure 8b1-4**). For the slow-turnover substrates s03, s05 and s07, M^pro^ complexes with the intact substrates were observed after 1-min incubation. For longer incubation times, complexes between M^pro^ and the products from these substrates emerged and increased in abundance (**Figure 8b5-6**).

The rank order of the substrates depends on the MS method used, likely due to the differences in the buffers and concentrations used: *i*.*e*., non-denaturing MS used ammonium acetate buffer and an M^pro^ concentration of 5 μM, which is higher than the 0.15 μM used in denaturing MS assays. The higher concentrations of both enzyme and substrate in the non-denaturing MS experiments explain the faster substrate turnover observed in comparison with the denaturing MS method, especially as the concentration of catalytically active M^pro^ dimer would be higher at a higher enzyme concentration.^6,55^

Regardless of the MS method used, a clear trend is observed in the catalytic turnover of the cleavage site-derived peptides. The rank order of substrate preference under denaturing MS conditions was s01 > s11 > s06 > s02 > s10 > s07 > s09 > s05 (Figures S2.20-21). Under non-denaturing conditions (Figure 8) it was: fast turnover (s01, s11, s06, s10, and s08), medium turnover (s04, s02, and s09), and slow turnover (s05, s03, and s07). Substrates s01, s11 and s06 demonstrated the most rapid turnover; while s07, s05 and s03 turnover was slow as measured by both methods. This is in broad agreement with the reversed-phase high performance liquid chromatography analysis of substrate turnover by SARS-CoV M^pro^, where s01 and s02 display fast turnover; s10, s11 and s06 manifest medium turnover; and the rest of the substrates s09, s08, s04, s03, s05, s07 display slow turnover.^43^ Both of our MS studies on SARS-CoV-2 M^pro^ indicate that s02 consistently displayed slower turnover than s11. Previous reports on SARS-CoV M^pro^ have shown evidence for cooperativity between subsites during substrate binding, in particular during autocleavage of the M^pro^ C-terminal site (s02), where the Phe at P2 induces formation of the S3’ subsite to accommodate the P3’ Phe residue.^56^ SARS-CoV-2 M^pro^ substrate s02 has Phe at P2, but not at P3’ (**Figure 1a**). The absence of a Phe at the P3’ position may explain in part the reduced activity of SARS-CoV-2 M^pro^ for s02 relative to s01, compared to the same pair in SARS-CoV M^pro^.^43^

The observed turnover of all 11 SARS-CoV-2 cleavage-site-derived peptides by M^pro^ is consistent with our atomistic models, where the peptides remain bound in the active site during MD simulations and where the scissile amide carbonyl remains well-positioned in the oxyanion hole (*e*.*g*., HB 8 in **Figure 3**) for reaction initiation. The stability of the M^pro^-peptide interactions involving the key S2 and S1 subsites as well as backbone-backbone HBs 2, 3, 10 and 11, could explain the observation in non-denaturing MS of complexes of M^pro^ with products — as a result of slow product dissociation. Nevertheless, we envisage that the order of substrate turnover rates is likely determined by various factors, including peptide conformations, the influence of the P2 and P1’ residues on the catalytic dyad (as highlighted by the BigDFT analysis), entropic effects, and rates of product dissociation, all of which prompt ongoing experimental and computational investigations.

## 3. *In silico* mutational analysis of substrate peptides to inform peptide inhibitor design

Building on insights gained from our binding studies of SARS-CoV-2 M^pro^ and the 11 SARS-CoV-2 polypeptide substrate sequences, we designed peptides that could bind more tightly than the native substrates. We computationally quantified the contribution of each residue of these sequences to the overall binding in the M^pro^ active site and suggested substitutes at each position that would increase affinity. We hypothesised that these peptides would: a) behave as competitive inhibitors, and b) provide counterpoints for comparison with natural substrates, shedding light on the requirements for M^pro^ binding and, perhaps, turnover.

### 3.1 *In silico* alanine scanning and saturation mutagenesis

We used the interactive web application BAlaS to perform computational alanine-scanning mutagenesis (CAS) using BudeAlaScan^57^ and the BUDE_SM algorithm.^58^ Both are built on the docking algorithm BUDE,^59^ which uses a semi-empirical free energy force-field to calculate binding energies.^60^ To identify key interactions determining the binding of the natural substrate peptides to M^pro^, the 11 substrates in complex with M^pro^ were first subjected to CAS using BAlaS. By sequentially substituting for alanine, the energetic contribution of each residue to the overall interaction energy between the singly-mutated peptide and M^pro^ is calculated using:

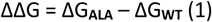

where ΔG_WT_ is the interaction energy between the peptide and M^pro^, and ΔG_ALA_ is the interaction energy for the peptide with a single alanine mutation at a given position. The more positive the value for each residue, the greater the contribution from the substrate residue to binding. This method was used later to evaluate potential inhibitor peptides.

Having identified residues contributing most to the binding energy of the natural M^pro^ substrates, each of the sequences was subjected to saturation mutagenesis using the BUDE_SM algorithm. This algorithm sequentially substitutes each residue with a range of residues (D, E, F, H, I, K, L, M, N, Q, R, S, T, V, W and Y) and calculates the ΔΔG = ΔG_**WT**_ – ΔG_**MUT**_ for the binding interaction of each, entire, singly mutated peptide with M^pro^. Substitutions predicted to improve binding over wildtype sequences have a positive ΔΔG. **Figure 9** shows an example of the BUDE_SM saturation mutagenesis results for all the P2 substitutions for the 11 substrate peptides (normally Leu, Phe, or Val in the 11 substrates). The most positive results suggest that Phe, Trp and Tyr favour increased predicted affinity at P2 (**Figure 9b**). However, although Tyr generally increased the predicted binding affinity (ΔΔG_SUM_ = 68.8 kJ mol^-1^), it was not considered for substitution at P2 due to its negative effect at this position in s11 (scoring −18.9, **Figure 9a**). Candidate residues for each position, P6 - P5’, were shortlisted similarly based on those with the best total, and the fewest unfavourable, scores (**Figure 9a**).

**Figure 9:**
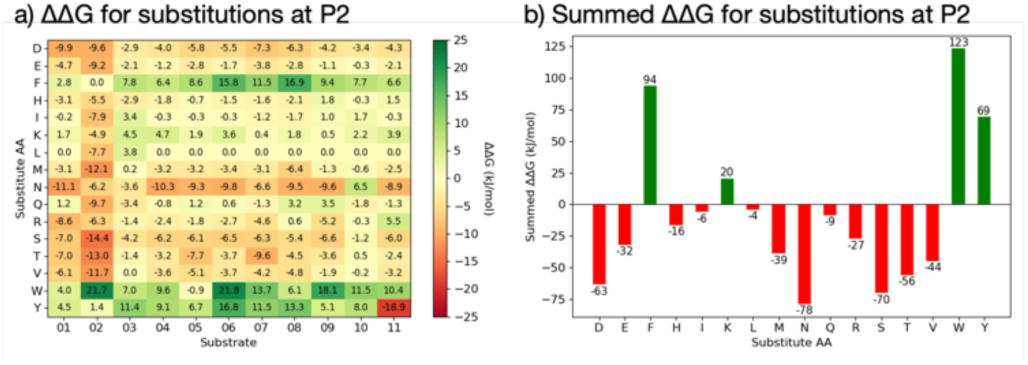
BUDE_SM saturation mutagenesis at the P2 position. a) Heat-map for BUDE_SM saturation mutagenesis at the P2 position for all the M^pro^ substrate sequences showing the ΔΔG = ΔG_WT_ – ΔG_MUT_ value calculated for each substitution. Mutations predicted to improve peptide binding have a positive ΔΔG and are greener; those disfavouring binding are in red. b) The summed ΔΔG values (ΔΔG_SUM_) for each of the residues substituted at P2 of the substrates.

In addition to the computed ΔΔG values for the substrates, the contribution each residue makes to promote an extended conformation was considered. All of s01-s11 adopt a largely extended conformation on binding; it was, therefore, decided that entropic penalties may be avoided if an inherently extended conformation could be favoured in the design of any potentially inhibiting peptide. Thus, the best β-forming (and therefore least α-forming) residues from the first triage were selected (**Figure 10a**).^61^ Another important consideration was solubility. This was achieved by limiting the number of hydrophobic residues in each designed peptide and ensuring a net positive charge (with the exception of p14, where we tested a neutral peptide).

**Figure 10:**
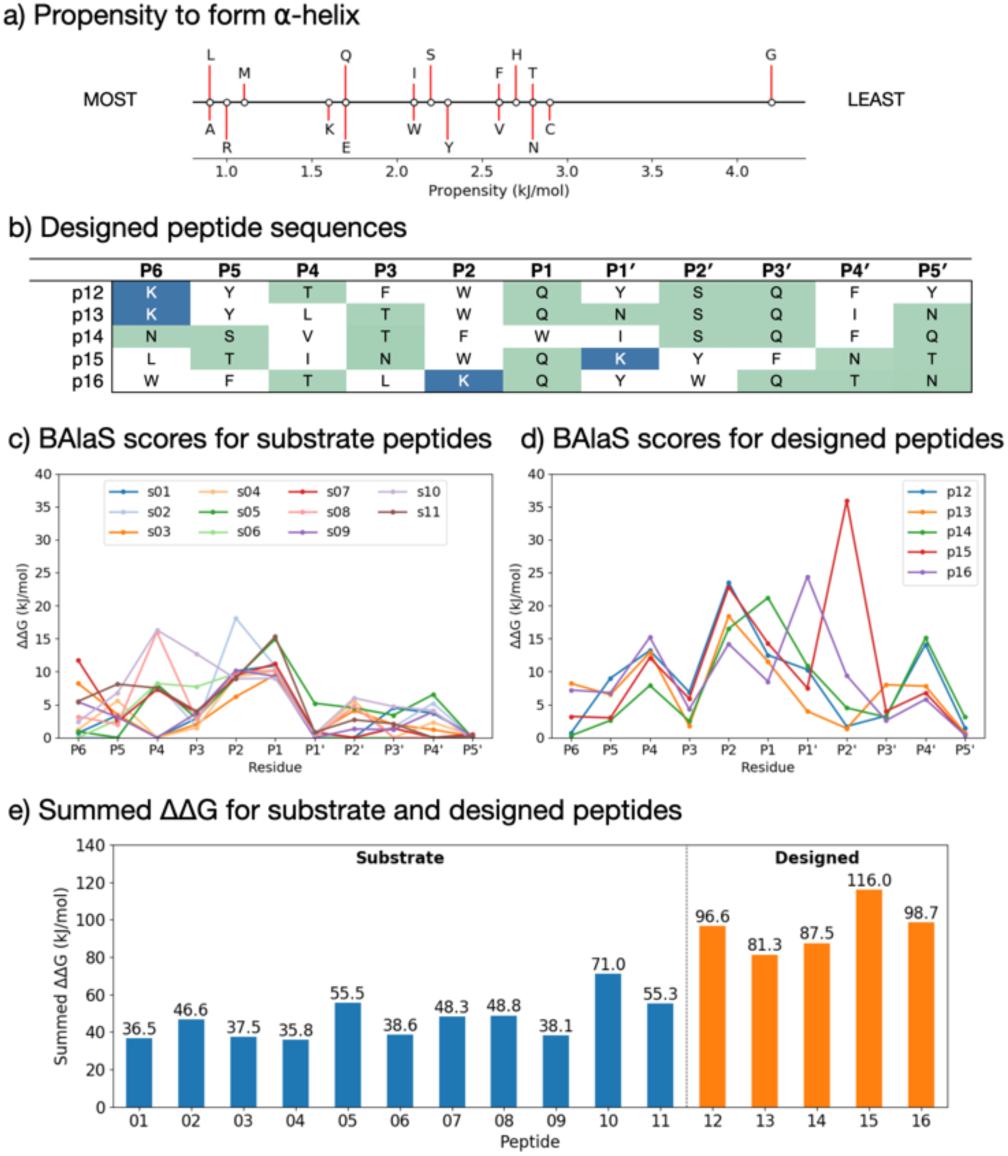
BAlaS-guided design of tight-binding peptides. a) Propensity scale of each amino acid to form an α-helical peptide conformation. b) Sequences of designed peptides p12-p16. Scatter plots with predicted BAlaS ΔΔG = ΔG_ALA_ – ΔG_WT_ values on substitution to alanine for each residue of c) the 11 M^pro^ natural substrates and d) peptides designed based on these. The more positive the value, the greater the contribution made by the sidechain to the overall binding energy. e) The BAlaS ΔΔG_SUM_ comparing values between complexes of M^pro^ with substrate and designed peptides as a proxy for predicting relative binding affinity (larger score = tighter binder).

### 3.2 Designed peptide sequences

Employing the criteria described above, five new peptides, p12-p16, were designed (**Figure 10b**). Comparison of the computed ΔΔG values for s01-s11 (**Figure 10c**) and p12-p16 (**Figure 10d**) reveals that substitutions at the P sites provide only occasional, moderate improvements to binding energy over the corresponding substrate P sites, with the notable exception of P2, which can accommodate Trp, Phe or Lys. These results are in line with the HB analysis which predicts that the sidechains of residues that are on the N-terminal side of the cleavage site (P sites) contribute more to binding than those C-terminal, at the P’ sites. The most striking difference between substrate and designed peptides is in this P’ region, where the predicted binding energy contributions for the designed peptides exceed those of the substrates; an advantage that is distributed over most of the designed P’ positions.

The final step in design was to assess the relative binding affinities of the substrates and designed peptides. Hence the summed ΔΔGs (**Figure 10e**) provide a proxy for the binding energies (BAlaS)^62^ for the substrates and designed peptides with M^pro^. The substrate/M^pro^ complexes are stabilised by an average of 46.5 kJ mol^-1^, whereas the designed-peptide/M^pro^ complexes are predicted to have, in some cases, double the interaction stability of the substrates, with an average of 96.0 kJ mol^-1^. The full analysis is in the **SI file** SI_BAlaS_BUDE_SM_07-06-2021.xls.

### 3.3 Experimental verification of the designed peptides

To test the designed sequences, p12, p13, p15 and p16 were synthesised with a carboxyl-amide terminus by automated solid phase synthesis. Their M^pro^ inhibitory activity was determined by dose-response analysis (**Table 1**) using SPE MS, monitoring both substrate s01 (1191.68 Da) depletion and N-terminally cleaved product (617 Da) formation. Ebselen which reacts multiple times with M^pro 63^ was used as a standard (IC_50_ = 0.14 ± 0.04 µM) (**Figure S3.1f**).

**Table 1:** Designed peptides inhibit M^pro^ in a dose-dependent manner. The assay conditions were 0.15 µM M^pro^, 2 µM s01 in 20 mM HEPES, pH 7.5, and 50 mM NaCl.

| Peptide | $IC_{50} / \mu\text{M}$ | Hill Slope |
| --- | --- | --- |
| p12 | $5.36 \pm 2.17$ | $1.25 \pm 0.06$ |
| p13 | $3.11 \pm 1.80$ | $0.94 \pm 0.09$ |
| p15 | $5.31 \pm 1.08$ | $1.17 \pm 0.16$ |
| p16 | $3.76 \pm 0.51$ | $1.19 \pm 0.16$ |

All four designed peptides manifested similar potency with IC_50_ values ranging from 3.11 µM to 5.36 µM (**Figure S3.1**). Strikingly, despite the presence of Gln at P1 in all the designed peptides that were assayed, no evidence for hydrolysis was observed by SPE MS. This observation was supported by LCMS studies of the peptides incubated overnight with M^pro^ (**Figure S3.3**). We probed the inhibition mode of the designed peptides by monitoring changes in IC_50_ while varying the substrate concentration. Dose-response curve analysis performed with 2 µM, 10 µM, 20 µM and 40 µM TSAVLQ↓SGFRK-NH_2_ s01 (*K*_m_ ∼14.4 µM)^63^ indicated a linear dependency between substrate concentration and calculated IC_50_ values (**Figure 11a-d**). This was not observed with a control 15-mer peptide or ebselen (**Figure 11e, f**). Analysis of the data using the procedure of Wei *et al*.^64^ implies competitive inhibition (**Figure S3.4** and **Tables S3.1-2**). By contrast, the same analysis for ebselen did not support competitive inhibition, consistent with MS studies showing it has a complex mode of inhibition.^63^

**Figure 11:**
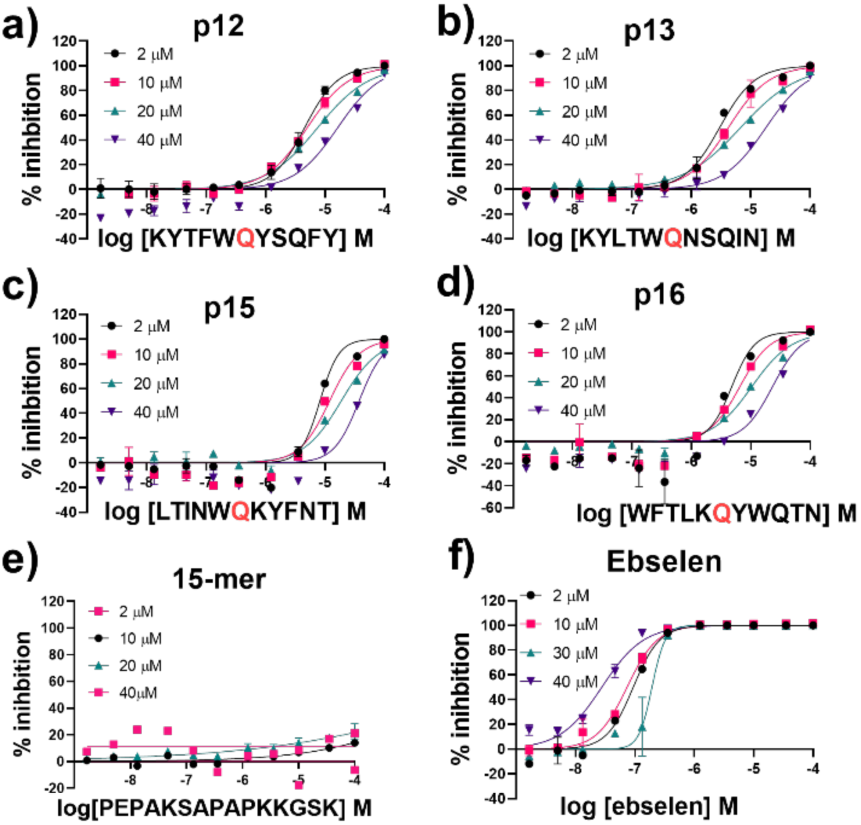
IC_50_ of designed peptides against M^pro^ with varied substrate concentrations. IC_50_s for a) p12, b) p13, c) p15, d) p16, e) 15-mer control peptide and f) ebselen with 2 µM, 10 µM, 20 µM and 40 µM of substrate peptide s01. IC_50_ values were calculated from means of technical duplicates and are provided in **Table S3.2**. See Experimental **Section S1.8** for assay details.

Three of the synthesised peptides — p12, p13, and p15 — have a Trp at P2 (**Figure 10b)** while the other, p16, has a Lys at P2. The 11 M^pro^ substrates all have hydrophobic residues (Leu, Val or Phe) at P2 (**Figure 1a**). To investigate if the nature of the hydrophobic residue at P2, or the hydrophilic nature of the Gln at P1, alters the interaction of the peptide and hence its catalysis in the active site, we synthesised p13-WP2L, s01-LP2W, and s01-QP1W. There was no evidence for cleavage of p13-WP2L or s01-QP1W. Only s01-LP2W underwent partial cleavage (12.6 ± 4.5) % after overnight incubation. These results suggest that the presence of a Trp at P2 hinders catalytically productive binding, at least with these peptides, and that other residues (including the P1’ and P2’ residues) play a role in orienting the substrates for cleavage (*vide infra*).

We then used non-denaturing protein MS to study enzyme-substrate/product/inhibitor complexes simultaneously with turnover. Complexes between M^pro^ dimer and p12 and p13 were observed, together with the uncomplexed M^pro^ dimer in the protein region of the mass spectra. No binding was observed for p15 and p16, due to relatively high noise in that *m/z* region. None of the designed peptides were cleaved by M^pro^, as recorded in the peptide region. As a control, s01 was added to the protein/inhibitor mixtures; for all the inhibitors, turnover of s01 was observed after 3-min incubation. Depletion of s01 was 95%, 91%, 70% and 78% in the presence of p12, p13, p15 and p16, respectively, with an 8-fold excess of inhibitor over M^pro^, *versus* >98% depletion for the M^pro^/s01 mixture without the inhibitor. In the protein region of the mass spectra, complexes between M^pro^ dimers and the s01-cleavage products were observed in the presence of p13, but the abundance of these complexes was lower than the abundance of M^pro^/p13 complexes (**Figure 12**). These results validate the above-described evidence that the peptide inhibitors both bind and competitively inhibit M^pro^.

**Figure 12:**
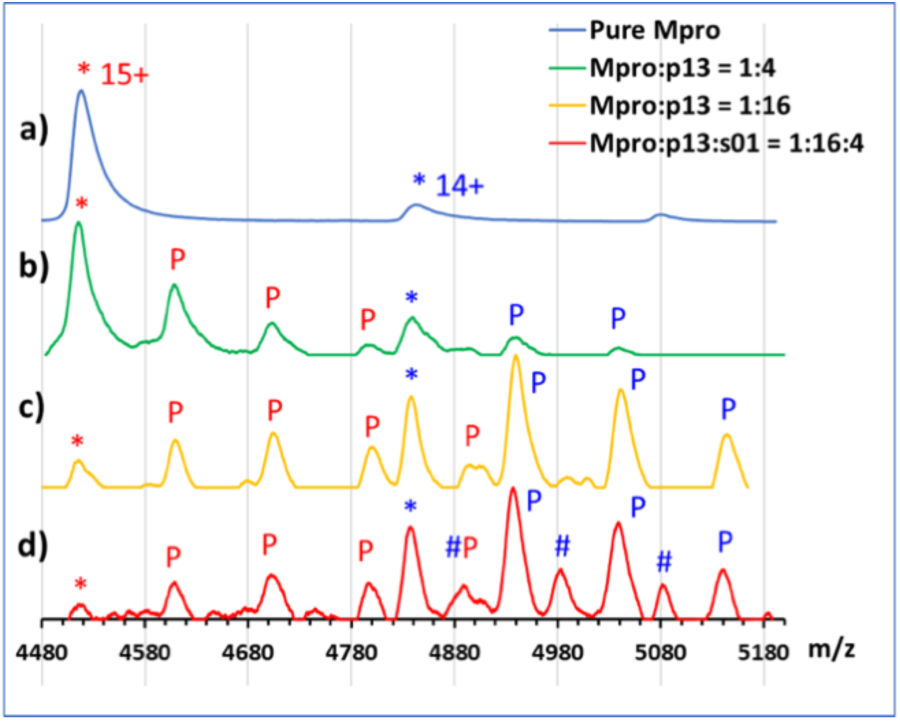
Non-denaturing MS analysis of designed peptides binding to the M^pro^ dimer. Inhibitor binding from non-denaturing MS in the *m/z* region around the 14+ and 15+ charge states of the M^pro^ dimer. (*) indicates unbound M^pro^ dimer peaks. a) 5 μM M^pro^ solution; b) 4-fold excess of p13 relative to the M^pro^ dimer; ‘P’ indicates sequential binding of p13 peptides to M^pro^ in the 15+ charge state (red) and 14+ state (blue); c) 16-fold excess of p13; d) 16-fold excess of p13 and 4-fold excess of s01; hash (#) indicates sequential binding of s01-cleavage products (note: the resolution is not sufficient to distinguish between the N- and C-terminal fragments; some non-specific binding of p13 is also observed in c) and d) due to the high concentration of the peptide).

### 3.4 Understanding the basis of SARS-CoV-2 M^pro^ inhibition by the designed peptides

To investigate binding of the designed peptides, models of dimeric SARS-CoV-2 M^pro^ complexed with p12 (KYTFWQ↓YSQFY) and p13 (KYLTWQ↓NSQIN) were constructed using a high-resolution structure (PDB entry 6yb7).^37^ Modelling of the peptides was carried out using conventional explicitly-solvated MD, iMD-VR, and linear-scaling DFT. For iMD-VR, each simulation was repeated five times, with both the dimeric M^pro^ and the peptide treated as fully flexible. For each peptide, each of the five structures created by iMD-VR was minimised, equilibrated for 4 ns, and subjected to 500 ns of MD. Separately, comparative modelling was also performed: for this, sidechain substitutions were performed to convert the earlier modelled M^pro^-bound s02 peptide into the designed sequences (**Figure S3.5**), before conducting three independent 200 ns MD simulations for each complex in explicit solvent.

#### 3.4.1 Explicitly-solvated MD and implicit solvent iMD-VR

Regardless of the methods employed, p12 and p13 were observed to bind stably at the active site over the course of simulations (**Figures S3.6-9**). Similar to the trend with the substrate complexes, the backbone-backbone HBs involving Glu-166, Thr-26, Thr-24, and the oxyanion hole-contributing Cys-145 were all formed and maintained (**Figures S3.10-12**). Conversely, HBs involving the P1-Gln sidechain of p12 and p13 showed greater variability.

The favourability of the P2 Trp, as predicted by the BAlaS scores, together with the plasticity observed at the S2 site in our structural analysis, prompted us to investigate its binding. Across the simulations, the Trp sidechain was observed to adopt a variety of conformations with varying degrees of immersion in S2 (**Figures 13 and S3.13-14**). In the case of the iMD-VR docked structures, the S2 pocket seemed to change relative to the crystalographically observed conformation, leaving more space in the pocket, resulting in many P2 Trp conformations. This could be due to disturbance of the S2 atoms in iMD-VR simulations, or due to innate plasticity of the S2 site (see **Section 2.2**). An additional iMD-VR docking simulation was performed for p12 and p13, where the focus was on not deforming the S2 pocket. The resulting docked structures of both p12 and p13 were subjected to three independent 100 ns implicit solvent MD simulations; representative P2 Trp conformations in each simulation are in **Figure 13**.

**Figure 13:**
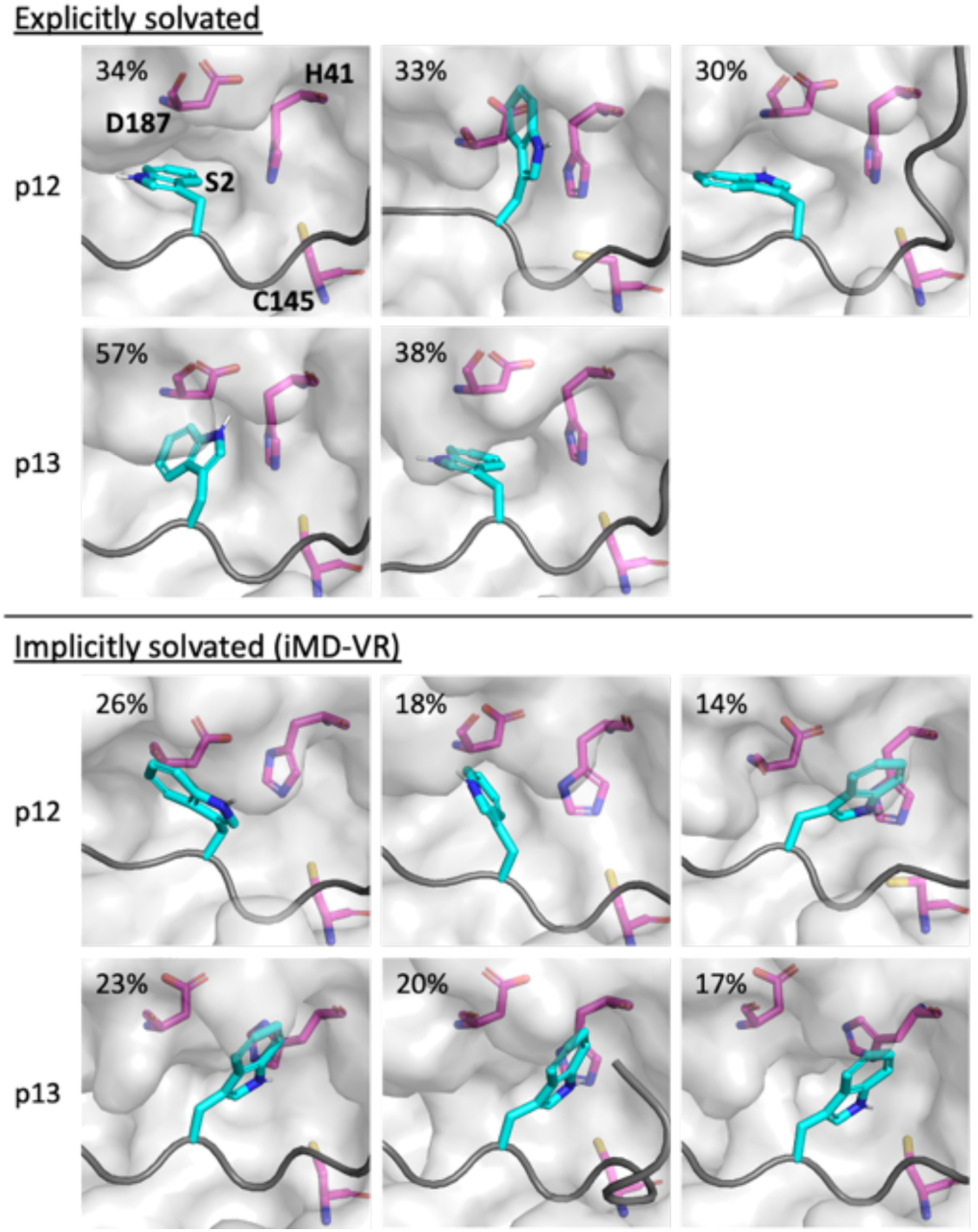
Binding of the P2 Trp residue in the designed peptides. Conformations adopted by the P2 Trp sidechain (cyan sticks; non-polar hydrogens omitted for clarity) in p12 and p13 (grey ribbon) observed during explicit and implicit solvent MD simulations, displayed using representative structures obtained by RMSD clustering. His-41, Cys-145 and Asp-187 are shown as magenta sticks. See **Figures S3.13-14** for cluster populations over the course of the MD. For each peptide, structures are displayed in descending order of occurrence of their respective clusters (above 10%).

Analysis of the poses of the highest populated cluster from MD using Arpeggio-generated hydrophilicity maps (**Figure S3.15**) reveals that the P2 Trp can become more deeply buried within S2 than the analogous P2 residues in the natural substrates, forming more than double the number of hydrophobic contacts in the cases of p12 and p13. Some conformations involve the indole group π-π-stacking or hydrogen-bonding (via the indole N-H) with the catalytic His-41 sidechain forming an extended HB network (**Figure 13**). It is likely these interactions will affect the reactivity of His-41 in deprotonating Cys-145 during the initiation step of peptide hydrolysis.

For comparison, we performed the same DFT-based interaction network analysis as for the substrates (**Figure 6b**) with p13 (**Figure 14**). The interaction graphs near the cleavage site are similar to those for s05, with the P2 Trp interacting with Asp-187. The interaction with His-41, which was not visible in the case of s02 (P2=Phe), is now restored as in the P2 Leu of the other substrates. This analysis suggests that s05 and p13 exhibit the same short-range interaction network.

**Figure 14:**
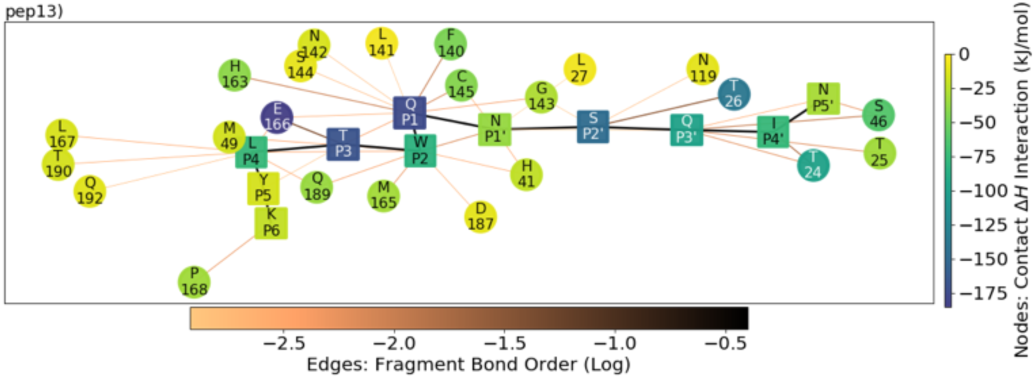
QM contact interaction graph for p13 and M^pro^. Interactions are computed using ensemble-averaged results of MD snapshots with the BigDFT code.^51^

#### 3.4.2 Comparative peptide docking

To investigate the ability of the M^pro^ subsites to recognise residues in the designed sequences, AutoDock CrankPep (ADCP) was used.^65^ A trial was performed by redocking the s01 sequence into the H41A SARS-CoV M^pro^ structure originally complexed with s01 (PDB entry 2q6g).^35^ ADCP successfully positioned the peptide mostly correctly in its top solution, with the C_a_ positions from P5 to P1’ deviating by less than 1 Å (**Table S3.4** and **Figure S3.16**). Deviations increased up to 16 Å at P5’ as the peptide coiled up in the P’ positions, but this is deemed acceptable since the S’ subsites are less well defined, as discussed in the earlier sections.

Following the promising redocking results with ADCP, s01-s11, p12, p13, p15, and p16 were docked with an M^pro^ structure originally complexed with the N3 inhibitor (PDB entry 7bqy; 1.70 Å resolution).^2^ In the case of the substrates, docked structures having the P4 and P2 residues correctly positioned in their corresponding S4 and S2 pockets were consistently found in at least one of the top 10 solutions (**Table S3.5**). From P1 onwards greater variation was observed, with some peptide backbones falling through the S1 pocket rather than adopting an extended conformation, likely due to the less well-defined S’ subsites (**Figure S3.17**). For the designed peptides, by contrast, docking appeared less successful (except p16), with none of the top 10 solutions positioning the peptide in the manner observed in our MD simulations (**Figure S3.18**). We anticipated that the existing S2 pocket in the structure, which accommodates the Leu sidechain of N3, might be too shallow, given the assumption of a rigid receptor in ADCP docking, to accommodate the larger Trp side chain. Hence, the four inhibitor peptides were also docked to the C145A M^pro^ structure in complex with the s02 cleaved product (PDB entry 7joy; 2 Å resolution),^66^ which has a deeper S2 pocket adapted to the P2 Phe sidechain in s02. Interestingly, for both p12 and p16, the top docked solution matched our design more closely (**Table S3.6** and **Figure S3.19**). Docking of p13 and p15 remained challenging, possibly due to the difficulty of recognising a larger Leu (p13) or Ile (p15) residue in the S4 pocket, which originally accommodates a Val side chain. This highlights the potential of M^pro^ active site pockets to adapt their shapes when binding to different substrates or inhibitors.

In summary, we used *in silico* saturation mutagenesis to design peptide sequences that were shown *in vitro* to inhibit M^pro^ competitively. Structures of p12 and p13 generated by both iMD-VR docking and comparative modelling were similar in terms of HB formation and peptide backbone RMSD and RMSF on performing MD. These studies demonstrate the ability of the S2 subsite to adapt its pocket size and interaction network via induced fit to accommodate different substrate or inhibitor P2 residues.

While these models suggested similarly stable binding modes as seen with the natural substrates, turnover of these inhibitor peptides by M^pro^ was not detected. This may be due to the more favourable binding affinity predicted by BUDE, in terms of both the higher overall interaction energies and the increased contribution by the P’ residues, in comparison with the natural substrates. This may as a consequence increase the stability of the designed peptide complexes. It is also possible that the larger P2 residue interferes with the catalytic His-41 residue, given their proximity seen in MD simulations (**Figure 13**).

## 4. Analysis of results from high throughput crystallography with fragments

Having identified key interactions by which M^pro^ recognises its substrates, we hypothesised that this information might be reflected in the extensive small-molecule inhibitor work on M^pro^ and could, in turn, be exploited for the design of novel small-molecule inhibitors. In particular, we explored whether ligands that adopt the same key contacts as the natural substrates could lead to better inhibitory activity. To investigate this, we utilised all 91 X-ray structures of small molecule fragments complexed with M^pro^ obtained by high-throughput crystallographic screening at Diamond’s XChem facility^67^ as well as the dataset of 798 inhibitors and 245 crystal structures obtained from the COVID Moonshot project^33^ and analysed them by their protein-ligand interaction patterns.

### 4.1 Interaction analysis of the XChem fragments

Our analysis divided fragment binding into non-active-site binders (25 fragments) and active-site binding / likely substrate competing (66 fragments) molecules (**Figure S4.1**). A fingerprint bit-vector was constructed for every active-site binding fragment, with each bit denoting the presence or absence of an interaction with every M^pro^ protein residue found to interact with either a substrate or fragment. A distance matrix was created by calculating the Tanimoto distance^68^ between interaction fingerprints. Fragments were then clustered using the interaction Tanimoto similarity index. A similarity score of 1 corresponds to perfect overlap of all fragment-residue level contacts, and 0 denotes no overlap. Note that the Tanimoto index is computed between Arpeggio-derived contact fingerprints rather than ligand structure extended-connectivity fingerprints (ECFPs). For this analysis, a tighter (0.7) and broader (0.5) threshold were chosen. The former was found to be better at distinguishing distinct binding modes, while the broader threshold was more useful in grouping binders around major significant interactions (see **Table S4.1** and **Figures S4.2-3** for further details). We focus on the results obtained from the broader 0.5 threshold clustering (**Figure 15**).

**Figure 15:**
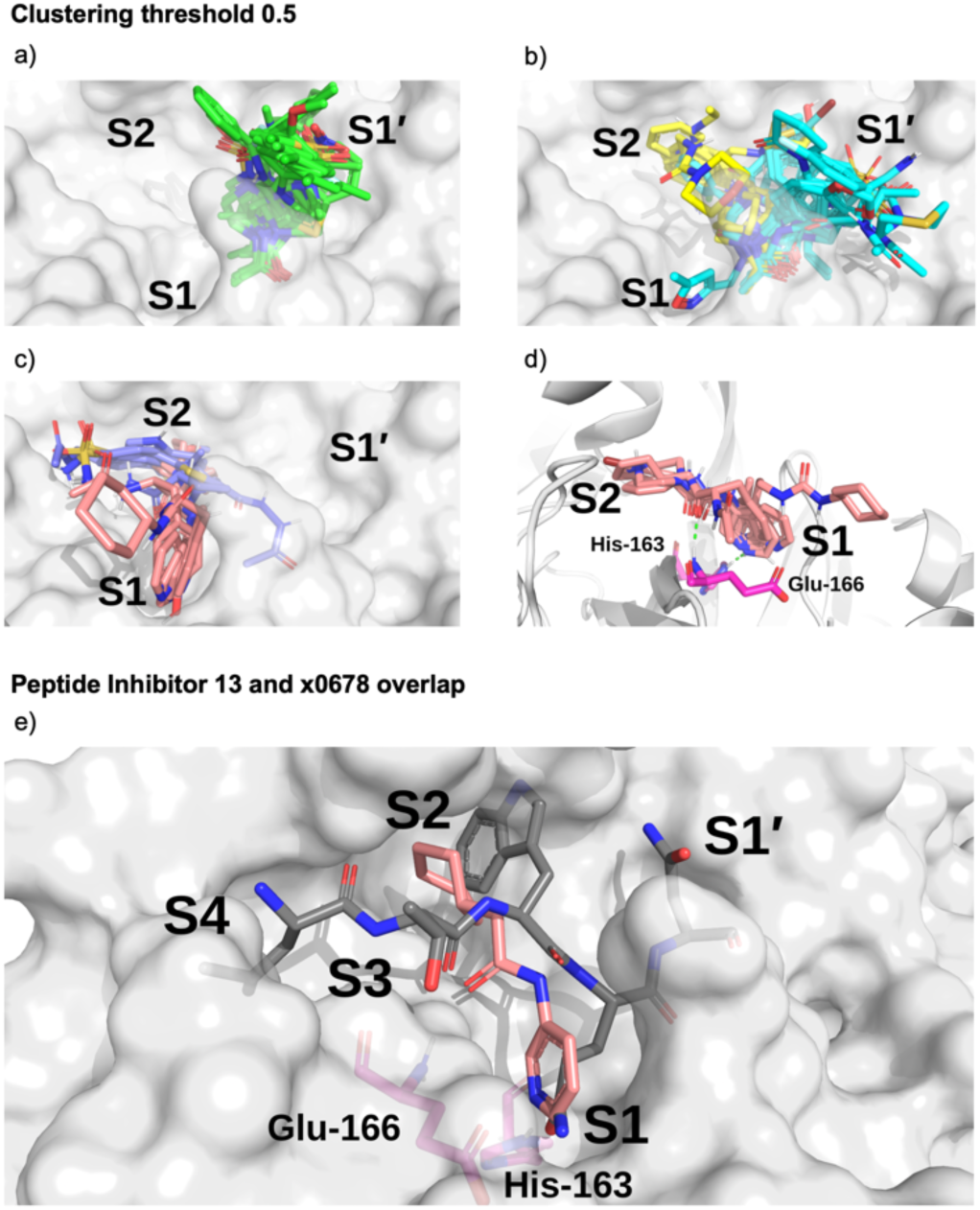
Clustering of XChem active site-binding fragments. Surface of the x0830-bound M^pro^ structure (in white) and the top 5 most populated fragment clusters using a clustering threshold of 0.5. a) Cluster 1 fragments tend to occupy S1’ (green); b) clusters 2 (cyan) and 3 (yellow) tend to span S1’ and S2; c) clusters 4 (lilac) and 5 (salmon) tend to occupy S2 and S1. d) Close-up on the binding pose of cluster 5. Shown in green dotted lines are the two key HBs between the fragment carbonyl oxygen and the backbone nitrogen of Glu-166 (HB 3 in **Figure 3**) and between the His-163 N*ε* and the heterocyclic nitrogen of the fragment (HB 6 in **Figure 3**). e) Overlay of the P4-P1’-truncated structure of peptide inhibitor p13 from an MD snapshot and cluster 5 binder x0678, on the corresponding x0678 co-crystal structure molecular surface.

All the fragments and ligands belonging to clusters 1-2 (except x0397, x0978 and x0981) are covalently bound to Cys-145. As a result, a highly-conserved binding mode is observed for the carbonyl-containing covalent warheads (*e*.*g*., chloroacetamides), where the carbonyl oxygen close to the covalent warhead binds into the oxyanion hole between residues Gly-143 and Cys-145, mimicking substrate HBs 8 and 9 identified in **Section 2.2.1** (**Figure 3**). Among the other clusters, cluster 5 stands out as it the only major cluster with fragments that bind deeply into the S1 pocket, having one of the main conserved contacts identified for the substrate peptides. This cluster shows a distinct binding motif primarily driven by (i) hydrogen bonding between a carbonyl oxygen on the fragment and the Glu-166 backbone NH-group, and (ii) a strong polar interaction between His-163 and the fragment. Notably, the position of the hydrogen on the imidazole sidechain of His-163 appears to depend on the HB fragment partner.

Based on the presence or absence of either a HB donor or acceptor on the fragment, the protonation state of His-163 can be inferred. This suggests that for x0107, x0434, x0540, x0678 and x0967, the His-163 *ε*-nitrogen is protonated, forming a HB to the pyridine nitrogen (x0107, x0434, x0540 and x0678) or phenol oxygen (x0967). For x1093, the *δ*-nitrogen is protonated, leaving the *ε*-nitrogen free to form a HB with the indole -NH of x1093, reversing the HB polarity compared to the other fragments in the cluster. Nonetheless, the same binding geometry is observed in both cases and the clustering algorithm correctly assigns the molecules into the same cluster. Overall, the primary functionality that facilitates interaction with His-163 is the nitrogen-containing heterocycle present in almost all fragments in cluster 5 (**Figure 16**); the exception is x0967, which forms the His-163 HB via its phenol oxygen. Such heterocycles are well suited to replace the substrate P1 Gln sidechain by mimicking its HB donor/acceptor abilities. In addition, most cluster 5 binders also reach into the hydrophobic S2 pocket, although no clear trend in functional group preference for S2 can be observed for cluster 5. This is in accord with our plasticity analysis, which shows that the S2 subsite residues are plastic and can accommodate a large variety of inhibitors. As seen in the overlap of peptide inhibitor p13 and cluster 5 representative x0678 (**Figure 15e**), the binding mode of both inhibitors in the S1 and S2 subsites are very similar, with both compounds forming HBs to His-163 and Glu-166 and binding deep in the S2 pocket.

**Figure 16:**
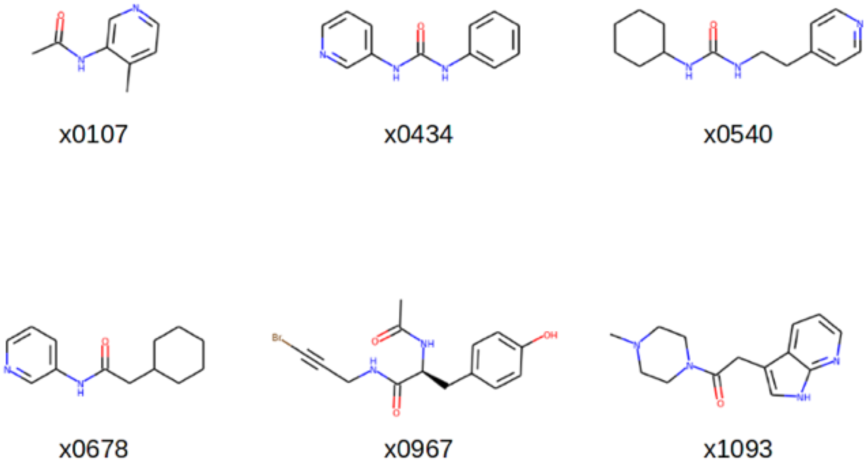
Structures of the cluster 5 XChem compounds. Note the prevalence of nitrogen-containing heterocycles, and the phenol-containing x0967.

The interactions between the fragments and M^pro^ were also analysed by employing linear scaling DFT, as for the substrates and designed peptides. Using short-range (*E*_cont_) DFT interactions with M^pro^ as a “descriptor” for clustering, a cluster containing both the substrates and the cluster 5 compounds (x0107, x0434, x0540, x0678, x0967 and x1093) was identified (**Figure S4.6**). This cluster also includes compounds x0426, x0946, x0195, x0995, x0104, x0874, x1077, x0161 and x0397. This analysis reveals fragments with similar interaction patterns to those exhibited by substrates. Unlike the Arpeggio-based analysis, BigDFT clustering can group fragments together that exhibit long-range interaction patterns that are similar to the natural substrates (highlighted in purple in **Figure S4.6**), identifying compounds that could potentially lead to tighter binding. This agnostic analysis provides a new and potentially powerful way to evaluate compounds of differing sizes and nature based on quantum mechanical descriptors obtained from biomolecular complexes.

Cluster 5 binders are therefore promising building blocks for substrate-competing inhibitor design. On the other hand, cluster 1 binders make up the bulk of covalent compounds in the fragment dataset and should therefore be considered when designing covalent inhibitors.

### 4.2 Interaction analysis of COVID Moonshot compounds

As of the 11th of January 2021, the COVID Moonshot M^pro^ inhibitor project had reported fluorescence assay data for 798 inhibitors and rapid fire assay data for 784 inhibitors,^18,33^ 245 of which had X-ray crystal structures in complex with M^pro^. To test our hypothesis that the interactions identified in the substrate and fragment analyses are important for inhibitor design, the Moonshot structures were processed with Arpeggio and ligand-M^pro^ interaction fingerprints were generated and compared to the most important interactions identified in fragment cluster 5. A Moonshot compound was classified as a cluster 5 binder if it shared at least 70% of the atom contacts found in cluster 5 binders. This analysis resulted in 101 assayed cluster 5 binders (**Figure 17**). Since the sensitivity of the activity assays are capped at 99 μM, weak binders could not be accurately quantified and were labelled with IC_50_ values of 99 μM.^33^ In the following analysis the fluorescence assay data (N=798 inhibitors) was used.

**Figure 17:**
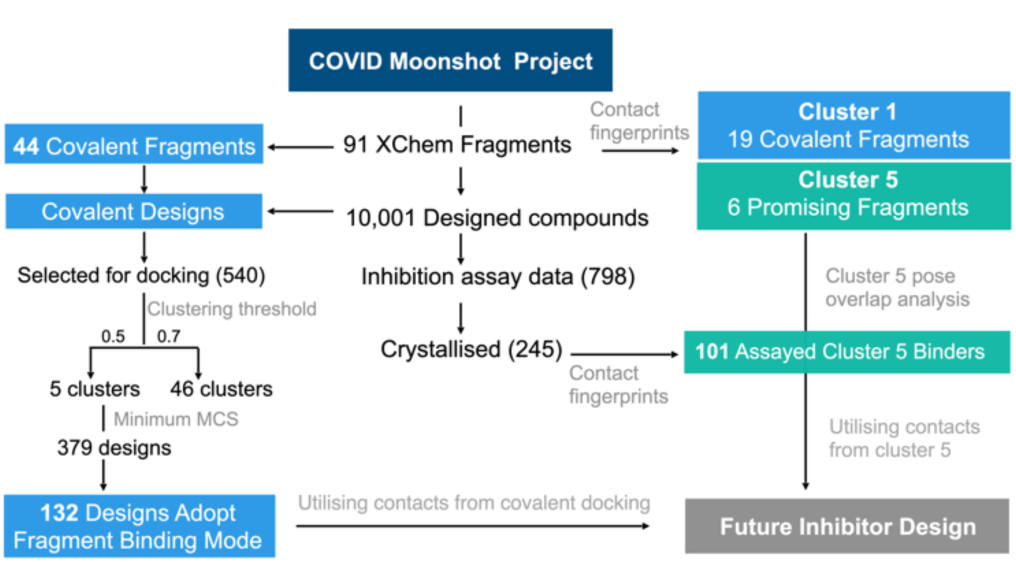
Analysis of fragment and designed compounds from the Moonshot project. Workflow used to identify promising fragments and guide novel designs.

When comparing the average IC_50_ of cluster 5 binders with the rest of the dataset, cluster 5 binders bind slightly more tightly with an average IC_50_ of 42.2 μM (95% confidence interval: 34.55-49.87 μM), while the average IC_50_ of all compounds is 54.0 μM (95% confidence interval: 50.95-57.07 μM). However, given the bimodal distribution of the data (see Figure S4.7), a better metric is the hit rate of good compounds and the number of ‘inactive’ compounds in the dataset: 2 of the 10 best Moonshot compounds are cluster 5 binders, as are 10 of the top 10% (81 compounds) of Moonshot compounds. Out of the 101 cluster 5 binders, only 15 were identified as weak binders, while out of all 798 assayed compounds 263 (33%) were weak or non-binders. As a result, this observed IC_50_ difference between cluster 5 binders and other inhibitors might be higher than currently calculated. Note, only 245 of the 798 assayed compounds had been structurally solved so it is unknown how many of the unsolved compounds are also cluster 5 binders.

Thus, by analysing fragment and substrate interactions with M^pro^, we were able to extract and cluster structural interaction fingerprints and found in cluster 5 a promising non-covalent binding mode in the M^pro^ active site. Analysis of the large set of assayed Moonshot compounds reveals that cluster 5 binders are more potent inhibitors with a higher hit rate of stronger actives than the rest of the dataset. Cluster 5 binders represent highly promising starting points for further optimisation.

### 4.3 Covalent docking of COVID Moonshot compounds

We selected the 540 covalently reacting compounds of the 10,001 designed Moonshot compounds and docked them using the knowledge-based pre-alignment docking method using AutoDock4 (**Section S1.10**).^69^ These compounds were designed by the Moonshot consortium, usually using one or more of the 44 covalent fragments as a core structure.^33^ Only compounds with a matching covalent warhead to the inspiration fragment that also cite a single covalent fragment as their inspiration were selected to form the dataset of 540 compounds. To take advantage of the many diverse induced-fit conformations of M^pro^, each designed compound was docked into the M^pro^ structure of the corresponding covalent “inspiration fragment”.^33^ The lowest energy docked pose in the highest populated cluster of each docking run was used to identify contacts using Arpeggio and generate the interaction Tanimoto distance matrix using a Tanimoto similarity threshold of 0.5 or 0.7 between poses of each cluster as described above for the XChem fragments. The broader clustering threshold of 0.5 leads to a total of 5 clusters, with the first cluster containing 477 of the 540 poses (88%) and no single-pose clusters; while the tighter threshold of 0.7 results in 46 clusters with 12 single molecule clusters. As expected, the diversity of binding modes of these compounds is much lower than in the original XChem fragment set, due to the limited number of fragments (44) and the reduced structural diversity of the designs, all being covalent S1/S1’ binders.

We subsequently analysed the ability of the procedure to recapitulate the binding pose of the parent fragment when docking the fragment-based Moonshot designs. We compared the shape and pharmacophoric (SuCOS) overlap of the lowest energy pose of the highest populated cluster for each Moonshot compound with the inspiration covalent XChem fragment referenced by the designers (**Figure S4.8**). A SuCOS score of 0.5 and higher was sufficient to consider the binding poses of the crystallographic fragment and docked design as conserved.^70^ Due to creative freedom in the design process, some of the designed compounds do not overlap significantly with the inspiration fragments and only have the covalent warhead in common in some extreme cases. When controlling for the smallest maximum common substructure (MCS) that encompasses at least the covalent warhead and one additional atom in the compound, 379 docked designs remain, from which 132 (34.8%) recovered the binding mode of the inspiration fragment. Given the high similarity between the fragments and the docked designed compounds, it is likely that these binding modes are more representative of the actual binding mode of the ligand.

At the point of analysis, only 6 of the 540 docked covalent compounds have been analysed crystallographically.^54^ These structures were used as a limited benchmark for the docking method; an overlay of the crystallographic conformation, the lowest energy pose of the highest populated cluster from docking, and the corresponding crystallographic structure of the inspiration fragment is shown in the SI (**Figure S4.9**). x10899 (**Figure S4.10**) was excluded from further analysis since it binds via a crystal contact to a third symmetry-related M^pro^ molecule, rather than the biologically relevant dimeric state.^4^ The binding modes of two compounds, x3077 and x10306 (**Figure S4.9a** and **S4.9e** respectively), were reproduced perfectly. The method places the aromatic sidechain of x3324 (**Figure S4.9b**) correctly into the S2 pocket of M^pro^ but varies on placement of the linker when compared to the crystal structure. However, in this case, the original fragment that was cited as inspiration has no overlap with the designed compound. As a result, the induced fit shape of the active site of the M^pro^ structure used for docking complements a completely different molecule and the alignment before docking pointless. For x3325 and x10172 (**Figure S4.9c** and **S4.9d** respectively), the selected lowest-energy pose of the highest populated cluster did not match the binding pose of the crystal structure.

In summary, the docking method identifies the correct binding mode when good overlap exists between the inspiration fragment and designed compound beyond the covalent warhead., However, it struggles to dock larger, more extended molecules or compounds where small changes substantially affect binding. For example, for designed compound x3325 and the fragment used as its inspiration, x1386, the change from the thiophene group to the acetylene group in x3325 leads to a major change in binding mode when comparing their crystal structures (**Figure S4.9**). Perhaps a more-human guided approach to docking (using iMD-VR) could be useful for these larger molecules, where chemical and spatial intuition can influence the resulting docked structure. As more structures become available, it will be possible to further validate the use of parent-fragment induced-fit structures in docking. The performance of our docking method appears to be in line with the active-guided docking hypothesis, where an induced fit structure and pre-alignment of fragment-derived compounds to known fragment crystal structures is only valid if the compound does not differ drastically from the parent fragment. All poses of the 540 docking runs and their SuCOS scores are available in the **SI** Files.

### 4.4 Implications for future inhibitor design

With a view to designing novel inhibitors, we compared the interactions of the cluster 5 binders observed in the crystal structures of the Moonshot designs, and those observed with the substrate peptides and the XChem fragments. Interestingly, unlike substrates, almost none of the cluster 5 binders interact with the oxyanion hole. The only cluster 5 compounds where this contact is made are a series of covalent inhibitors, none of which showed promising potency (**Figure S4.11**). An exhaustive search of Moonshot structures showed that no non-covalent inhibitor has ever been tested that includes both the typical cluster 5 binding mode while also being able to interact with the oxyanion hole.

We then compared the structures of the top 10 compounds in cluster 5 (part of the dataset analysed in **Section 4.2**) to the docked structures of covalent Moonshot designs (**Section 4.3**). Two docked compounds (FOC-CAS-e3a94da8-1 and MIH-UNI-e573136b-3) were selected based on the high SuCOS overlap with their inspiration fragments, strongly suggesting that the docked binding mode reflects the actual pose of the ligand. Both docked compounds bind into the oxyanion hole as well as into S1 and S2, providing a clear opportunity for extension of the cluster 5 binders **(Figure S4.12)**. Most cluster 5 binders connect the aromatic heterocycle binding into the S1 site with the carbonyl hydrogen bonding to Glu-166 through an amide linker. The position of this amide nitrogen overlays perfectly with the ring amine present in the docked compound FOC-CAS-e3a94da8-1 (**Figure 18a**). As a result, extension of cluster 5 binders into the oxyanion hole might best be performed through the addition of a substituent at the amide nitrogen, circumventing the issue of stereochemistry present in the covalent designs FOC-CAS-e3a94da8-1 and MIH-UNI-e573136b-3, where the connection of the pyridine to the core six-membered ring creates a stereocenter on the core ring. The most promising basis for the extension is perhaps x10789, which makes a HB with the backbone oxygen of Glu-166 (**Figure 18a**).

**Figure 18:**
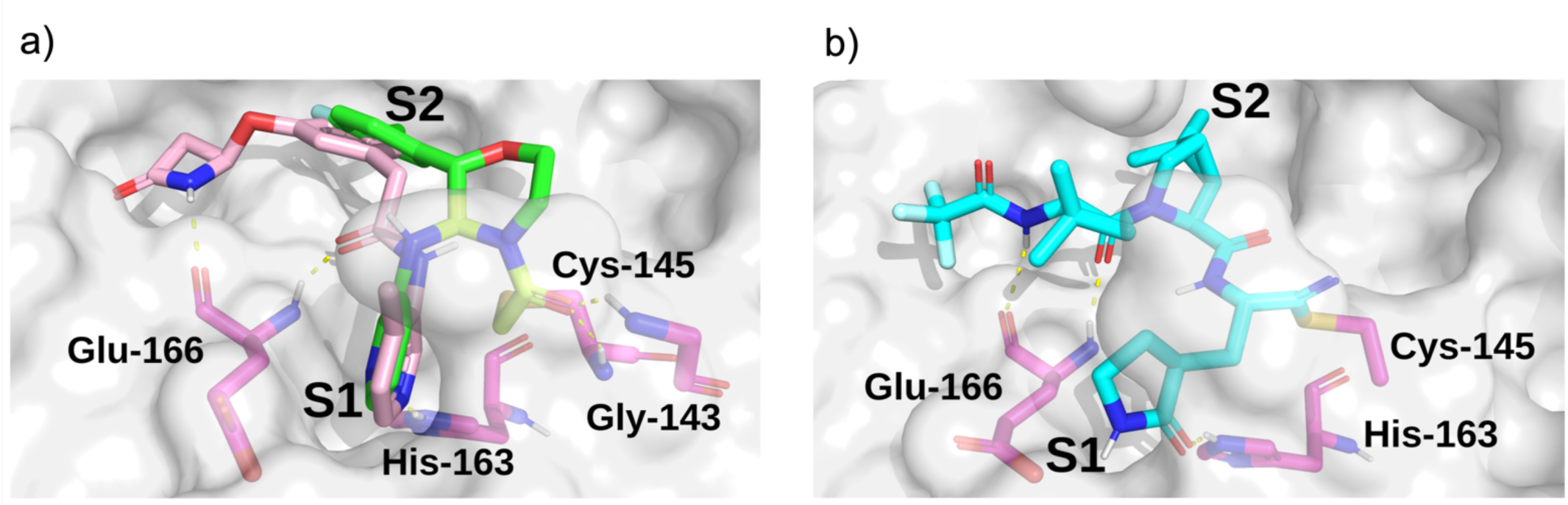
Docking informs novel inhibitor design. a) Overlay of the docked pose of FOC-CAS-e3a94da8-1 (green) with the crystal structure of x10789 (salmon) on the M^pro^ surface (PDB entry 5RER; 1.88 Å resolution).^67^ HBs between selected residues and the ligands are shown as dotted yellow lines. Extension of x10789 into the oxyanion hole could be achieved by attaching a methylene amide group present in x0830 (highlighted in yellow). b) Docked pose of Pfizer’s Phase I covalent inhibitor PF-07321332, covalently docked to M^pro^ (PDB entry 6XHM; 1.41 Å resolution).^71^ HBs between selected residues and the ligands are shown as dotted yellow lines, and protein residues are in magenta. PF-07321332 (cyan) is covalently attached to Cys-145. The docked pose of PF-07321332 adopts the same major contacts as the ‘combination’ of x10789 and x0830, namely the double HB to the backbone of Glu-166, the HB to His-163 in the S1 subsite, and a series of hydrophobic interactions in the S2 subsite.

When comparing interactions exhibited by cluster 5 binders (Glu-166, His-163) or covalent fragments (Gly-143, Cys-145) with the contacts present in the docked structure of the recently published Phase 1 clinical trial candidate PF-07321332 by Pfizer (**Figure 18b**),^11,12^ we see an almost identical interaction pattern to the cluster 5 binding motif. However, it is notable that for reacted PF-07321332, AutoDock4 was unable to place the negatively charged azanide in the oxyanion hole, which would be its expected position given its similarity to related warheads previously described and docked (see **SI**), where a carbonyl oxygen is almost always docked correctly in the oxyanion hole.

## 5 Conclusions

A wealth of crystal structures of SARS-CoV-2 M^pro^ is available, including hundreds with ligands. There is thus the question of how best to use this static information to help develop M^pro^ inhibitors optimised in terms of efficacy and safety for COVID-19 treatment. The dimeric nature of M^pro^, coupled with its multiple substrates, makes it challenging to understand the structural and dynamic features underpinning selectivity and catalysis, as is the case for many other proteases. Such an understanding is, of course, not essential to develop medicines, as shown by work with other viral proteases. Still, it may help improve the quality of such medicines and the efficiency with which they are developed. It will also lay the foundation for tackling anti-COVID-19 drug resistance — a challenge we will likely encounter as experience with the HIV global pandemic implies. The scale of global efforts on M^pro^ makes this system an excellent model for collaborative efforts linking experimental biophysics, modelling, and drug development (**Figure 19**).

**Figure 19:**
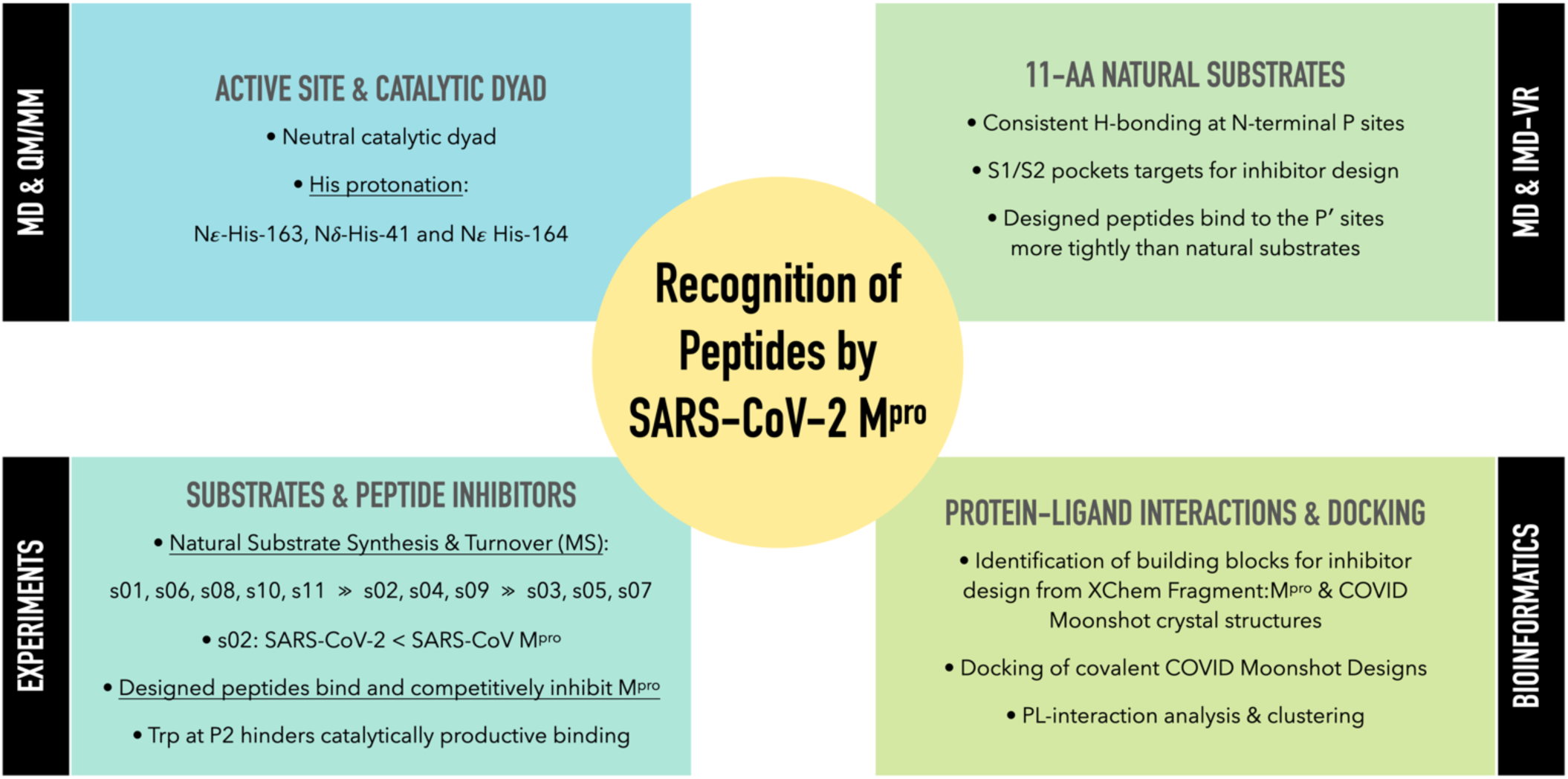
Summary of the research and results obtained in this work.

The results of our combined computational studies, employing classical molecular mechanics and quantum mechanical techniques, ranging from automated docking and MD simulations to linear-scaling DFT, QM/MM, and iMD-VR, provide consistent insights into key binding and mechanistic questions. One such question concerns the protonation state of the ‘catalytic’ His-41/Cys-145 dyad, an important consideration in the rate of reaction of covalently linked M^pro^ inhibitors which ultimately relates to their selectivity and potency. Our results indicate that a neutral catalytic dyad is thermodynamically preferred in M^pro^ complexed with an unreacted substrate, justifying the neutral state for MD simulations. A more reactive thiolate anion may be deleterious to the virus, as it will be susceptible to reaction with electrophiles. Importantly, analysis of the active site suggests that the precise mechanism of proton transfer in the His-41/Cys-145 dyad involves dynamic interactions with other residues, including His-163, His-164, Asp-187, and a water hydrogen bonded to the latter two residues and His-41. Proton transfer may be considered a relatively simple part of the overall catalytic cycle — these results thus highlight how M^pro^ catalysis is likely a property of (at least) the entire active site region, with a future challenge being to understand motions during substrate binding, covalent reaction, and product release.

The models we have developed of M^pro^ in complex with its 11 natural substrates provided a basis for analysis of key interactions involved in substrate recognition and for comparison with (potential) inhibitor binding modes. Notably, the P’ (C-terminal) side of substrates appears to be much less tightly bound than the P (N-terminal) side, where there is remarkable consistency in the hydrogen bonding patterns across the substrates. This difference may in part reflect the need for the P’ side to leave (at least from the immediate active site region) after acyl-enzyme complex formation and prior to acyl-enzyme hydrolysis. The tighter binding of the N-terminal P-side residues suggests these are likely more important in substrate recognition by M^pro^. This is also reflected in potent inhibitors, such as N3 and peptidomimetic ketoamides,^2,4^ which predominantly bind in the non-prime S subsites. The development of S-site-binding inhibitors may also reflect the nature of the substrates used in screens leading to them, which typically comprise an S-site binding peptide with a C-terminal group enabling fluorescence-based measurement. Our results imply that there is considerable scope for developing inhibitors exploiting the S’ subsites, or both S and S’ subsites, though relatively more effort may be required to obtain tight binders compared to targeting the S subsites.

Consistent with prior studies, our work highlights the critical role of the completely conserved P1 Gln residue in productive substrate binding and analogously in inhibitor binding. However, the nature of the P2/S2 interaction is also important in catalysis. In the natural substrates (**Figure 1**), the P2 position is Leu in 9 of the 11 substrates, Phe in s02 (which displays medium turnover efficiency), and Val in s03 (which is a poor substrate). Our results show that the S2 subsite plays a critical role in recognition and inhibition. This site is highly plastic (**Figure 7**) and can accommodate a range of different ligands, including larger groups, though not necessarily in a productive manner. The observation that substrates with a P2 Leu vary in efficiency reveals that interactions beyond those involving P1 and P2 are important, reinforcing the notion that (likely dynamic) interactions beyond the immediate active site are important in determining selectivity both in terms of binding and rates of reaction of enzyme-substrate complexes. Notably, the results of computational alanine scanning mutagenesis, aimed at identifying peptides that would bind more tightly than the natural substrates, led to the finding that substitution of a Trp at P2 ablates hydrolysis creating a substrate-competing inhibitor. There is thus scope for the development of tight binding peptidic M^pro^ inhibitors, for use in inhibition and mechanistic/biophysical studies.

Finally, the combined analysis of interactions involved in substrate binding and extensive structural information on inhibitor/fragment binding to M^pro^ enabled us to identify a cluster of inhibitors whose interactions relate to those conserved in substrate binding (*e*.*g*., involving the Glu-166 backbone, His-163 sidechain, and/or the oxyanion hole formed by the Cys-145 and Gly-143 backbones). Building out from these ‘privileged’ interactions (**Figure 18**) might be a useful path for inhibitor discovery. Indeed, an M^pro^ inhibitor now in clinical trials^11,12^ exploits the same ‘privileged’ interactions that we identified. We hope the methods and results that have emerged from our collaborative efforts will help accelerate the development of drugs for treatment of viral infections, and particularly COVID-19.

## 6 Methods

A detailed description of the experimental and computational methods employed in this work is provided in the supporting information. Further information has also been made publicly available in GitHub (https://github.com/gmm/SARS-CoV-2-Modelling).

## 7 Author Contributions

R.M.T. carried out QM/MM studies with subsequent contributions from K. Ś. and V.Mo.. G.M.M. generated natural substrate structures. H.T.H.C. carried out classical MD simulations and ADCP peptide docking. R.K.W. and H.M.D. performed iMD-VR simulations. M.A.M. performed protein-substrate, protein-peptide inhibitor and protein-ligand contact analysis, fragment clustering, COVID Moonshot covalent docking and future inhibitor suggestions. M.A.M. and L.G. performed bioinformatic study of fragment binding to M^pro^. L.G. performed linear-scaling QM-DFT calculations, and W.D., T.N. and T.J.W. contributed with its setup and analysis. D.K.S. carried out mutagenesis analysis and designed novel peptides (using software devised by R.B.S.). E.S., P.L., C.S.D., C.D.O. and M.A.W. carried out M^pro^ production and purification. T.R.M. and T.J. synthesised and purified natural and designed peptides. T.R.M. also performed kinetic analyses. V.M. undertook non-denaturing MS analyses. V.Mo. and A.L. contributed to discussions. C.J.S., A.J.M., D.K.S., F.D. and G.M.M. conceptualised and supervised the study. H.T.H.C., R.K.W., M.A.M., T.R.M., R.M.T., V.M., C.J.S., D.K.S., L.G., A.J.M., F.D. and G.M.M. wrote the manuscript.

## 9 Acknowledgements

We thank Prof. Michel Sanner for helpful discussions about AutoDock CrankPep. H.T.H.C. thanks the EPSRC Centre for Doctoral Training in Synthesis for Biology and Medicine (EP/L015838/1) and the Clarendon Scholarship. M.A.M. thanks the EPSRC University of Oxford Mathematics, Physical, and Life Sciences Division (MPLS) Doctoral Training Partnership (DTP) Grant Number EP/R513295/1 and GlaxoSmithKline. R.K.W. thanks the EPSRC for a PhD studentship. T.R.M. thanks the BBSRC via BB/M011224/1, and Dr Anthony Aimon for dispensing peptibitors using ECHO 550 for the mode of inhibition studies. R.M.T. and T.J.W. acknowledge the EPSRC Centre for Doctoral Training in Theory and Modelling in Chemical Sciences (EP/L015722/1). T.J. was supported by the Oxford-GSK-Crick Doctoral Programme in Chemical Biology, EPSRC (EP/R512060/1) and GlaxoSmithKline. E.S. thanks Anastasia Kantsadi and Prof. Ioannis Vakonakis for providing the M_pro_ plasmid in a pFLOAT vector. D.R.G. acknowledges funding from the Royal Society (URF\R\180033). L.G. thanks Michel Masella for useful discussions and for the force field comparison. L.G., W.D. and T.N. gratefully acknowledge the CEA-RIKEN collaboration. J.S. thanks A.J.M. thanks EPSRC (EP/M022609/1). A.J.M. and D.K.S. thank BrisSynBio, the BBSRC and EPSRC Synthetic Biology Research Centre (BB/L01386X/1). A.J.M., J.S. and H.M.D. thank the British Society for Antimicrobial Chemotherapy for support (BSAC-COVID-30). G.M.M. acknowledges the EPSRC and MRC for their indirect support via EP/S024093/1 and EP/L016044/1. This project made use of time on JADE granted via the UK High-End Computing Consortium for Biomolecular Simulation (HECBioSim, hecbiosim.ac.uk), supported by the EPSRC via EP/P020275/1. MD simulations were also carried out using the computational facilities of the Advanced Computing Research Centre, University of Bristol (http://www.bris.ac.uk/acrc) under an award for COVID-19 research. D.K.S. and A.J.M. thank EPSRC via HECBioSim for providing ARCHER/ARCHER2 time through a COVID-19 rapid response call; and Oracle Research for Oracle Public Cloud Infrastructure time under an award for COVID-19 research (http://cloud.oracle.com/en_US/iaas). L.G., W.D. and T.N. acknowledge RIKEN through the HPCI System Research Project (Project ID: hp200179 and hp210011) for computer time at the Fugaku supercomputer facility. L.G. is also grateful to the TGCC of CEA for granting of compute hours (gch0429 and gen12047 projects), and to the MaX Center of Excellence. C.J.S. thanks the Wellcome Trust, Cancer Research UK and the Biotechnology and Biological Sciences Research Council for funding. This research was funded in whole, or in part, by the Wellcome Trust (grant no. 106244/Z/14/Z). For the purpose of open access, the author has applied a CC BY public copyright license to any Author Accepted Manuscript version arising from this submission.

