## Supplementary Information for "Discovery of SARS-CoV-2 M^pro^ Peptide Inhibitors from Modelling Substrate and Ligand Binding"

### S1. Methods

#### S1.1 QM/MM studies on catalytic dyad protonation state

A crystal structure of the TSAVLQ↓SGFRK (↓ indicating scissile amide bond) substrate peptide-bound inactivated H41A mutant of SARS-CoV BJ01 M$^{pro}$ (PDB entry 2q6g; 2.50 Å resolution)[1] was used as a starting point for modelling SARS-CoV-2 M$^{pro}$ (M$^{pro}$) in complex with the substrate peptide s01. The mutated residue Ala-41 was converted back to His-41 using the automodel routine in Modeller.[2] Crystal waters were conserved, and hydrogens were added in silico. Protonation states of titratable residues were assigned using PROPKA3;[3] the hydrogen bonding (HB) network was optimised using the protein preparation wizard in Maestro (Schrodinger).[4] Neutral and zwitterionic catalytic dyad models were prepared for the three possible protonation states of His-163 ($\delta$-, $\varepsilon$-, or both nitrogens protonated), resulting in six scenarios. The FF14SB force field[5] was used to describe the protein and substrate. A solvation shell of TIP3P water molecules[6] was created 5 Å around the protein using SOLVATE.[7] Further solvation was achieved by construction of a truncated octahedral cell of TIP3P water using LEaP (AmberTools19),[8] with a 10 Å distance from the initial solvation cell to the edge of the box. $Na^+$ and $Cl^-$ ions were added randomly throughout the solvent with a concentration of 0.1 M NaCl.

Each system was minimised by restraining the protein and allowing relaxation of the solvent and ions. This was followed by a minimisation restraining only the backbone atoms, a further minimisation restraining only C$\alpha$s, and a final minimisation allowing the full system to relax. Each minimisation included 1000 steps of steepest descent, followed by 10,000 steps of conjugate gradient minimisation. The systems were heated by increasing the temperature to 310 K over 100 ps. Langevin dynamics was used with a collision frequency of 5 $ps^{-1}$, and backbone atoms were restrained with a force constant of 5 kcal $mol^{-1}$ $Å^{-2}$. SHAKE was used to restrain bonds to hydrogen atoms.[9] A timestep of 2 fs was used. NPT equilibration was carried out for a total of 9 ns, slowly releasing backbone restraints, with a Monte Carlo barostat maintaining the pressure at 1.01325 bar. Three repeat simulations of 250 ns of production MD were carried out on each of the six systems using pmemd.cuda (AMBER18).[8]

The following QM/MM protocol was followed to carry out umbrella sampling simulations of the proton transfer (PT) between Cys-145 and His-41. Simulations were performed using sander.MPI (AMBER18).[8] The QM region consisted of the sidechains of Cys-145 and His-41. The reaction coordinate was defined as the difference between the S-H and N-H distances and was restrained with a force constant of 50 kcal $mol^{-1}$ $Å^{-2}$. The reaction coordinate describing the PT was varied between -1.0 Å and 1.0 Å in steps of 0.1 Å, where a value of -1.0 Å denotes the neutral catalytic dyad and a value of 1.0 Å denotes the zwitterionic catalytic dyad. The reaction coordinate was followed in the forward and reverse directions, with starting snapshots selected from the MD trajectories of the neutral and zwitterionic catalytic dyad, respectively, to ascertain if hysteresis affects the free energy surface for PT. A total of 100 ps of sampling at the DFTB3/MM[10] level of theory was carried out in each reaction coordinate window, but the first 25 ps of sampling was treated as equilibration and discarded. A further backward run of 5 ps of sampling per window was carried out at the ωB97X-D/6-31G*/MM[11-13] level of theory for the structure with HID-163 using the interface to Gaussian 16.[14]

#### S1.2 Comparative modelling of the SARS-CoV-2 M$^{pro}$-peptide complexes

The crystal structure of the TSAVLQ↓SGFRK 11-mer peptide substrate bound to the H41A mutant of dimeric SARS-CoV M$^{pro}$ (PDB entry 2q6g)[1] was superimposed with a crystal structure of unmodified dimeric SARS-CoV-2 M$^{pro}$ (PDB entry 6yb7; 1.25 Å resolution).[15] The substrate was transferred over to the chain A active site of the catalytically-competent SARS-CoV-2 M$^{pro}$ structure. The sequences of the 11 native cleavage sites processed by SARS-CoV-2 M$^{pro}$ (s01-s11) were identified by aligning the sequences of the ORF1ab polyproteins of both SARS-CoV (GenBank accession code NC_004718.3)[16] and SARS-CoV-2 isolate Wuhan-Hu-1 (accession code MN908947.3)[17] using MUSCLE.[18] For each of the 11 cleavage sites, atomic models of an 11-mer peptide matching positions P6 to P5′ and charged N- and C-termini were constructed using the mutagenesis tool of the open source version of PyMOL (v. 2.3.0).[19] For every sidechain from positions P6 to P5′, apart from Gly and Ala, the highest-probability backbone-dependent conformer with the least steric clash and the most chemical complementarity was selected.[20] Using CCG MOE version 2019.0104,[21] each of the resulting 11 models of the SARS-CoV-2 M$^{pro}$ dimer complexed with each 11-mer substrate in the A-chain active site underwent structure preparation protonation using Protonate 3D. Each model was then solvated using 0.1 M NaCl and explicit water and subjected to energy minimization using the AMBER10:EHT force field[22,23] and periodic boundary conditions, until convergence with an RMS of 0.4184 kcal $mol^{-1}$ per iteration was reached.

#### S1.3 Explicit-solvent molecular dynamics

Pre-solvation models of the dimeric M$^{pro}$-peptide complexes constructed as described above (**Section S1.2**) were used as starting points for MD simulations. All additives and crystallographic water molecules were removed from PDB entry 6yb7, except HOH 644 which provides bridges between His-41, His-164 and Asp-187 (see main text for details). Protonation and rotameric states of histidines and other titratable residues were assigned at pH 7.4 based on a combination of Reduce (MolProbity, Duke University),[24] H++ (Virginia Tech),[25] PROPKA3 (PDB2PQR),[3] and visual inspection, with a final M$^{pro}$ monomeric charge of -4. His-41 (protonated on its $\delta$-nitrogen) and Cys-145 were assigned neutral.

MD simulations were performed using GROMACS (v. 2019.2)[26] employing the AMBER99SB-ILDN force field.[27] Each of the constructed complexes was solvated (TIP3P water model)[6] in a rhombic dodecahedral box (1.0 nm buffer), neutralised, and minimised using the steepest descent algorithm until the maximum force was below 1000 kJ mol$^{-1}$ nm$^{-1}$. For each peptide sequence, three independent simulations were initiated by random velocities at 298.15 K. In each case, the system was equilibrated under NVT (200 ps; 1 fs timestep) and NPT (200 ps; 1 fs timestep) conditions, before being subjected to 200 ns MD simulation (2 fs timestep) at 298.15 K and 1 bar, during which protein-peptide interactions were monitored. All simulations were performed with three-dimensional periodic boundary conditions. Long-range electrostatics was calculated using the smooth particle mesh Ewald method.[28] All bond lengths involving hydrogen atoms were constrained with the LINCS algorithm.[29] Hydrogen bonds between M$^{pro}$ and the peptides were monitored over the course of the simulations, defined using a combined criteria on the donor-acceptor distance ($d_{D\text{-}A} \leq 3.5$ Å) and the proton-donor-acceptor angle ($\angle$(H-D-A) $\leq 30°$).

Models of the designed sequences p12 and p13 complexed with SARS-CoV-2 M$^{pro}$ (PDB entry 6yb7) were built using a comparative modelling approach similar to that described above (**Section S1.2**), starting from the previously constructed model of the M$^{pro}$-s02 complex. Each constructed complex was then solvated, minimised, equilibrated, and subjected to 3 $\times$ 200 ns MD as described above, except the retention of a backbone restraint during NPT equilibration to allow longer relaxation of the non-native peptide side chains and the M$^{pro}$ binding pockets.

To generate representative structures of M$^{pro}$-peptide complexes for interaction analysis, frames extracted every ns from the concatenated 3 $\times$ 200 ns MD trajectories were fitted using the M$^{pro}$ backbone, before performing clustering based on the heavy-atom RMSD of the peptide, using the gromos algorithm as implemented in GROMACS (v. 2019.2).[30] A cut-off of 2.0 Å (for native substrates) or 2.5 Å (for p12 and p13, due to heavier residues in their terminal regions) was used.

#### S1.4 Interactive Molecular Dynamics in Virtual Reality (iMD-VR) and subsequent implicit-solvent MD

The same crystal structure of apo dimeric SARS-CoV-2 M$^{pro}$ (PDB entry 6yb7)[15] was used as the target for substrate and peptide inhibitor docking using iMD-VR. Protonation states of histidines and other titratable residues were the same as described in main text **Section 2.1**. The M$^{pro}$, three natural substrates (s01, s02, s05), and two peptide inhibitors (p12 and p13) tested were parameterised using the LEaP programme (AMBER19)[8] employing the AMBER99SB-ILDN force field[27] and the OBC2 implicit solvent water model (igb=5).[31] M$^{pro}$ was minimised using OpenMM[32] prior to iMD-VR simulation.

For all iMD-VR simulations, a temperature of 300 K was used with a timestep of 0.5 fs. M$^{pro}$, all substrates, and both peptide inhibitors remained fully flexible. Whilst in VR, each substrate and peptide inhibitor was docked to M$^{pro}$ following the guidance of 'trace atoms' representing where the s01 substrate should bind; this visual representation was taken from the positions of the s01 backbone atoms in the crystal structure of the H41A mutant of SARS-CoV M$^{pro}$.[illegible][1][illegible] These 'trace atoms' were used as a rough visual guide to aid the docking, and the main focus whilst in VR was on establishing key hydrogen bond contacts between the protease and the three substrates and the protease and the two peptide inhibitors.

Once each substrate was docked in VR, a structure where the oxyanion hole interactions were successfully reformed was extracted, and these structures were minimised, equilibrated, and subjected to 3 x 200 ns replicates of production MD in implicit solvent to ensure the substrates remained bound. In the case of the docked peptide inhibitors, the docking was repeated 5 times, and structure where the oxyanion hole interactions were successfully reformed was extracted, resulting in 5 docked structures per peptide inhibitor. The 5 docked structures were minimised, equilibrated, and subjected to 500 ns of production MD in implicit solvent. The process of minimisation and equilibration was the same for all substrate and peptide inhibitor structures, and is as follows: First, the structures were iteratively energy minimised at 10 K using slowly decreasing degrees of positional restraint. Restraints of 5 kcal mol$^{-1}$ Å$^{-2}$, 2.5 kcal mol$^{-1}$ Å$^{-2}$, and 1.25 kcal mol$^{-1}$ Å$^{-2}$ were applied to all backbone atoms for the first three rounds of minimisation respectively, and no restraints were applied for the final round. The system was heated by running 10 stages totaling 20 ps of MD, starting at 0 K and linearly increasing the temperature by 30 K at each stage until a temperature of 298 K was reached (each step had a backbone atom restraint of 5 kcal mol$^{-1}$ Å$^{-2}$). 8 rounds of 500 ps of NPT MD with slowly decreasing backbone restraints were run. Restraints were initially 5 kcal mol$^{-1}$ Å$^{-2}$ and halved after each step; once backbone restraints were below 1 kcal mol$^{-1}$ Å$^{-2}$, only the restraints on C$\alpha$ atoms were retained. The eighth and final stage had no restraints in the system at all.

For the docked substrate structures (s01, s02, and s05), following minimisation and equilibration, 3 x 200 ns replicates of production MD in OBC2 implicit solvent[31] was run for each docked substrate structure, with a protein backbone restraint of 5 kcal mol$^{-1}$ Å$^{-2}$, resulting in 3 x 200 ns MD trajectories for each substrate. In the case of the peptide inhibitors, following minimisation and equilibration, 500 ns of production MD in OBC2 implicit solvent[31] was run for each iMD-VR docked peptide structure, with a protein backbone restraint of 5 kcal mol$^{-1}$ Å$^{-2}$, resulting in 5 x 500 ns MD trajectories for each peptide inhibitor (due to 5 independent docked structures from iMD-VR).

#### S1.5 Contact interaction mapping

#### S1.5.1 Contact maps

Snapshots from MD models as well as XChem crystal structures and covalent docking poses were analysed using Arpeggio.[33] The ligand-$M^{pro}$ complex was processed as described by Jubb et al.[33] by cleaning the PDB file using PDBtools[34] and running Arpeggio on all ligand-$M^{pro}$ contacts. For the MD snapshots of the substrate and designed peptides, a representative snapshot for each complex was chosen by selecting the highest populated cluster and the conformation within the cluster that has the lowest RMSD to all other snapshots in the cluster. From the docked covalent Moonshot submission compounds, the lowest energy pose of the highest populated cluster was chosen. The analysis of the XChem fragments and Moonshot designs was done using the published crystallised conformation.[35-37] The resulting Arpeggio contact map consists of a bit vector for each identified atom-atom contact and classifies them as "Clash", "Covalent", "VdW Clash", "VdW", "Proximal", "Hydrogen Bond", "Weak Hydrogen Bond", "Halogen Bond", "Ionic", "Metal Complex", "Aromatic", "Hydrophobic", "Carbonyl", "Polar" or "Weak Polar".[33]

#### S1.5.2 Hydrophilicity maps

To calculate whether a given protein subsite corresponds to a hydrophilic or hydrophobic pocket, a hydrophilicity score was introduced. All identified atom-atom contacts that interact with a given residue in the substrate are classified as either hydrophobic (Hydrophobic, Aromatic, Halogen Bond) or hydrophilic (Hydrogen Bond, Weak Hydrogen Bond, Ionic, Carbonyl, Polar), excluding the "VdW" and "Weak Polar" interaction types since they were deemed too insignificant and usually redundant as individual atom-atom contacts. The sum of all hydrophobic atom-atom contacts was then subtracted from the sum of all hydrophilic atom-atom contacts to create a hydrophilicity score for each subsite.

#### S1.5.3 Interaction fingerprints

A bit vector was created for every analysed protein-ligand complex, denoting the absence (0) or presence (1) of an interaction of a single ligand with every protein residue that was found to interact with any of the known actives (namely the substrates or the XChem fragments). In order to compare the interaction networks of the ligands, a Tanimoto distance can be calculated between the fingerprint bit vectors using the Jaccard distance[38] and the ligands clustered by their calculated Tanimoto distance.

To investigate potential fragment elaboration pathways, the atom-atom contacts present in each fragment cluster were used as a baseline to investigate fragment growth. To identify if a designed small molecule ligand exhibits the same binding profile as one of the identified fragment clusters, a standardised cluster profile was created for each fragment cluster which records the presence of a residue level contact if it was classified by Arpeggio as one of the following major contacts: Aromatic, Hydrophobic, Halogen Bond, Polar, Hydrogen Bond, Ionic, Carbonyl. If more than 70% of all recorded residue level contacts of a particular cluster are occupied for an individual ligand, we classify the ligand as a member of that cluster.

#### S1.6 MM-GBSA calculations

The contribution of each residue to protein-peptide binding was evaluated quantitatively using per-residue decomposition of binding energy,[39] estimated using the molecular mechanics-generalised Born surface area (MM-GBSA) method as implemented in MMPBSA.py (v. 14.0)[40] in combination with *sander* (Amber18).[8] The single trajectory protocol was employed, with the $M^{pro}$ dimer defined as the receptor and the 11-mer peptide as the ligand. Snapshots were extracted every 5 ns from the 3 $\times$ 200 ns MD trajectories for each substrate (120 frames per substrate). The polar solvation term was calculated using the OBC2 model (igb=5) with mbondi2 radii at 0.15 M salt concentration.[31] Non-polar solvation terms were computed from surface area (recursive approximation from icosahedra)[39] and a surface tension of 0.005 kcal $mol^{-1}$ $Å^{-2}$.[31]

#### S1.7 BigDFT calculations

Snapshots generated from MD (see main text **Section 2.2**) were studied by Quantum Mechanical (QM) modelling, as implemented in the BigDFT suite.[41] The approach employs Daubechies wavelets to express the electronic structure of the assemblies in the framework of Kohn-Sham (KS) formalism of Density Functional Theory (DFT).[42] With such an approach, the code provides QM results for full systems of large sizes, thanks to the systematic approach offered by wavelets. The electronic structure is expressed by both the density matrix and the KS hamiltonian operator in an underlying basis set of so-called support functions, which are a set of localised functions that are adapted to the chemical environment of the system. Such functions are then expressed in Daubechies wavelets and there are only a few per atom (between 1 and 4 according to the chemical species). The code delivers excellent performance on massively parallel supercomputers and provides the user with the possibility of treating the entire system with the same QM level of theory. A single calculation on one MD-clustered snapshot at the PBE-D3 DFT level requires about 2 h of walltime on 2048 CPU cores (16 nodes) of the IRENE-Rome supercomputer at the TGCC Supercomputing centre in Saclay (Paris). We employed this computational setup, with the inclusion of frozen-core approximation enforced by norm conserving pseudopotentials, for all the DFT calculations presented here. The information to set up the full QM calculation (input file, code version) is available in the GitHub project associated with this publication.

The electronic density matrices as well as the KS hamiltonians expressed in the BigDFT basis were analysed to provide quantum observables on the various portions of the systems. Such a method of analysis has been employed previously and has proven to be able to i) evaluate reliable physico-chemical observables on the systems' moieties, thereby decomposing an observable into fragment-

`

based pseudo-observables and ii) assess the pertinence of a given partitioning, by providing an indicator of the quality of the pseudo-observables. In particular, we have analysed the strength of the QM interaction on each of the systems' residues, calculated as the matrix elements of the KS hamiltonian reduced on the amino acids. Such analysis provides a linear-response approximation of the energetic contribution by the corresponding residue to the enzyme-peptide interaction and enables a characterisation of the chemical bonding between portions of the system.

For the XChem crystallographic positions, a scheme to equilibrate the position was used, as follows. The pdbfixer program[43] was employed to optimise the crystallographic positions. Water molecules were removed, as well as the hydrogen lost when a ligand formed a covalent bond. Only the $M^{pro}$ monomer-ligand complex was considered for this preliminary dataset. The resulting positions were then optimised with the GFN-FF force field provided by the XTB program.[44]

#### S1.8 Experimental studies on $M^{pro}$ activity and inhibition

##### S1.8.1 Protein production and purification

Protein was produced and purified as reported.[45]

##### S1.8.2 Peptide synthesis

Peptide synthesis was performed as reported.[45] s01, s01LP2W , s01QP1W, p12, p13, p13WP2L, p15 and p16 were synthesised on a 0.1 - 0.25 mmol scale from C- to N-terminus on Rink amide-MBHA resin (100–200 mesh, 0.6–0.8 mmol $g^{-1}$ loading, AGTC Bioproducts) using a microwave assisted LibertyBlue peptide synthesizer (CEM) and N-Fmoc protected α-amino acids (CS Bio, Novabiochem, Sigma-Aldrich, TCI, Alfa Aesar, Merck or AGTC Bioproducts). N,N′-diisopropylcarbodiimide (TCI Europe) and Oxyma Pure (Merck) in DMF and 20% (v/v) piperidine in DMF (peptide synthesis grade, AGTC Bioproducts) were used for iterative cycles of coupling and deprotection respectively under the manufacturer's standard protocol. Following the terminal Fmoc-deprotection step, the resin was washed with $CH_2Cl_2$, dried in air, then treated with 5-10 mL of a deprotection solution (2.5:2.5:2.5:92.5 (v/v) 1,3-dimethoxybenzene, triisopropylsilane, MilliQ water and trifluoroacetic acid) for 3 h at ambient temperature. The resulting mixture was filtered and the filtrate was diluted with cooled $Et_2O$ (3 x 45 mL) to precipitate the peptide. $Et_2O$ was decanted, peptide dried on air and lyophilised overnight.

Peptides apart from P1 mutant of s01 were dissolved in DMSO and quantified by spiking the sample with 3 mg $mL^{-1}$ of an internal standard 3-(Trimethylsilyl)propionic-2,2,3,3-d4 acid sodium salt. The eleven substrate peptides (s01-s11) were also purchased from GLBioChem (Shanghai).

##### S1.8.3 Substrate turnover analysis under denaturing conditions

20 µM stock of all 11 native substrates were prepared in the assay buffer (20 mM HEPES, pH 7.5, 50 mM NaCl). E1-ClipTip™ Bluetooth™ Electronic multichannel pipette (ThermoFisher) was used to dispense 5 µL/well (x24) of each peptide in a single row of a 384 well plate. The first column was treated with a final concentration of 1% (v/v) aqueous formic acid to obtain 0 min time point. $M^{pro}$ was dispensed using Multidrop to obtain a final concentration of 0.15 µM $M^{pro}$ with 2 µM peptides in all wells. Each column was sequentially quenched with 1% (v/v) aqueous formic acid every minute. Samples were analysed by solid-phase extraction (SPE) coupled to mass spectrometry (MS) using a RapidFire Mass Spectrometer. The operating parameters in the positive ion mode were: capillary voltage (4000 V), nozzle voltage (1000 V), fragmentor voltage (365 V), drying gas temperature (280 °C), gas flow (13 L $min^{-1}$), sheath gas temperature (350 °C) and sheath gas flow (12 L $min^{-1}$). The sample was loaded onto a SPE C4-cartridge, which was then washed with 0.1% (v/v) aqueous formic acid to remove non-volatile buffer salts (5.5 s, 1.5 mL $min^{-1}$) followed by elution with aqueous 85% (v/v) acetonitrile in 0.1% (v/v) formic acid (5.5 s, 1.25 mL $min^{-1}$). The cartridge was equilibrated with 0.1% (v/v) aqueous formic acid (0.5 s, 1.25 mL $min^{-1}$) prior to every sample injection. Data was exported in a plate list mode and processed in Excel to calculate percentage product turnover.

##### S1.8.4 Substrate binding and turnover analysis under non-denaturing conditions

Non-denaturing mass spectra were obtained using a Waters Synapt HDMS Q-TOF mass spectrometer coupled to an automated chip-based nano-electrospray ion source (TriVersa Nanomate, Advion). A larger concentration of $M^{pro}$ than the one in the denaturing MS assays was used to provide sufficient sensitivity. 5 µM of $M^{pro}$ was mixed with 13-fold molar excess of a substrate (s01-s11) in 200 mM of ammonium acetate (pH 6.9) at room temperature and electrosprayed (1.77 kV spray voltage, 0.55 psi spray backing gas pressure and 4.3 mbar inlet pressure). The sample and extractor cone voltages were kept at 180 V and 1 V, respectively; no in-source dissociation of $M^{pro}$ dimers was observed at these voltages. Mass spectra were recorded after 1, 3, 6, 9 and 12 min incubation. Measurements were taken in duplicate for each substrate. Data collection and analysis were carried out using Waters MassLynx software. Integrated peak areas of the substrate ions and cleavage product ions were compared at different time points: the sum of substrate and product ions intensities was set at 100% for each measurement, and the level of depletion of the substrate ions was used as a measure of the turnover efficiency.

##### S1.8.5 Dose response curve analysis

Methods for SPE coupled RapidFire MS-based assay are reported.[45] In brief, in a 384 polypropylene well plate, 100 µL of 2.5 mM stocks of the designed peptides were transferred. 11 point 3 fold serial dilutions of the peptides were performed in 60 µL using E1-ClipTip™

Bluetooth™ Electronic multichannel pipette (ThermoFisher) in DMSO with 5 mix cycles of 30 µL volume for mixing. 10 µL was drawn from each well and 5 µL was transferred to two wells of a new destination 384 well polypropylene plate. 5 µL of DMSO (positive control) and 5 µL of 10% (v/v) aqueous formic acid (negative control) were added to 16 wells each on every destination plate. 25 µL/well of x2 stock of enzyme in assay buffer (20 mM HEPES, pH 7.5, 50 mM NaCl) was dispensed using Multidrop Combi and incubated for 15 minutes followed by dispensation of x2 stock of substrate in each well to obtain 0.15 µM $M^{pro}$ and 2 µM s01 concentration. The reaction was allowed to progress for 10 min (~ 50% turnover in DMSO control), then quenched with 5 µL of 10% (v/v) aqueous formic acid. The plates were centrifuged for ~15 s after addition of each reagent at 2500 rpm (Star lab) to ensure all dispensed solutions were pooled at the bottom of the plate. The plates were analysed by SPE coupled MS under the conditions specified in **Section S1.8.3**. RapidFire integrator was used to extract and integrate abundance peaks of the +1 charge states of the substrate (1191.68 Da) and N-terminal cleaved product (617.34 Da). Data was exported in a plate list mode and processed in Excel to calculate percentage product turnover, normalisation of percentage activity followed by deduction of percentage inhibition. Normalised percentage inhibition data was exported to GraphPad Prism 8 and non-linear regression analysis was performed to obtain $IC_{50}$ values. Top and bottom constraints of 100% and 0% were applied respectively for the analysis of reported $IC_{50}$ values curves. Z' of the assay was always ≥0.8.

#### S1.8.6 Dose response curve analysis with varying substrate concentrations

The designed peptides were dispensed using an Echo 550 acoustic liquid handling robot. Samples were prepared as described above (**Section S1.8.5**) with final substrate concentrations of 2 µM, 10 µM, 20 µM and 40 µM TSAVLQ/SGFRK-$NH_2$ (s01) with 10, 10, 15 and 20 minutes of incubation with substrates, respectively.

#### S1.8.7 Designed peptide turnover analysis under denaturing conditions

100 µM stocks of p12, p13, p15, p16, p13WP2L, s01LP2W and s01QP1W were prepared. 0.15 µM of enzyme was dispensed and incubated with 2 µM peptide (**Section S1.8.3**); the reaction was allowed to proceed overnight at 37°C, 300 rpm in a thermomixer. Samples were analysed by SPE coupled MS. After integration using RapidFire Integrator, the data was analysed in Excel and presented using GraphPad Prism 8.

**Table S1.1**: Observed mass (Da) and (***m/z***) charge states of the peptides that were extracted using RapidFire Integrator for peak integration.

| Peptides | Sequence | substrate (Da) (*m/z* charge state) | Product (Da) (*m/z* charge state) |
|---|---|---|---|
| s01 | TSAVLQ↓SGFRK | 1191.68 (+1) | 617.34 (+1) |
| s02 | SGVTFQ↓SAVKR | 1177.65 (+1) | 637.30 (+1) |
| s03 | KVATVQ↓SKMSD | 1191.62 (+1) | |
| s04 | NRATLQ↓AIASE | 1171.55 (+1) | |
| s05 | SAVKLQ↓NNELS | 1200.57 (+1) | 644.37 (+1) |
| s06 | ATVRLQ↓AGNAT | 1099.53 (+1) | 686.36 (+1) |
| s07 | REPMLQ↓SADAQ | 1243.51 (+1) | 772.39 (+1) |
| s08 | PHTVLQ↓AVGAC | 2185.98 (+2) | |
| s09 | NVATLQ↓AENVT | 1157.60 (+1) | 644.35 (+1) |
| s10 | TFTRLQ↓SLENV | 1305.62 (+1) | 764.37 (+1) |
| s11 | FYPKLQ↓SSQAW | 1352.59 (+1) | 794.37 (+1) |
| p12 | KYTFWQYSQFY | 1558.75 (+1) | |
| p13 | KYLTWQNSQIN | 1392.70 (+1) | |
| p15 | LTINWQKYFNT | 1427.62 (+1) | |
| p16 | WFTLKQYWQTN | 1514.70 (+1) | |
| p13WP2L | KYLTLQNSQIN | 1319.71 (+1) | |

`

| | | | |
|---|---|---|---|
| s01LP2W | TSAVWQ↓SGFRK | 1264.65 (+1) | 690.33 (+1) |
| s01QP1W | TSAVLWSGFRK | 1249.68 (+1) | |

**S1.8.8 LCMS analysis for designed peptides**

LCMS experiments were performed using an Agilent Infinity Series II System attached to QTOF 6650 using an Agilent Zorbax C-18 Extend column. Solvent A: LCMS grade water with 0.1% formic acid, and solvent B: 100% acetonitrile in 0.1% (v/v) formic acid was used at 0.2 mL min$^{-1}$ flow rate to elute the peptides over a gradient of 22-55% of solvent B over 8 minutes. The operating parameters for the LCMS were the same as above (**Section S1.8.3**). In a 96 well plate, samples consisting of 0.15 µM M$^{pro}$ were prepared. p12, p13, p15, p16 and s01 were transferred from source wells to destination wells with M$^{pro}$ using the multi injector programme and samples injected immediately after mixing. 30 min, 3 h, 6 h, 1 day and 2 days time points were obtained for peptides. Samples were covered with a polypropylene cover to prevent evaporation of samples.

**S1.8.9 Designed peptide binding and turnover analysis under non-denaturing conditions**

The binding of designed peptides p12, p13, p15 and p16 to M$^{pro}$ dimers and their effects on substrate turnover were investigated using non-denaturing mass spectrometry (**Section S1.8.4**). 5 µM of M$^{pro}$ was mixed with designed peptides at different levels of peptide excess in 200 mM of ammonium acetate (pH 6.9) at room temperature. Non-denaturing mass spectra were recorded for different protein-peptide molar concentration ratios (up to 16-fold excess of peptide relative to the protein). At the final step, the native s01 substrate was added to the protein-peptide mixture at 4-fold excess over the protein, and its turnover recorded after 3- and 6-min incubation.

**S1.9 Peptide docking**

Docking of substrate and inhibitor peptides was performed using AutoDock CrankPep (ADCP) in the ADFRsuite (v. 1.0) package.[46] For redocking trials, the structure of s01-bound H41A SARS-CoV M$^{pro}$ (PDB 2q6g)[1] chain A was prepared in ADFRsuite as the receptor. For docking to SARS-CoV-2 M$^{pro}$, the N3 inhibitor-bound (PDB 7bqy; 1.70 Å resolution)[47] or the C-terminal autocleavage site product-bound C145A (PDB 7joy; 2.00 Å resolution)[48] M$^{pro}$ structure was used. The dimeric M$^{pro}$ structure was used following processing on MolProbity,[24] correction of histidine states (**Table S2.1**), and conversion to pdbqt format with ADFRsuite. The most probable peptide-binding site on the receptor surface was predicted with AutoSite (v1.1).[49] While the peptide-binding site was successfully identified with the monomeric structure from PDB 2q6g, with the monomeric structure from PDB 7bqy binding site identification was unsuccessful. When the dimeric structure (PDB 7bqy) was inputted, however, the active site was successfully identified. Hence, subsequent docking with SARS-CoV-2 M$^{pro}$ structures was performed with the dimer. Affinity maps on the receptor were calculated with AutoGridFR (v. 1.2).[50] Each docking run was performed with 100 replicas and 11 million steps, starting from the extended peptide conformation, with solutions internally clustered by a native contact threshold of 0.8. The clustered solutions were then evaluated against the binding mode found in the original structure (SARS-CoV) or in the minimised comparative model (SARS-CoV-2). To allow for flexibility in the less tightly bound terminal regions, and to filter out solutions where the peptide positioning was offset by one residue, solutions were assessed on the criteria of < 2 Å deviation in at least three C$\alpha$ atoms, after fitting to the M$^{pro}$ backbone with VMD (v. 1.9.4).[51] Out of the top 10 poses, the highest-scoring filtered pose, or if none of the poses passed the filter, the pose with the lowest C$\alpha$ RMSD, was presented.

**S1.10 Protein-ligand docking**

**S1.10.1 Dataset**

A large-scale crystal-based fragment screen against M$^{pro}$ has been conducted using the Diamond synchronton.[37] >500 fragments were screened leading to the discovery of 92 active fragments, 44 of which are covalently bound to Cys-145. The structures are made available on Fragalysis.[52] The Poster.AI Moonshot project was then started, which crowdsourced the design of inhibitors based on the original fragment screen. All submissions are made available on the Moonshot project GitHub.[36] For our covalent docking workflow, the dataset as of the 12th of July, 2020 was used, which features 10001 submissions. A subset was created by selecting only submissions with a matching covalent warhead that cite one covalent fragment as their inspiration, correcting for duplicate and incorrect structures, which gave a final dataset of 540 compounds.

**S1.10.2 Docking workflow**

The goal of the workflow was to match each compound design to the corresponding covalent origin fragment and to include the binding pose information of the fragment into docking. The inspiration fragment was cited by the designer of the compound and a list of all designs and inspirations can be found on the Moonshot project GitHub.[36] Each design was matched with the corresponding fragment, the maximum common substructure (MCS) between them was identified, and the conformation of the design was aligned to the fragment before docking. The alignment was performed using a custom alignment script similar to the constrained alignment method in RDKit[53] to force the corresponding atom positions of the MCS into the same conformation, followed by a constrained energy minimisation, keeping the conformation of the MCS constant. Docking was performed using AutoDock4 (AD4), which considers ring conformations to be rigid when sampling ligand conformations before docking.[54] As a result, all rings present in the MCS are already aligned to the experimentally determined binding pose. Each design was docked to the corresponding $M^{pro}$ crystal structure of the origin fragment, after generation of the homodimer and charge optimization using Protonate3D in MOE.[21]

We used the FlexRes method in AD4 for covalent docking.[54] The covalent adduct of the COVID Moonshot design *after* reaction with the active site Cys-145 was selected as the flexible residue and a water molecule included as the "dummy" ligand. Docking and grid parameter files were generated for each Cys-145-inhibitor adduct individually with the rest of the corresponding co-crystallised dimeric $M^{pro}$ structure treated as the rigid receptor molecule. Docking with AD4 was performed using the Lamarckian Genetic Algorithm (LGA) and the following AD4 hyperparameters: population size 300; maximum number of energy evaluations 250 000; maximum number of generations 27 000; number of dockings 100.

The scoring function used by AD4[54] includes a pairwise evaluation of intermolecular interactions of the ligand and the protein, intramolecular interactions between residues of the protein and covalent adduct, and an estimation of the conformational entropy lost upon binding. For the evaluation of the covalent docking procedure, only the *intramolecular* terms are relevant, since they correspond to the changes in the energy of the flexible residues plus covalent adduct. Since AD4 automatically clusters docking results by the total estimated free energy of binding, covalent docking results must be re-clustered using the "Final Total Internal Energy" instead (as reported in the DLG docking log file). For clustering the docked poses of the covalent adducts, the same hierarchical clustering procedure as used in the native AD4 method is employed. A new cluster is seeded with the lowest energy pose, and all remaining poses within a threshold (< 2 Å RMSD) are added to that cluster. The procedure is repeated for the next lowest energy pose, until all docked poses have been assigned to a cluster. RMSD values between poses were calculated using the Open Drug Discovery Toolkit (ODDT),[55] to account for intramolecular symmetry, such as equivalent methyls in tertiary butyl groups.

Docked poses were compared to the original origin fragment crystal structure using SuCOS.[56] SuCOS produces normalised scores to a value between 0 and 1, where 1 indicates perfect overlap and identical molecules. Both the shape and pharmacophoric feature overlaps are weighted equally in the SuCOS score. Based on work by Leung *et al.*,[56] a SuCOS score of 0.55 between two molecules was found to be equivalent to a pose-pose RMSD of 2 Å.

Covalent docking for PF-07321332 was performed identically to the other covalent Moonshot designs with the exception that no pre-alignment of the ligand was performed prior to docking. Instead, a random conformation of the ligand was used to seed the docking process. The azanide nitrogen was assigned a negative charge prior to docking.

`

### S2. Supplementary Results – Substrate Binding and Recognition

#### QM/MM studies of proton transfer in the catalytic dyad

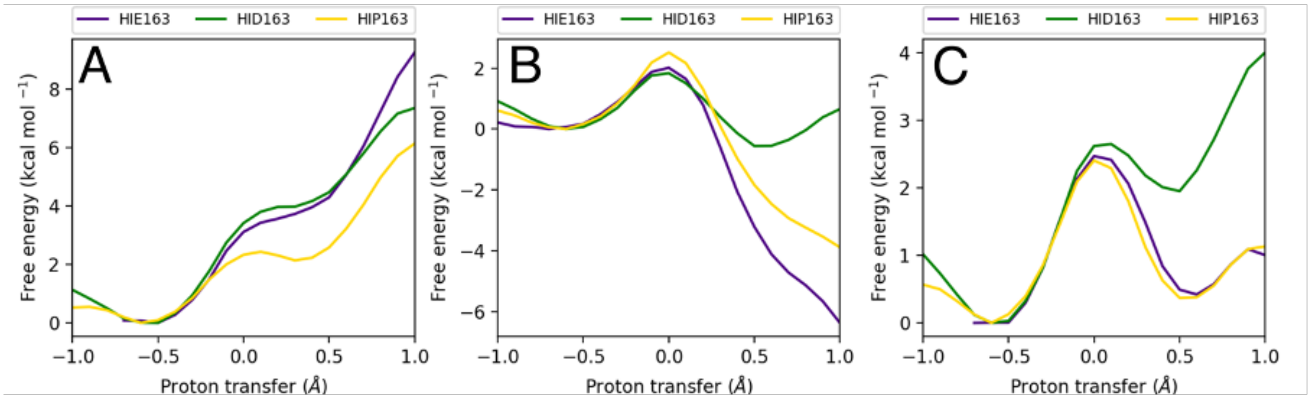

**Figure S2.1**: Free energy profiles of the proton transfer (PT) between Cys-145 and His-41 in s01-bound SARS-CoV-2 M$^{pro}$ with three different protonation states of His-163. (**A**) Free energy profiles from the neutral dyad. (**B**) Free energy profiles from the zwitterionic dyad. (**C**) Combined free energy profiles from the profiles in **A** and **B**. All free energy profiles were generated at the DFTB3/MM level of theory. A reaction coordinate value of -1.0 represents the neutral catalytic dyad and a value of 1.0 represents the zwitterionic catalytic dyad. HIE, HID and HIP refer to the εN, δN and doubly protonated models respectively in the Amber force field naming scheme.

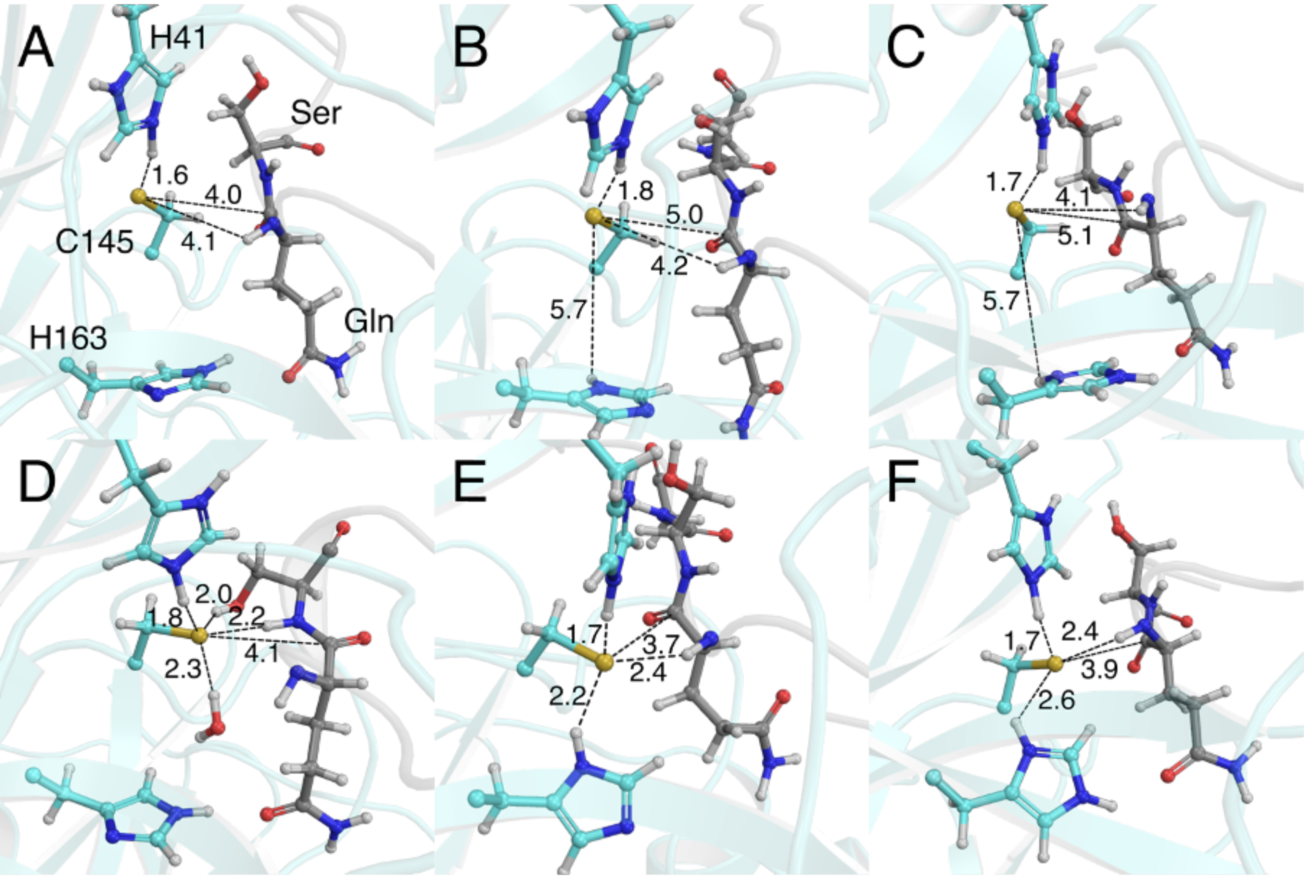

**Figure S2.2**: MD snapshots from umbrella sampling windows of the zwitterionic states of different free energy trajectories showing important interactions with the Cys-145 thiol(ate). (**A**) HIE-163, forwards. (**B**) HID-163, forwards. (**C**) HIP-163, forwards. (**D**) HIE-163, backwards. (**E**) HID-163, backwards. (**F**) HIP-163, backwards. Distances are in Angstroms.

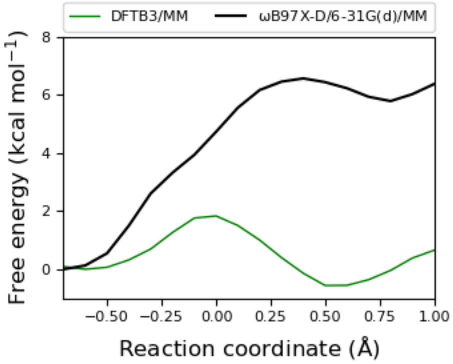

**Figure S2.3**: Free energy profiles of the catalytic dyad PT (simulated backwards) in the HID-163 system, at the DFTB3/MM and ωB97X-D/6-31G(d)/MM levels of theory.

`

#### Tautomeric and conformational states of other histidine residues

**Table S2.1**: SARS-CoV-2 M$^{pro}$ histidine protonation states adopted in this study, based on the apo crystal structure (PDB entry 6yb7).[15] Using standard nomenclature in the AMBER force fields, $\delta$-protonated His residues are denoted HID and $\varepsilon$-protonated His residues HIE.

| His | State | Reason |
|---|---|---|
| 41 | HID | Facilitates deprotonation of Cys-145 with the unprotonated $\varepsilon$-nitrogen. |
| 64 | HIE | Solvent exposed (HIE chosen as default). |
| 80 | HID | Allows $\delta$-NH to form hydrogen bond with Asn-63 side chain oxygen. |
| 163 | HIE | Allows $\varepsilon$-NH to form hydrogen bond with substrate P1 Gln side chain oxygen. In the apo structure, the sulfoxide oxygen of a DMSO molecule takes the place of this oxygen. |
| 164 | HIE | Uncertain based on inspection of crystal structure, but preliminary MD simulations (*vide infra*) suggested HIE to be more stable. |
| 172 | HIE | Allows $\varepsilon$-NH to form hydrogen bond with Glu-166 side chain oxygen. |
| 246 | HIE | Allows $\delta$-N to accept hydrogen bond from Thr-243 backbone NH. |

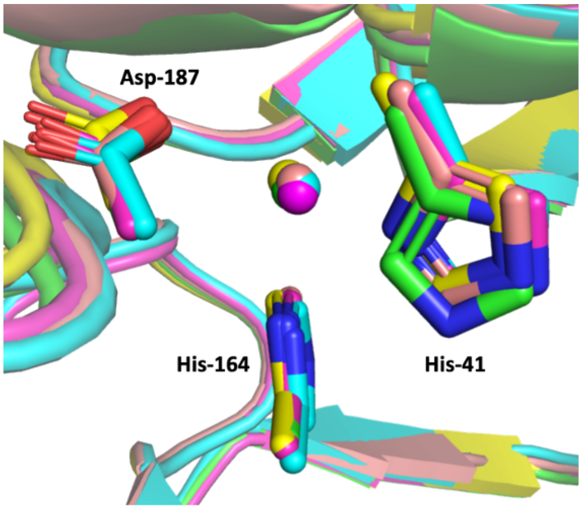

**Figure S2.4**: A conserved water molecule (shown as sphere in identical colour as the protein) is located between the sidechains of His-41, His-164 and Asp-187, in various M$^{pro}$ crystal structures (aligned using chain A), including PDB 6yb7 (HOH 644; green),[15] 6lu7 (HOH 445; cyan),[47] 7bqy (HOH 570; magenta),[47] 6y2g (HOH 560; yellow)[57] and 6wqf (HOH 417; salmon).[58] Note that His-41 in PDB entry 6yb7 has a unique conformational state.

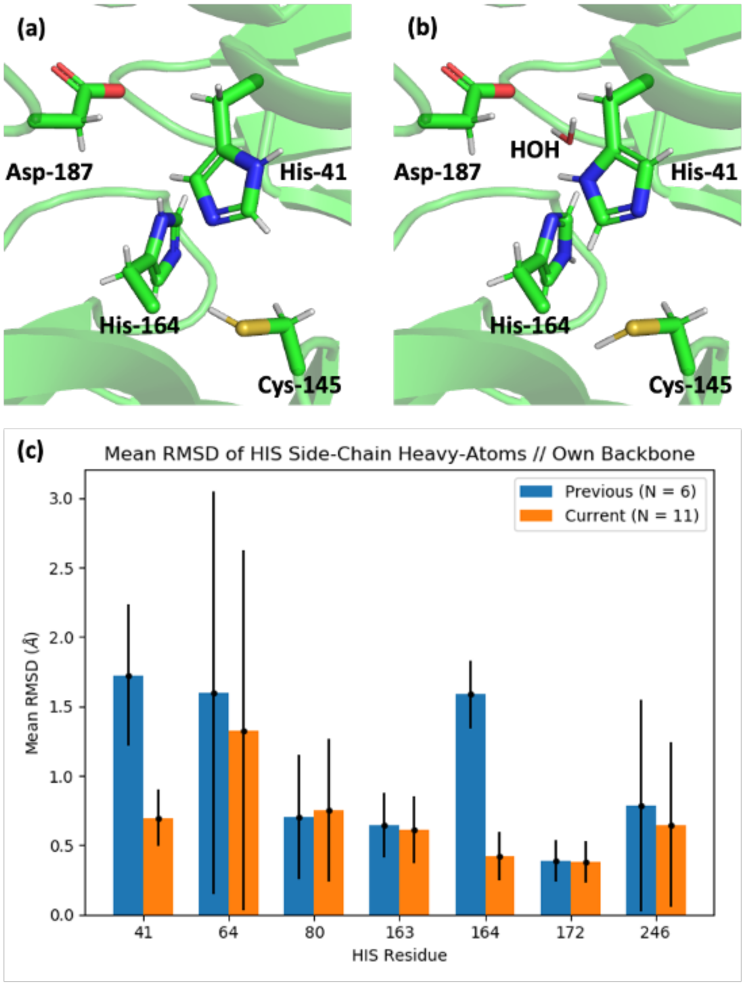

**Figure S2.5**: Comparison of the (a) previous and (b) current setups of His-41 and its surrounding residues for MD simulations (PDB 6yb7). The changes involve rotation of His-41, $\varepsilon$-protonation on His-164, and the retention of water HOH 644. Protons were added automatically by GROMACS (v. 2019.2). (c) The bar plot compares the mean RMSD of the side chain heavy atoms of each His residue over 100 ns MD relative to their starting positions, after fitting to its respective backbone (N, C$\alpha$, C) atoms. Error bar refers to mean standard deviation. A higher mean RMSD indicates greater deviation from the setup, while a larger error bar reflects higher flexibility during MD. The current setup resulted in lower and less fluctuating RMSD values for both His-41 and His-164. The previous set of simulations involved the M$^{pro}$ dimer in complex with truncated s01 substrates (Ac-AVLQSG-NMe) in both active sites, whereas the current simulations involved the 11 M$^{pro}$-substrate (s01-s11) complexes each with one active site occupied.

`

### Models of $M^{pro}$-substrate peptide complexes

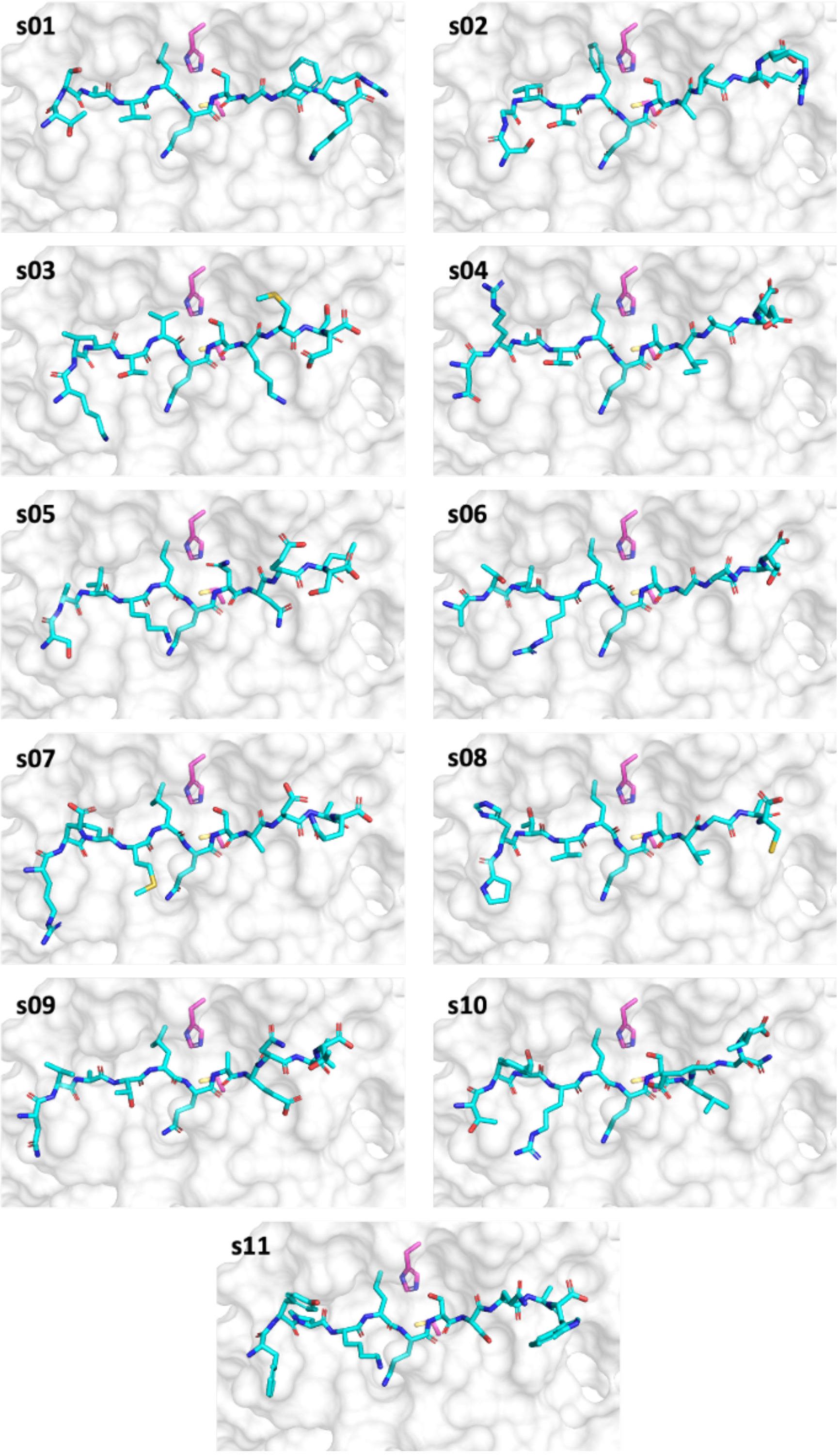

**Figure S2.6**: Starting conformations of the 11 native substrates (s01-s11; cyan) in complex with SARS-CoV-2 $M^{pro}$ (PDB entry 6yb7;[15] shown as a white surface with the dyad His-41 and Cys-145 in magenta), constructed by a comparative modelling approach. The crystal structure of the H41A SARS-CoV $M^{pro}$-s01 complex (PDB 2q6g) was used as a template.[1]

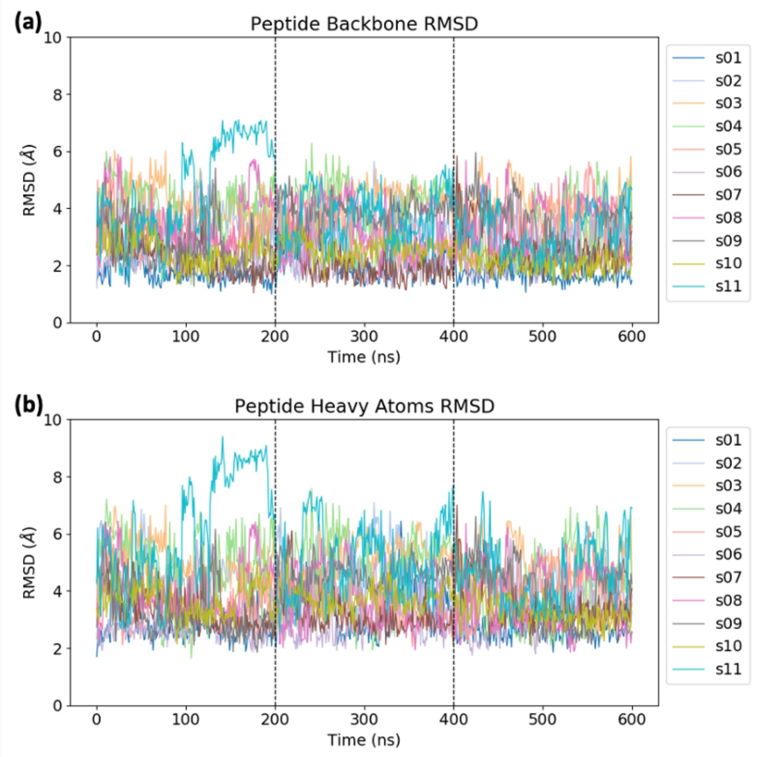

**Figure S2.7**: RMSD of (a) the peptide backbone (N, C$\alpha$, C) and (b) all peptide heavy atoms during the concatenated 3 × 200 ns explicitly-solvated MD simulations of the 11 $M^{pro}$-substrate complexes relative to the initial configuration, with trajectories fitted using the $M^{pro}$ dimer backbone.

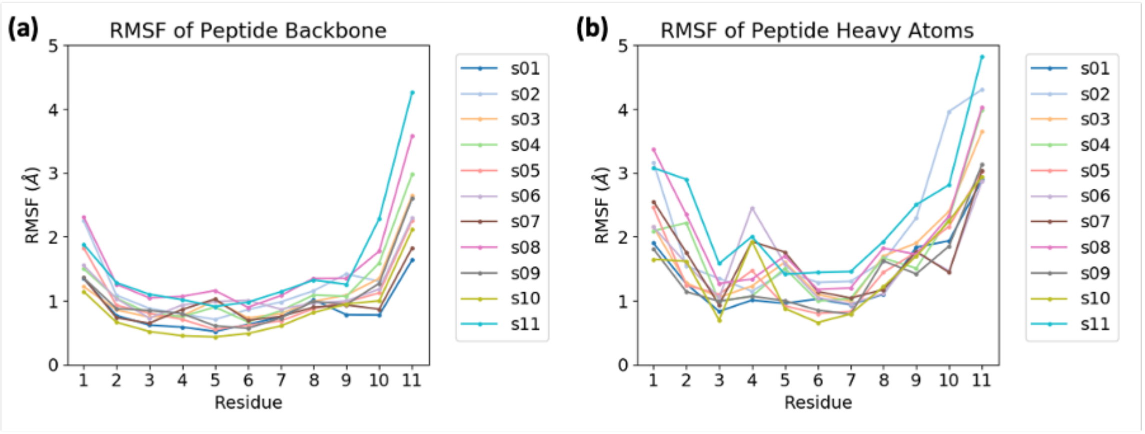

**Figure S2.8**: RMSF of (a) the peptide backbone (N, C$\alpha$, C) and (b) all peptide heavy atoms averaged per residue during the explicitly-solvated MD simulations of the 11 $M^{pro}$-substrate complexes.

`

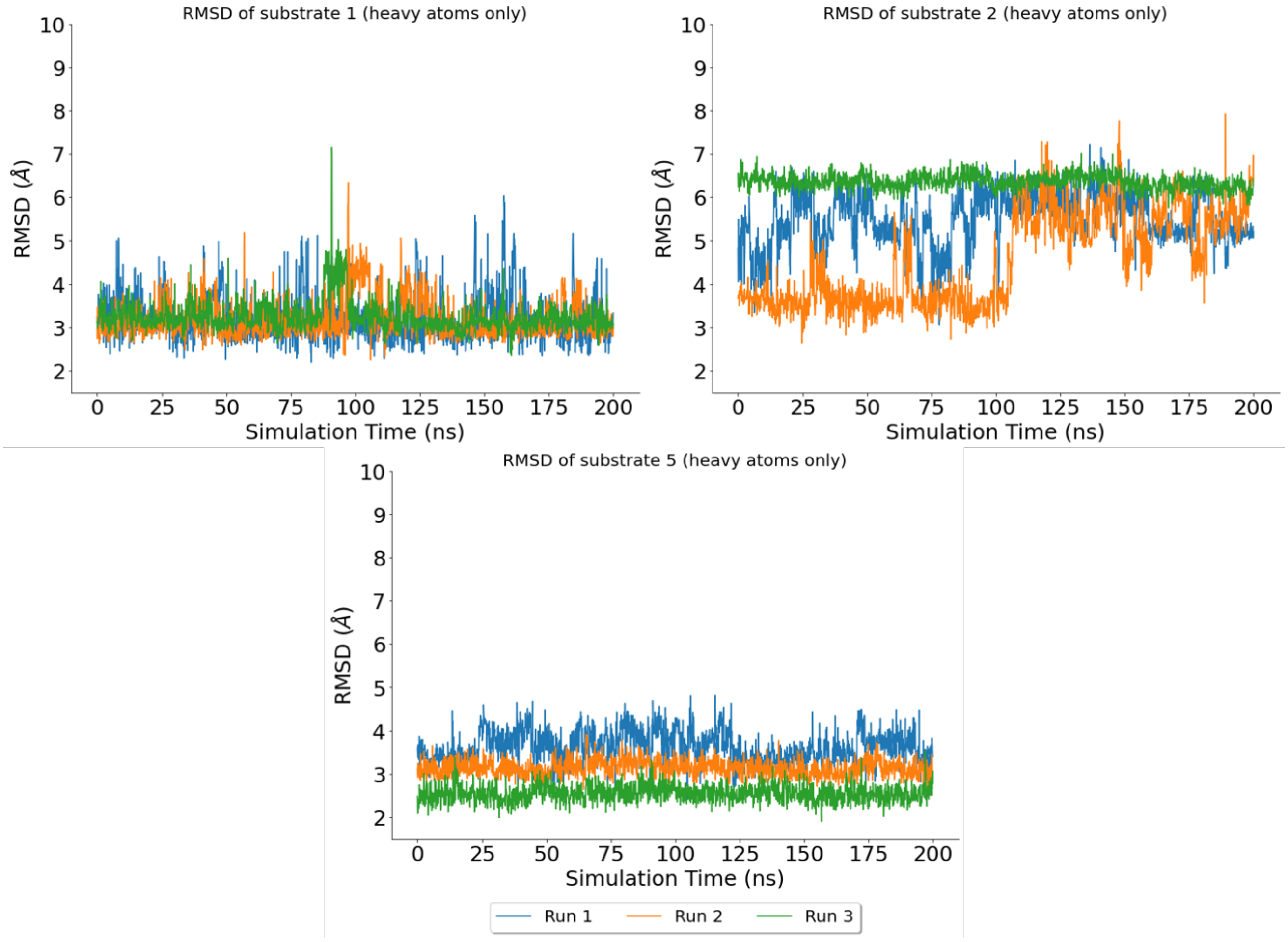

**Figure S2.9**: RMSD of substrate heavy atoms (i.e. not including hydrogens) across 3 × 200 ns MD simulation of iMD-VR docked structures, based on comparison to the starting structure (i.e. the docked structure from iMD-VR). Substrates s01 and s05 have a lower RMSD on average compared to s02. This could be due to the lack of HBs formed in s02 simulations, compared with s01 and s05 (**Figure S3.12**).

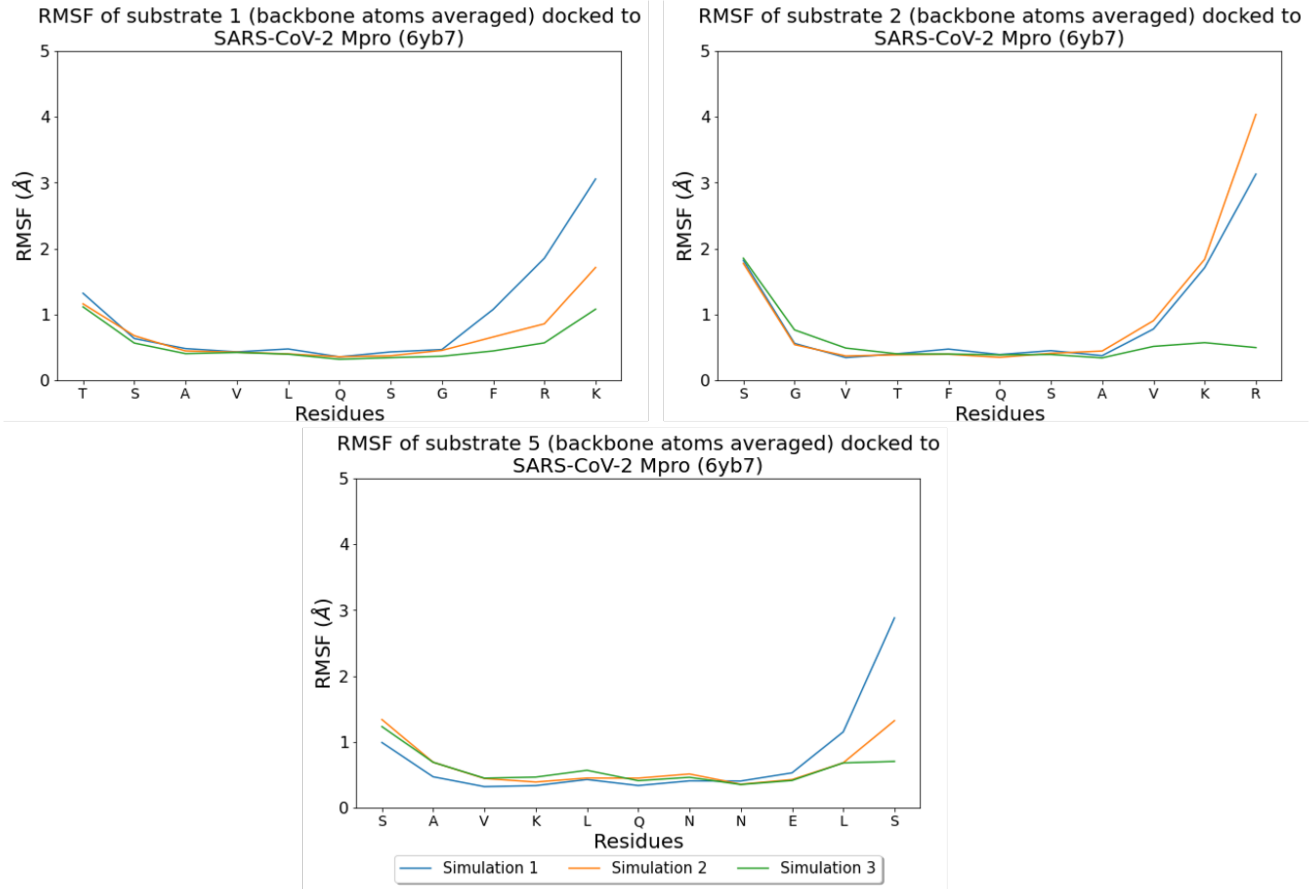

**Figure S2.10**: RMSF of substrate backbone atoms across 3 × 200 ns MD simulation of iMD-VR docked structures. In most cases, the P' (C-terminal) side of the substrates is more flexible than the P (N-terminal) side.

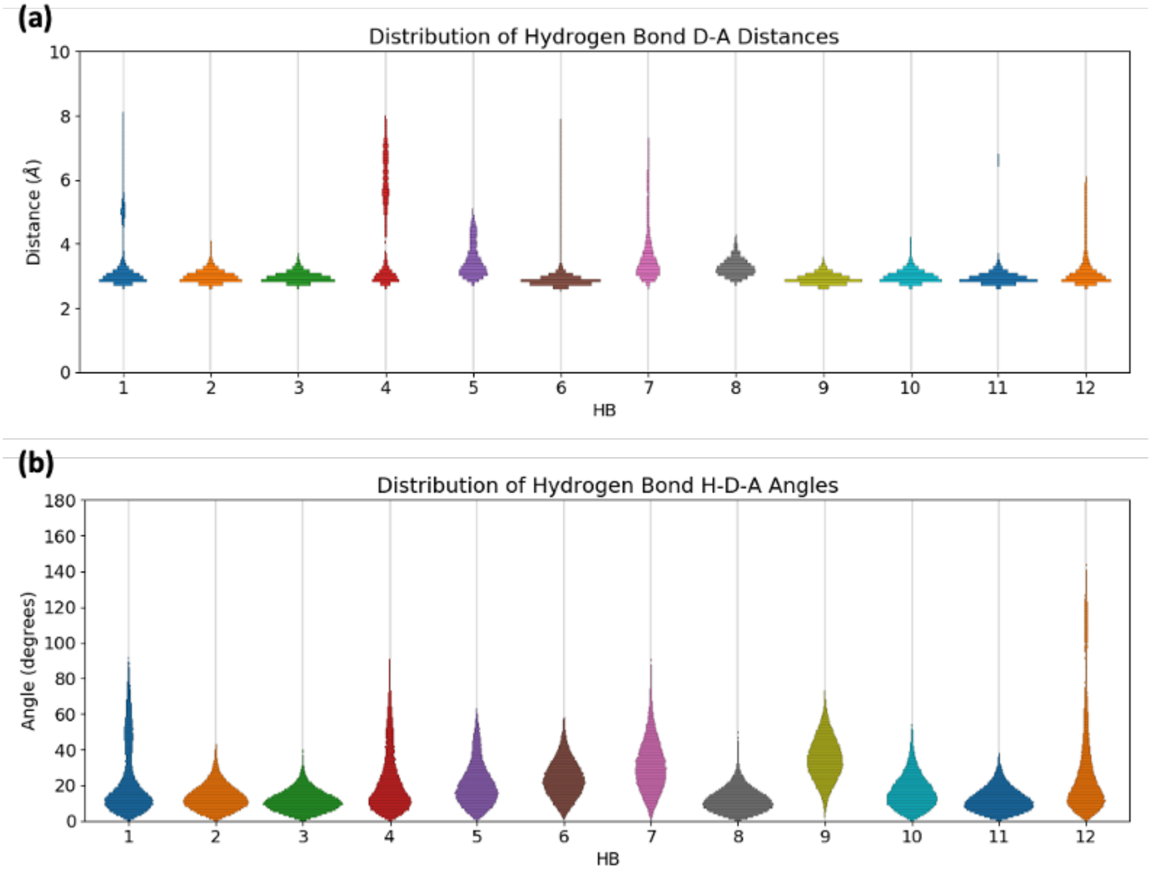

**Figure S2.11**: **Conserved hydrogen bond interactions**. Overall distributions of the (a) donor-acceptor distances ($d_{D\text{-}A}$) and (b) proton-donor-acceptor angles (∠(H-D-A)) corresponding to the 12 monitored $M^{pro}$-substrate hydrogen bonds (main text **Figure 3**), over the 3 × 200 ns explicitly-solvated MD simulations performed on each of the 11 $M^{pro}$-substrate complexes. A HB is defined using the combined criteria of $d_{D\text{-}A}$ ≤ 3.5 Å and ∠(H-D-A) ≤ 30°.

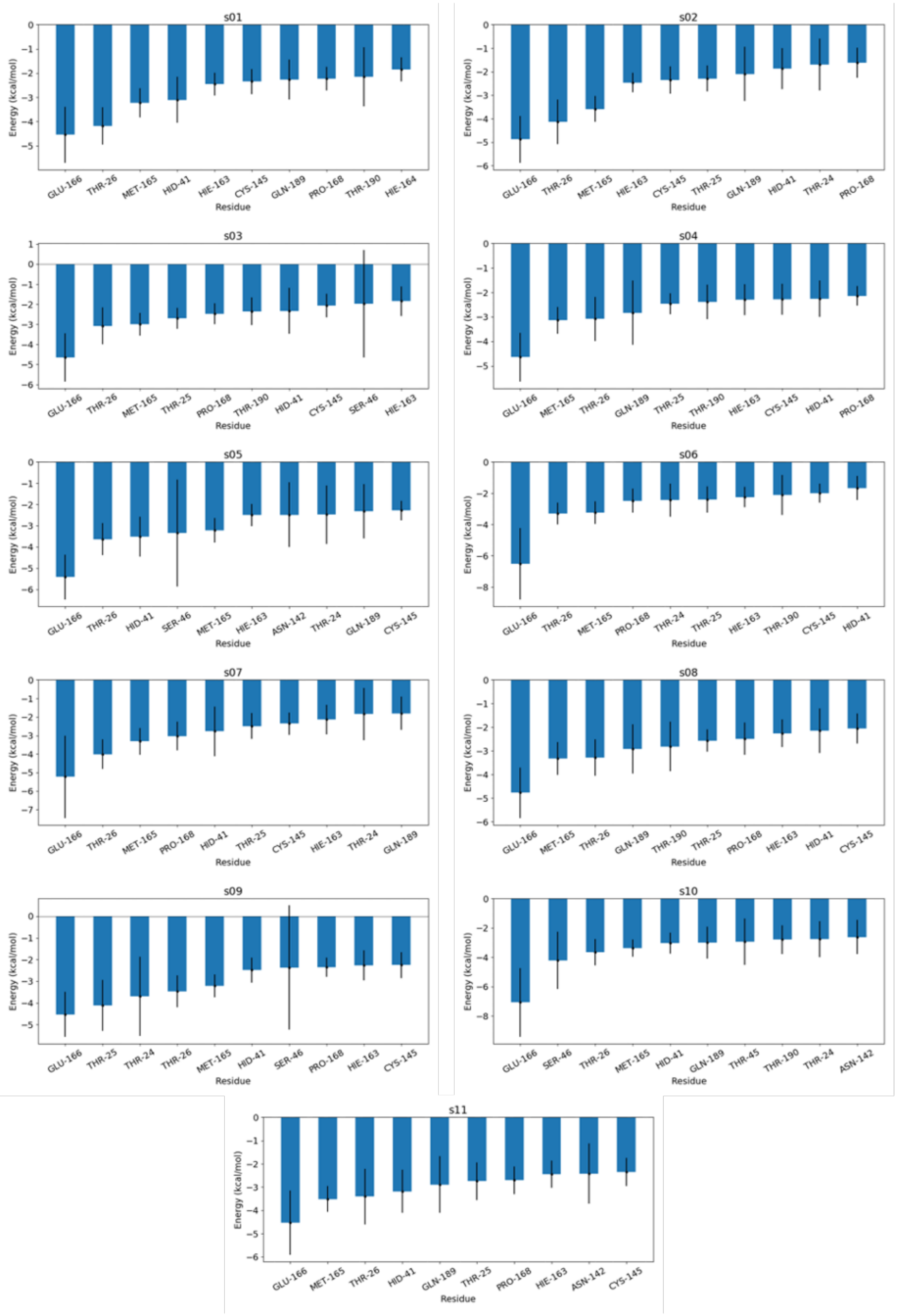

**Figure S2.12**: **MM-GBSA analysis.** The ten M^pro residues which contribute most to the MM-GBSA binding energy (error bar = standard error of mean over 120 frames) for each of the 11 M^pro-substrate complexes.

`

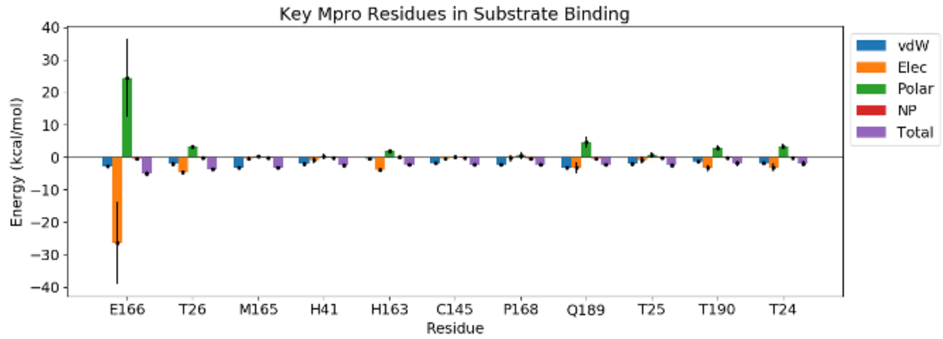

**Figure S2.13**: Contributions to the MM-GBSA binding energy by each hotspot residue (mean ± standard deviation across 11 systems), decomposed into the type of interactions: van der Waals (vdW), electrostatic (Elec), polar solvation from the generalised Born model (Polar), non-polar solvation from surface area calculation (NP), and total contribution. Given the varieties of charges and charged residue distributions across the substrates, there are relatively large variations in the electrostatic and polar solvation contributions by Glu-166, which is the only charged residue out of the identified conserved hotspot residues on M[pro].

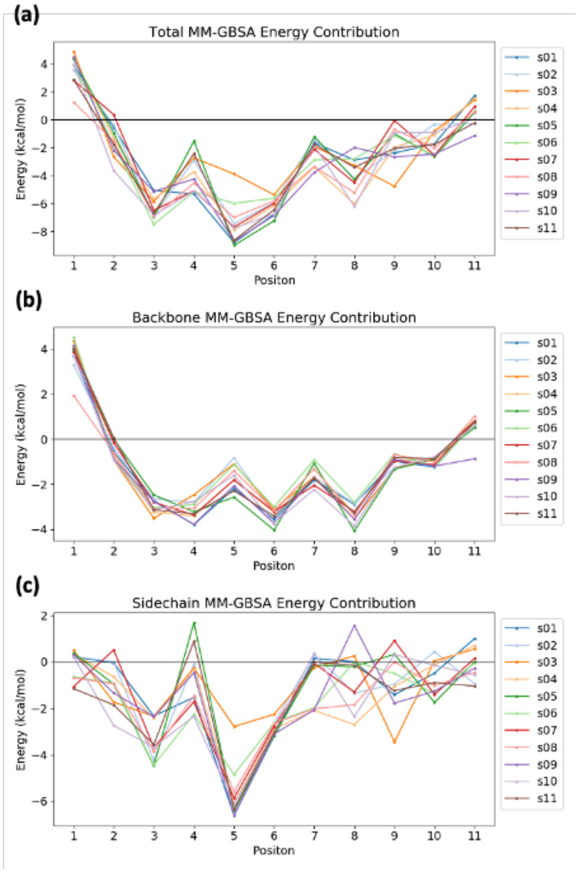

**Figure S2.14**: Contributions to the MM-GBSA binding energy for each residue of the substrate peptide, showing (a) the total contribution, as well as decomposition into (b) backbone and (c) sidechain contributions.

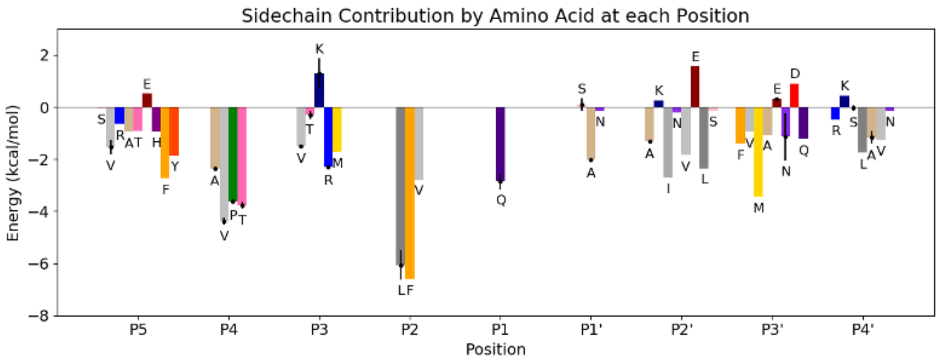

**Figure S2.15**: Average sidechain contributions to the MM-GBSA binding energy by each amino acid identity (except Gly) along the substrate peptide. Error bars (standard deviations across substrates) are shown only if the amino acid occurs in multiple sequences. Terminal residues are omitted due to their small contributions (**Figure S2.14**).

`

### Density functional theory analysis of the interaction network

#### Comparison of energetic panorama between DFT and force field

In what follows we will compare the energetic panorama that is provided by DFT and the force field (FF), and show that they appear correlated (which goes in favour of a reasonable MD configurational sampling). Such a comparison enables insight into the nature of the interaction between the substrate and the protein. Then, the interaction is analysed by post-processing QM results, which will enable insight into the possible interaction patterns (or "interaction signatures") that each of the substrates generates on the protein. In particular, the ability of treating the whole system by a consistent level of theory enables us to analyse both short-range and long-range interactions on equal footing.

Using this approach, energies were computed for the substrate (S), the protein dimer (P) and the total assembly (A) from which an interaction energy was calculated as follows: $E_{int} = E_A - (E_S + E_P)$; solvent molecules were omitted for easy comparisons. While absolute estimates of physical quantities are not the object of the analysis here (due to the omission of entropy and solvent contributions), this approach provides an unbiased first-principles approach, complementary to the one provided by classical FFs, to identify the main features involved in substrate-protein binding.

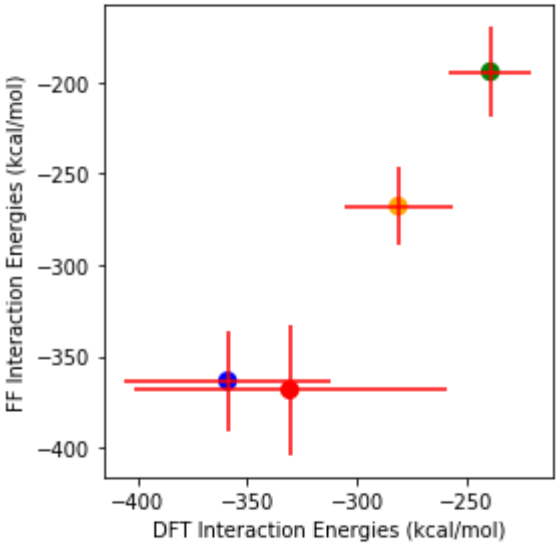

**Figure S2.16**: Comparison of the three-point interaction energies (kcal mol$^{-1}$) coming from full DFT calculation of the clustered MD snapshots. DFT interaction energies (x-axis) are represented together with FF three-point interaction energies (y-axis). We regroup the data on a per-peptide basis, for peptides s01 (blue), s02 (red), s05 (green), and p13 (orange). Statistic distributions are averaged taking into account the weights of the clusters in the trajectories.

The results (**Figure S2.16**) show a good correlation between the different approaches. The energetic panorama offered by the MD of the assemblies is likely to be similar to what would have been found by employing a first-principle approach. That goes in favour of a reasonable conformational sampling offered by the MD. Second, this constitutes an indication that charge-polarisation, which would be captured (at least partially) by DFT, does not play a major role for the peptide-enzyme interaction. This is related to the fact that the most charged peptides are ones which exhibit more attractive interactions. Within this assumption the QM interaction energy would then be efficiently approximated by the electrostatic peptide-enzyme interaction. Indeed, we show (**Figure S2.17**, left) that there is a remarkable correlation between the three-point interaction and such an approximated term (which is calculated by a multipole expansion of the DFT charge density result). To validate this, we performed the same analysis by employing the polarizable FF Polaris(MD),[59] the results of which show almost perfect correlation between the two FFs (**Figure S2.17**, right).

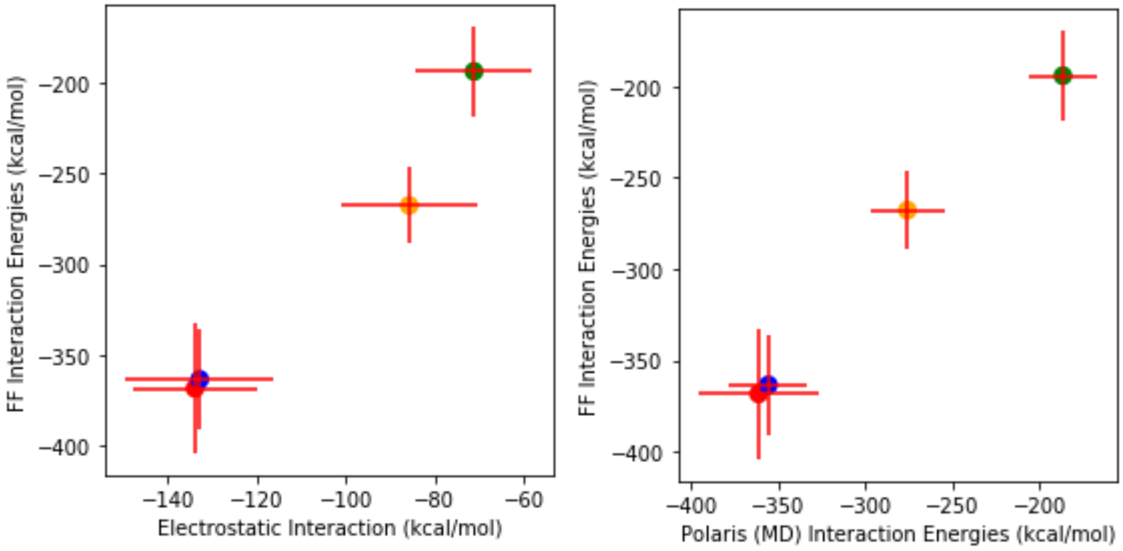

**Figure S2.17**: Comparison of the FF three-point interaction energies (y-axis) in the clustered MD snapshots, and on the x-axis (left) the electrostatic peptide-enzyme interaction or (right) interaction energy calculated with Polaris(MD). Each data point represents a peptide sequence: s01 (blue), s02 (red), s05 (green), and p13 (orange). Statistic distributions are averaged taking into account the weights of the clusters in the trajectories.

Note that the contribution to the interaction energy that is provided by vdW terms (the semi-empirical D3 dispersion term) is approximately constant (around 100 kcal $mol^{-1}$), and therefore the trends for the various interactions are provided by considering only the DFT-PBE contributions.

**Contact map derived from electronic structure**

A quantity which is of particular interest in determining the interaction network is the system's Hamiltonian, which gives a measurable indication of the strength of the chemical bonding among the electron clouds of each of the fragments. The partial traces of this operator, once projected on the system's fragments, provide an indication of the fragments which mostly participate in the chemical bondings during the trajectories. Such "contact interaction energy", $E_{cont}$, can be interpreted as short-range sharing of electrons between two fragment residues. For each system, we have plotted this term for the ten most stabilising interactions (**Figure S2.18**).

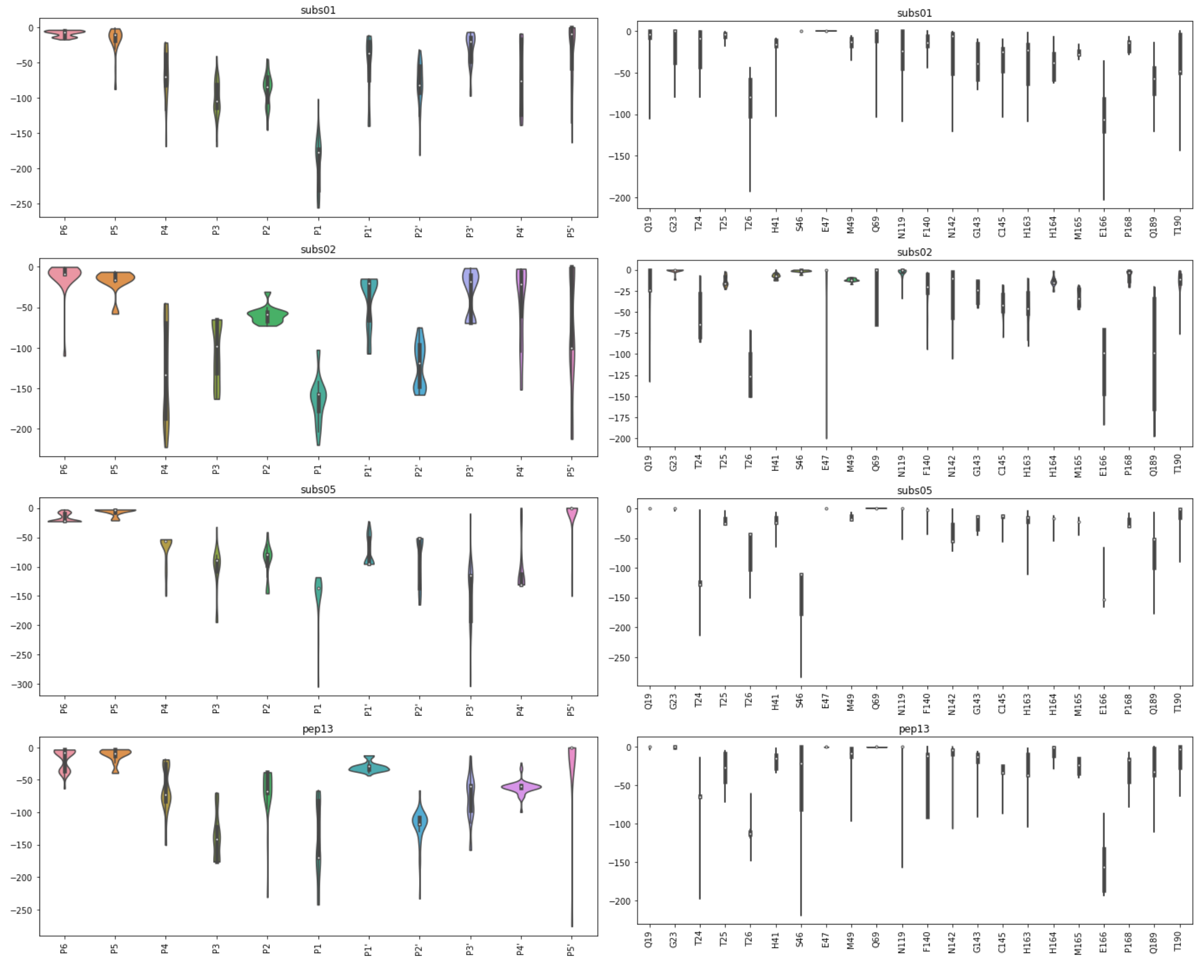

**Figure S2.18**: Violin plot distributions of contact interaction energies (kcal mol$^{-1}$) between substrate/designed peptides (s01, s02, s05, p13) and selected M$^{pro}$ residues (which rank in the top ten in terms of stabilising interactions with the peptide in at least one of the systems), displayed (left) on the peptide or (right) on the enzyme residues.

#### Long-range electrostatic interaction patterns

The QM-FF comparison above (**Figure S2.17**, left) suggests that peptide binding trends can be estimated, at first approximation, by only considering the long-range electrostatic interactions. As we have employed a full QM calculation on the entire system it is interesting to show which are the emerging patterns of these interactions for the different peptides. This is helpful in defining other "interaction signatures" which are based on long-range patterns. Below (**Figure S2.19**) we show how such interactions behave during the dynamics. For clarity, we only represent residues whose magnitude of the (MD-averaged) interaction is larger than 7 kcal mol$^{-1}$ for at least one of the systems. Some residues belonging to the other M$^{pro}$ monomer (chain B) also appear to be relevant, due to their relative geographical proximity with the substrates. Obviously, the main overall pattern is dictated by the total charge of the bound peptide, which disfavours the neutral s05 in comparison with the other positively charged peptides (s01, s02, p13). The strong binding role of Glu-166 is clearly visible.

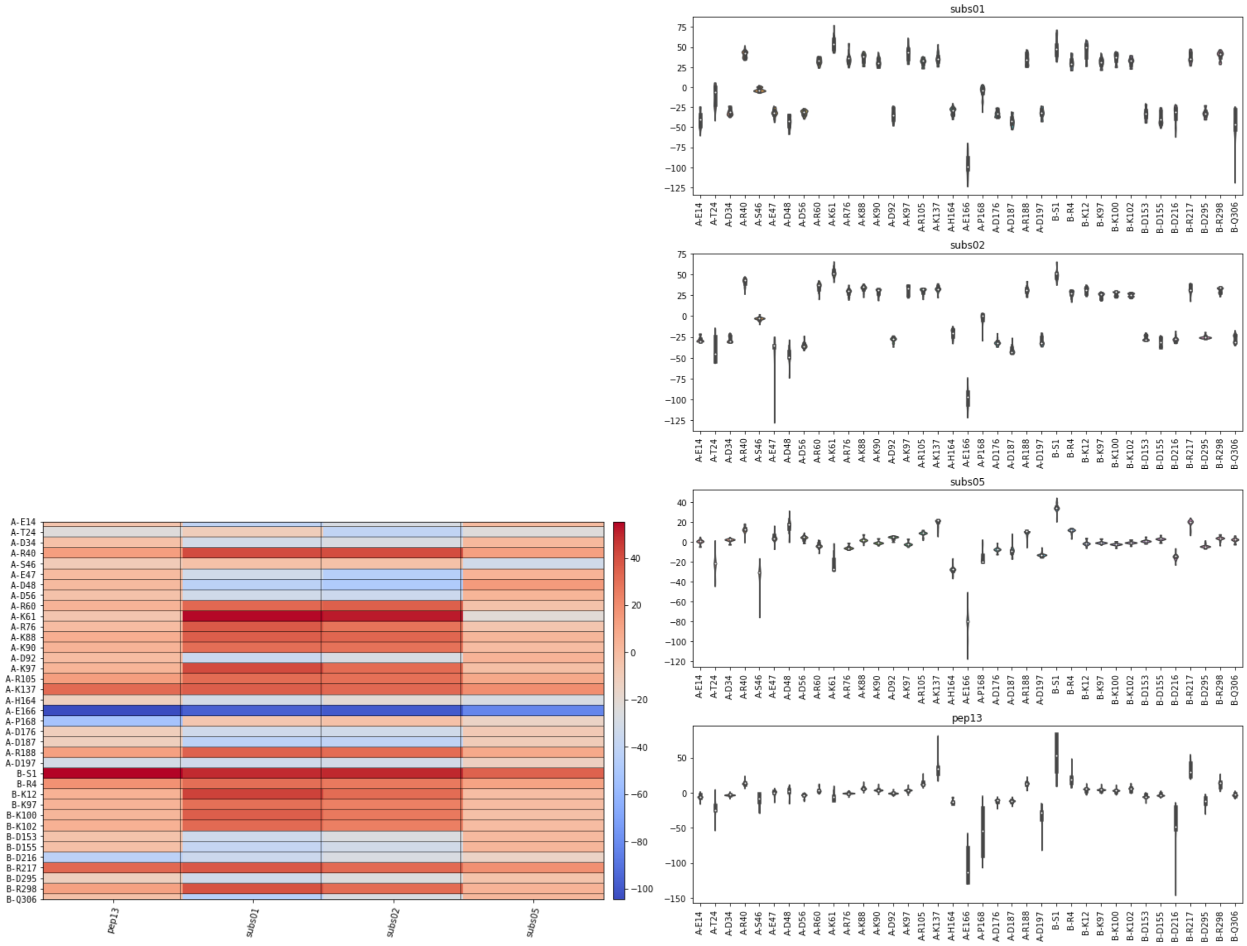

**Figure S2.19**: Distribution of long-range electrostatic interaction energies (kcal mol$^{-1}$) between substrate/designed peptides (s01, s02, s05, p13) and selected M$^{pro}$ residues (which show a >7 kcal mol$^{-1}$ interaction with the peptide in at least one of the systems), displayed as (left) a heatmap and (right) a box plot.

`

**Monitoring of substrate sequence hydrolysis by mass spectrometry**

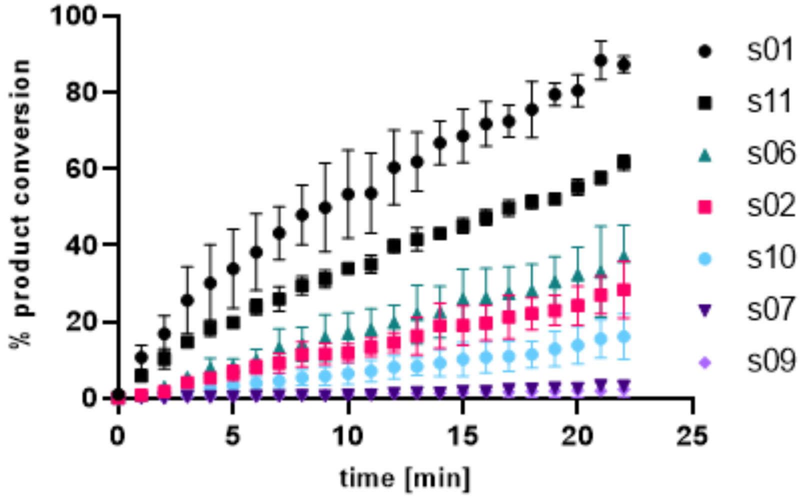

**Figure S2.20**: Ranking of catalysis of native M$^{pro}$ substrate peptides s01-s11 using denaturing MS conditions. No evidence for cleavage was observed for s03, s04, s05 and s08 under these conditions. Conditions: 0.15 µM M$^{pro}$, 2 µM substrate peptide in 20 mM HEPES, pH 7.5, 50 mM NaCl.

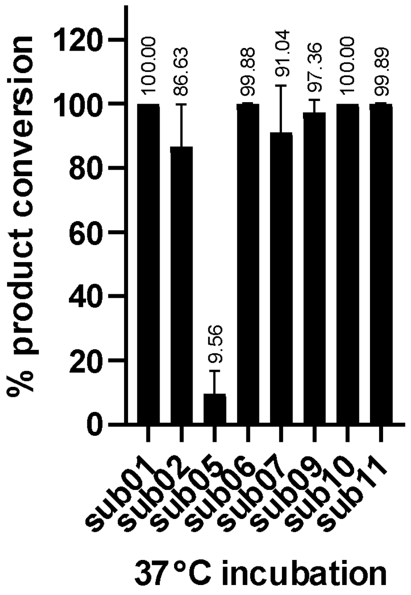

**Figure S2.21**: **Prolonged treatment of substrate peptides with M$^{pro}$ at 37°C**. Peak observed corresponded to the +1 charge state of the N-terminal cleaved product of s05; Of the analysed substrates, this was the only product not detected at room temperature [20° C] studies (**Figure S2.20**). Depletion of s04 was also observed but the apparently hydrophilic products were not observed potentially due to weak retention by the SPE C4 cartridge. Conditions: 0.15 µM M$^{pro}$, 2 µM substrate peptide in 20 mM HEPES, pH 7.5, 50 mM NaCl at 37°C, 300 rpm.

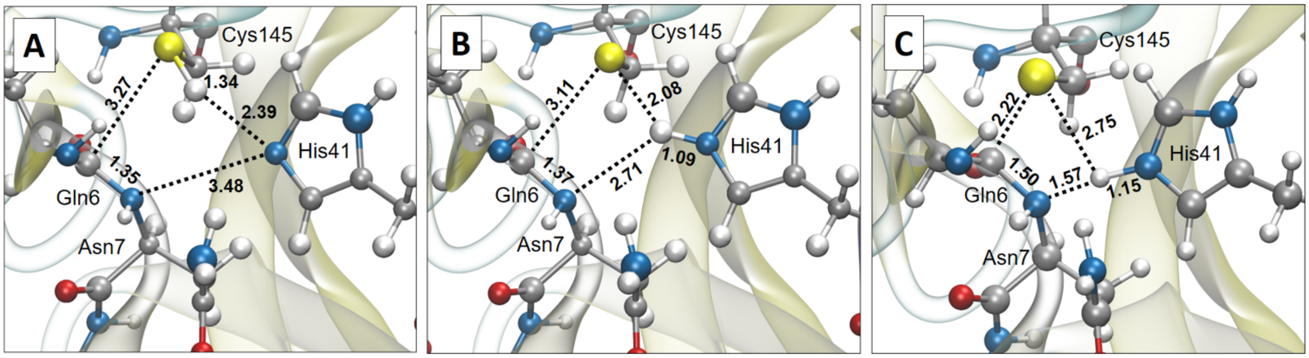

**Figure S2.22**: M06-2X(6-31+G(d,p))/MM structures of the neutral dyad reactants state (A), the pseudo-stable ion pair dyad state (B) and the transition state of the proteolysis (C) that connects the neutral dyad with the covalent bond adduct formed between the protein and the substrate. Structure of the transition state was fully optimized and characterized (imaginary frequency = 349.985i cm-1) from coordinates of the most populated clusters of peptibitor p13, including all water molecules (and counter ions). The structures of the ion-pair dyad and the neutral dyad were localized along the IRC path computed at the same level of theory from the optimized transition state. Importantly, structure of the ion pair dyad does not appear as a minimum in the IRC path, but as a shoulder. Animation of the full IRC path is reported as a mp4 file (attached). All distances are reported in Ångstroms. The QM sub-set of atoms include the side chain of His-41 (link atom between Cα and Cβ), Cys-145 together with carboxyl group of Ser-144 and part of Gly-146 (link atoms between C and Cα in both of them) and part of the substrate that includes full Gln-6 and Asn-7 together with carbonyl group of Trp-5 and Ser-8 (link atoms between C and Cα in both of them), which were treated at M06-2X(6-31+G(d,p)) with Gaussian 09.[14] The rest of the protein, solvent water molecules, and counterions were described by AMBER and TIP3P force fields,[60] as implemented in the fDynamo library.[61,62]

### S3. Supplementary Results – Peptide Inhibitor Design

#### Experimental verification of the designed peptides

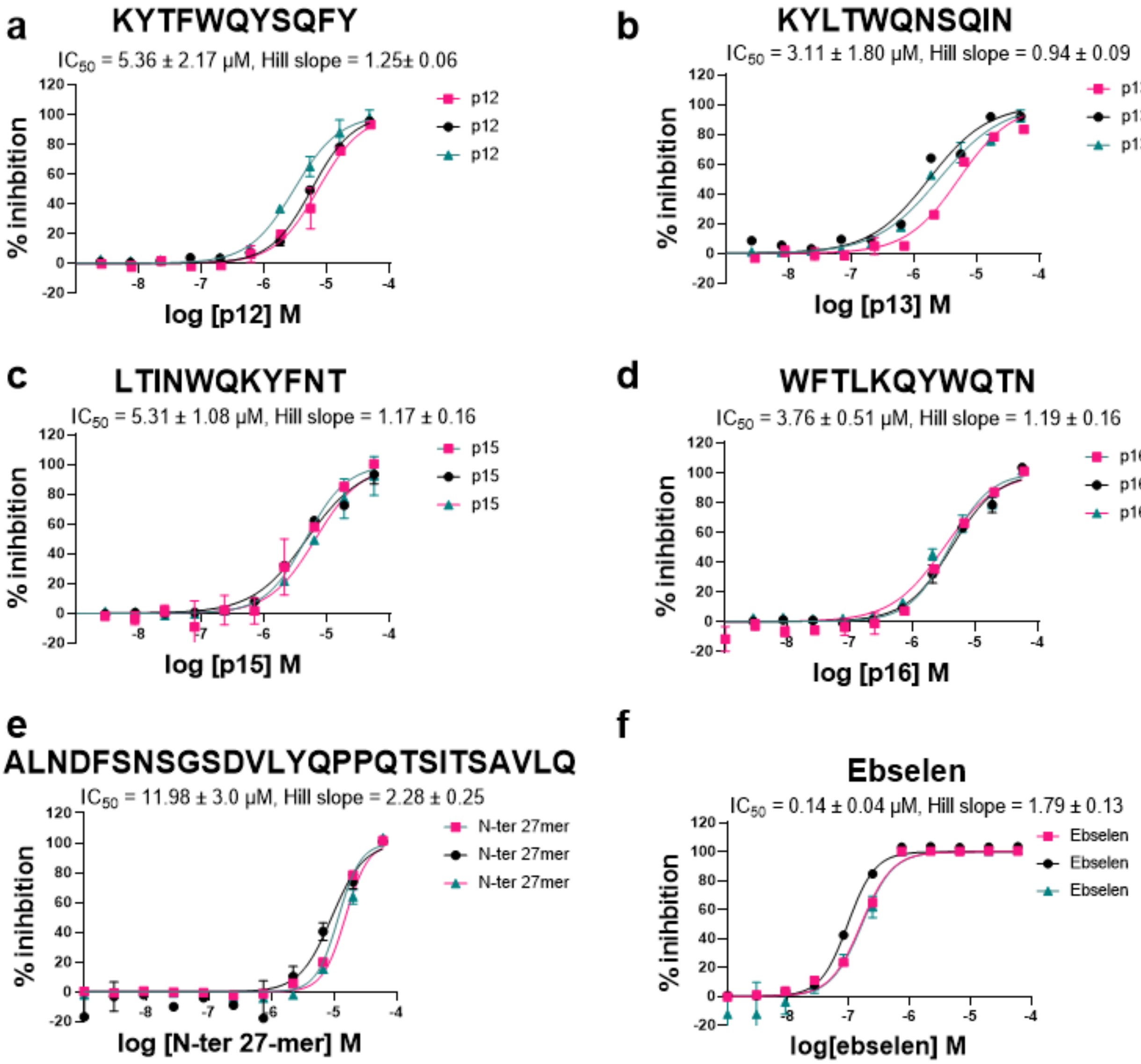

**Figure S3.1:** Designed peptides inhibit $M^{pro}$. $IC_{50}$s for a) p12, b) p13, c) p15, d) p16, e) 27-mer N-terminal cleavage product of s01 and f) ebselen. Reported $IC_{50}$s are means of independent repeats each composed of technical duplicates (n= 3 ± SD). Note: See **Experimental Section S1.8** for assay details. One of the independent repeats of p13 (black) was only comprised of a single dataset.

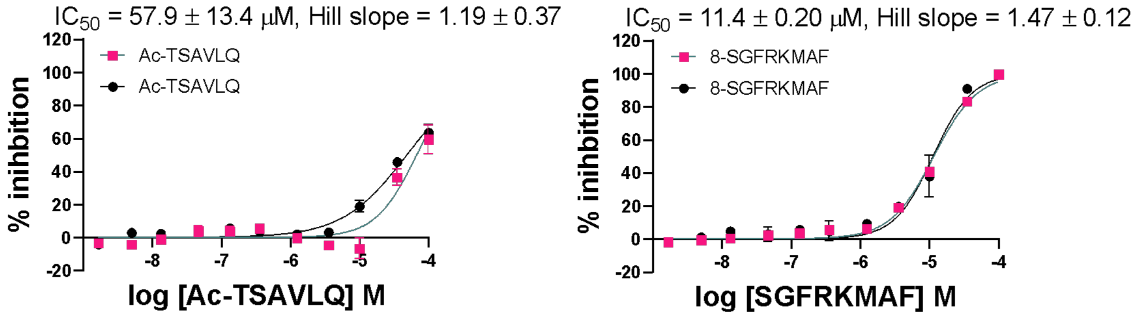

**Figure S3.2**: Acetylated N-terminal and extended C-terminal cleaved products of s01 inhibit M$^{pro}$ in a dose dependent manner. Conditions: 0.15 µM M$^{pro}$, 2 µM substrate peptide (s01: TSAVLQ/SGFRK-$NH_2$), 2 µM inhibitory peptides in 20 mM HEPES, pH 7.5, 50 mM NaCl.

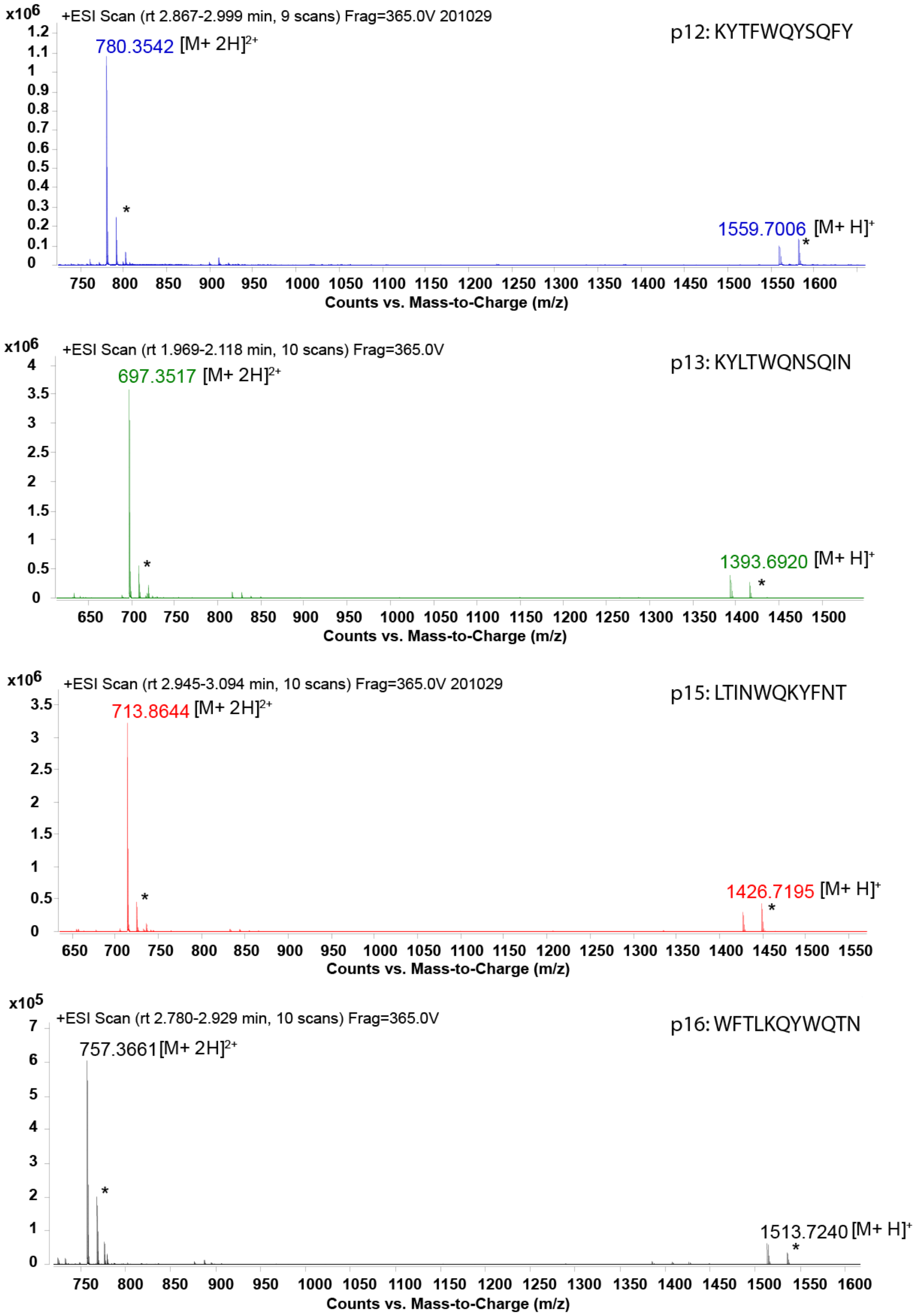

**Figure S3.3**: Potential cleavage products for the designed peptides (p12, p13, p15, p16) (2 µM) were not observed after overnight incubation with M$^{pro}$ (0.15 µM) at ambient temperature. With positive mode ionisation, +2 and +1 charge states for all intact peptides were observed, with the former being the predominantly observed charge state. Sodiated adducts $[M+H+Na]^{+}$ and $[M+2H+Na]^{2+}$ for all peptides were observed; indicated above with an asterisk. See **Experimental Section S1.8** for details.

**Table S3.1**: The method of Wei et al. was used to investigate the mode of inhibition.[63] This approach involves determining the $IC_{50}$ ratio (R) at two substrate concentrations (S1 and S2). The value of R is then used to assign the compound a (likely) competitive ($R_1$), non-competitive ($R_2$), uncompetitive ($R_3$) or mixed ($R_4$) inhibition mode. Using S1 = 2 µM and S2 = 40 µM (highlighted with a rectangle), the calculated R value for competitive inhibition ($R_1$) is 3.38. For the designed peptides (p12, p13, p15, p16), the results suggest competitive mode of inhibition with respective R values of (3.61), (6.02), (4.56) and (4.85).

| Mode of inhibition | | | Competitive | Non-competitive | Uncompetitive | | Mixed |
|---|---|---|---|---|---|---|---|
| S1 | S2 | $K_M$ | $R_1$ | $R_2$ | $R_3$ | α | $R_4$ |
| 2 | 10 | 14 | 1.50 | 0.30 | 1.00 | 1.00 | 1.00 |
| 2 | 20 | 14 | 2.13 | 0.21 | 1.00 | 2.00 | 1.33 |
| 2 | 25 | 14 | 2.44 | 0.20 | 1.00 | 3.00 | 1.60 |
| 2 | 30 | 14 | 2.75 | 0.18 | 1.00 | 4.00 | 1.85 |
| 2 | 35 | 14 | 3.06 | 0.18 | 1.00 | 5.00 | 2.10 |
| **2** | **40** | **14** | **3.38** | **0.17** | **1.00** | **6.00** | **2.34** |
| 2 | 45 | 14 | 3.69 | 0.16 | 1.00 | 7.00 | 2.58 |
| 2 | 50 | 14 | 4.00 | 0.16 | 1.00 | 8.00 | 2.81 |
| 2 | 55 | 14 | 4.31 | 0.16 | 1.00 | 9.00 | 3.05 |
| 2 | 60 | 14 | 4.63 | 0.15 | 1.00 | 10.00 | 3.28 |
| 2 | 65 | 14 | 4.94 | 0.15 | 1.00 | 11.00 | 3.52 |
| 2 | 70 | 14 | 5.25 | 0.15 | 1.00 | 12.00 | 3.75 |
| 2 | 75 | 14 | 5.56 | 0.15 | 1.00 | 13.00 | 3.98 |
| 2 | 80 | 14 | 5.88 | 0.15 | 1.00 | 600.00 | 5.82 |

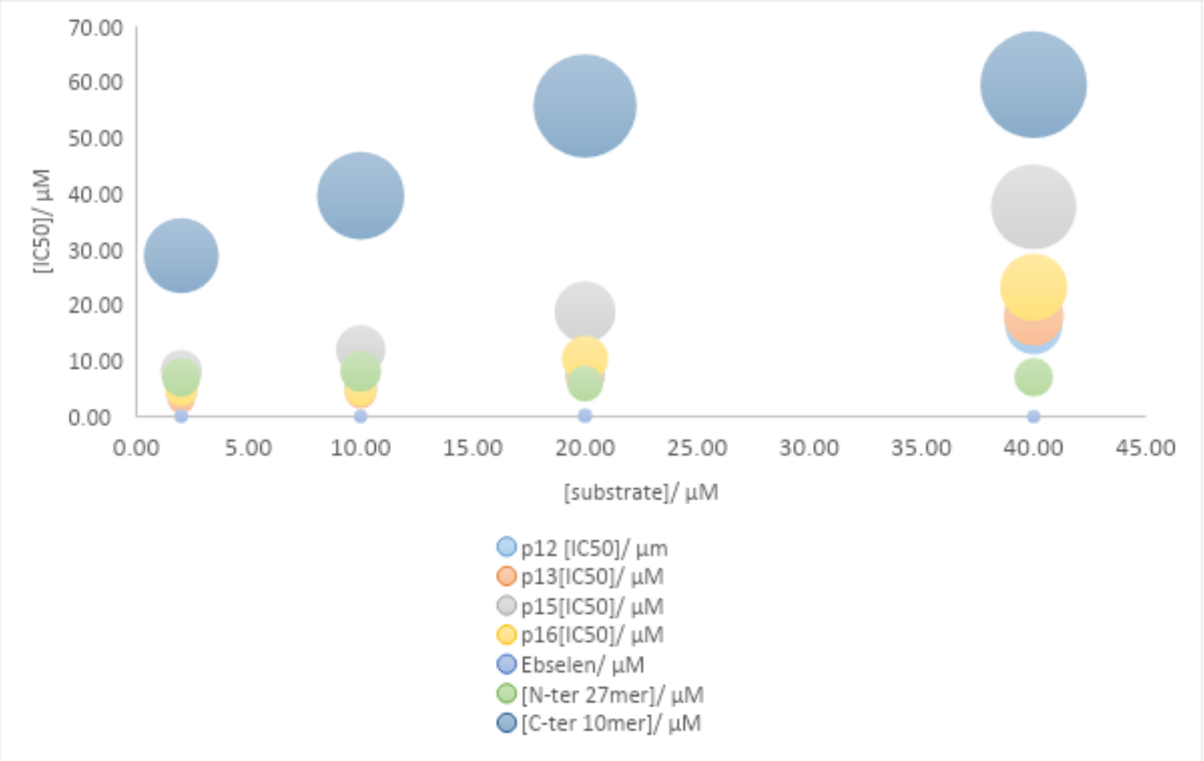

**Figure S3.4**: A linear increase in calculated $IC_{50}$s was observed in dose response studies with increasing substrate concentrations (2 µM, 10 µM, 20 µM and 40 µM). $IC_{50}$s are reported in **Table S3.2**.

`

**Table S3.2**: $IC_{50}$s of designed peptides at varying substrate (s01: TSAVLQ↓SGFRK-$NH_2$) concentrations of 2 µM, 10 µM, 20 µM and 40 µM.

| [S]/ µM | p12 [$IC_{50}$]/ µM | p13[$IC_{50}$]/ µM | p15[$IC_{50}$]/ µM | p16[$IC_{50}$]/ µM | Ebselen/ µM | [N-ter 27mer]/ µM | [C-ter 10mer]/ µM |
|---|---|---|---|---|---|---|---|
| 2.00 | 4.52 | 3.00 | 8.26 | 4.78 | 0.09 | 7.04 | 28.86 |
| 10.00 | 4.99 | 4.22 | 11.98 | 4.77 | 0.07 | 8.11 | 39.62 |
| 20.00 | 7.49 | 6.78 | 18.78 | 10.38 | 0.19 | 5.91 | 55.69 |
| 40.00 | 16.29 | 18.09 | 37.64 | 23.19 | 0.03 | 7.07 | 59.50 |

### Explicitly-solvated MD and implicit solvent iMD-VR

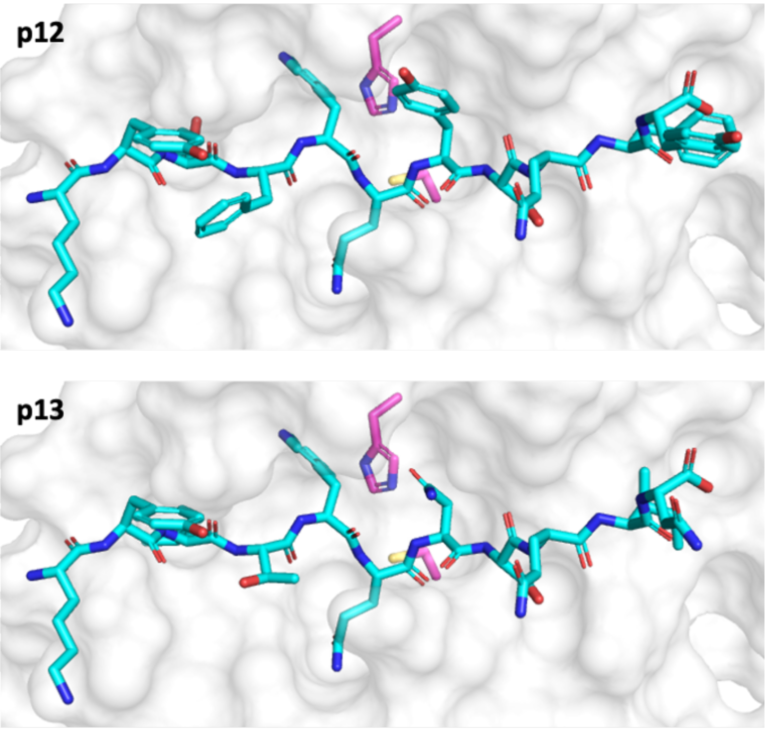

**Figure S3.5**: Starting configurations of the designed peptides p12 and p13 (cyan) in complex with SARS-CoV-2 M$^{pro}$ (PDB entry 6yb7;[15] white surface with the catalytic dyad residues His-41 and Cys-145 in magenta), constructed by a comparative modelling approach based on the starting configuration of the M$^{pro}$-s02 complex.

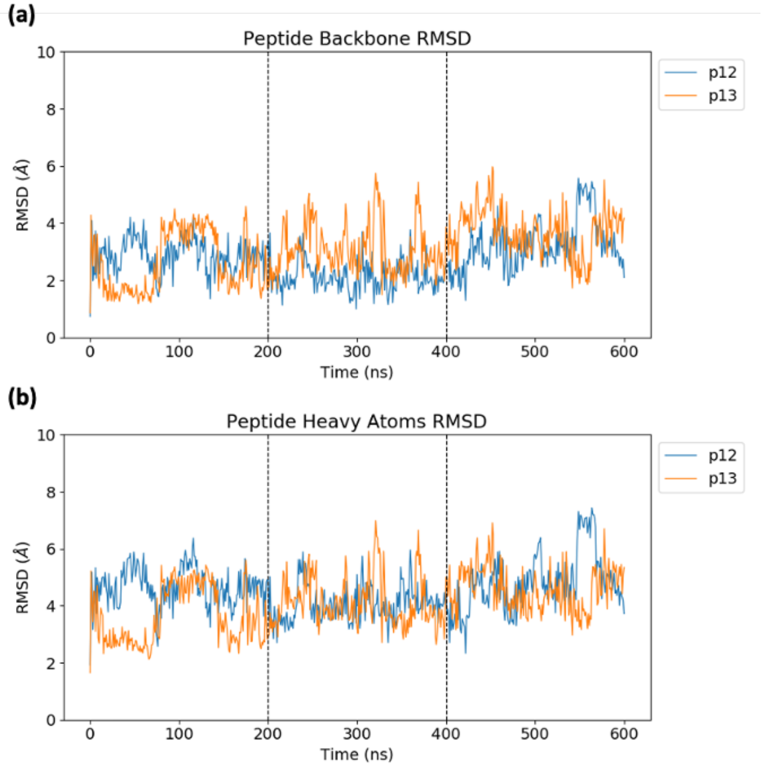

**Figure S3.6**: RMSD of (a) the peptide backbone (N, C$\alpha$, C) and (b) all peptide heavy atoms during the concatenated 3 × 200 ns explicitly-solvated MD simulations of M$^{pro}$ in complex with p12 and p13 relative to the initial configuration, with trajectories fitted using the M$^{pro}$ dimer backbone.

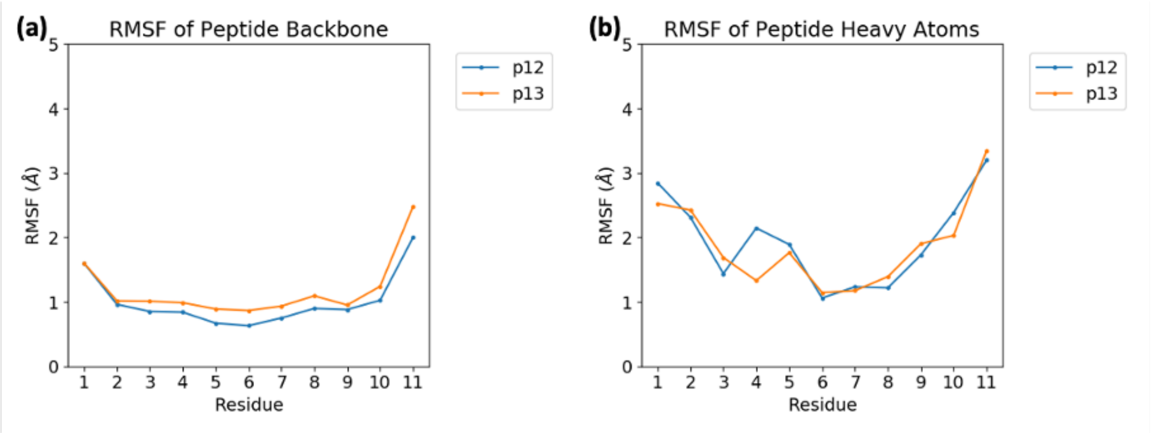

**Figure S3.7**: RMSF of (a) the peptide backbone (N, C$\alpha$, C) and (b) all peptide heavy atoms averaged per residue during the explicitly-solvated MD simulations of M[pro] in complex with p12 and p13.

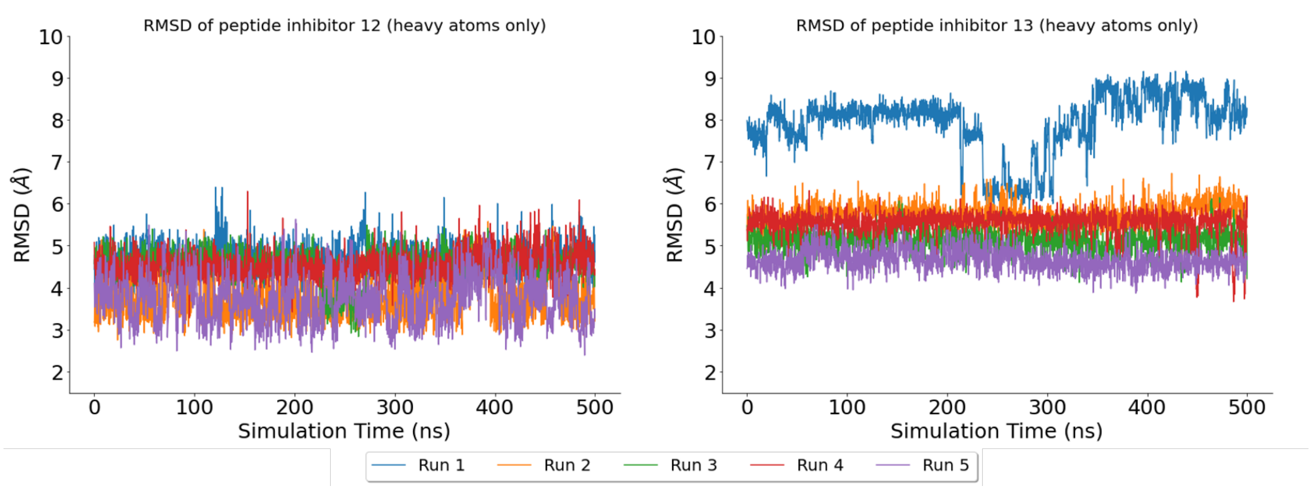

**Figure S3.8**: RMSD of p12 and p13 heavy atoms (i.e. not including hydrogens) across 5 × 500 ns MD simulations of iMD-VR docked structures, in comparison with the starting structure (i.e. the docked structure). The RMSD of p12 stays around 2.5–5.5 Å across all simulations, indicating that a common, stable bound structure has been found. On the other hand, the RMSD of p13 is higher, with values ranging 4.5–8.5 Å.

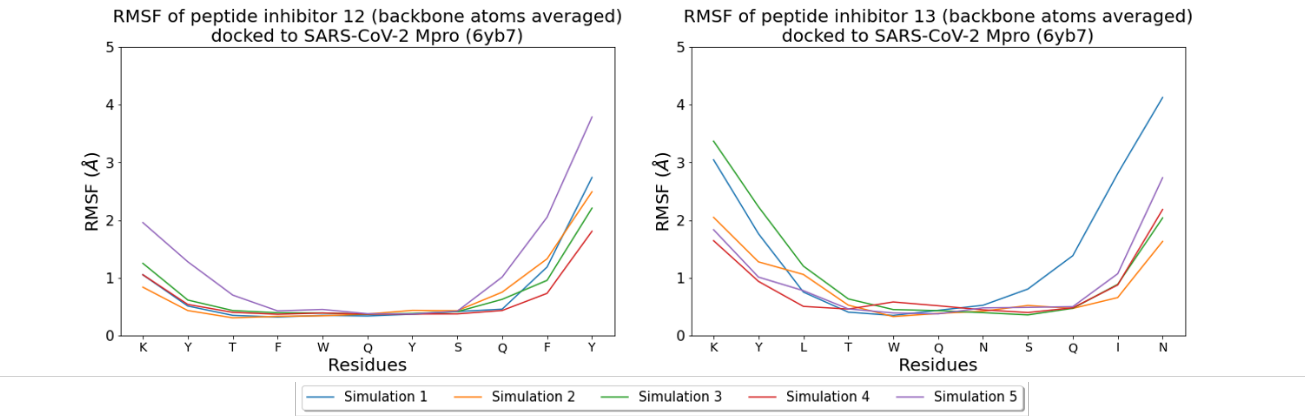

**Figure S3.9**: RMSF of p12 and p13 backbone atoms across 5 × 500 ns MD simulations of iMD-VR docked structures. In simulations of p12, the P' side of the peptide (residues -YSQFY) is more flexible than the P side (residues KYTFWQ-). In simulations of p13, the P' and P sides of the peptide seem equally flexible, with residues near the catalytic dyad (residues -WQN-) having a low RMSF.

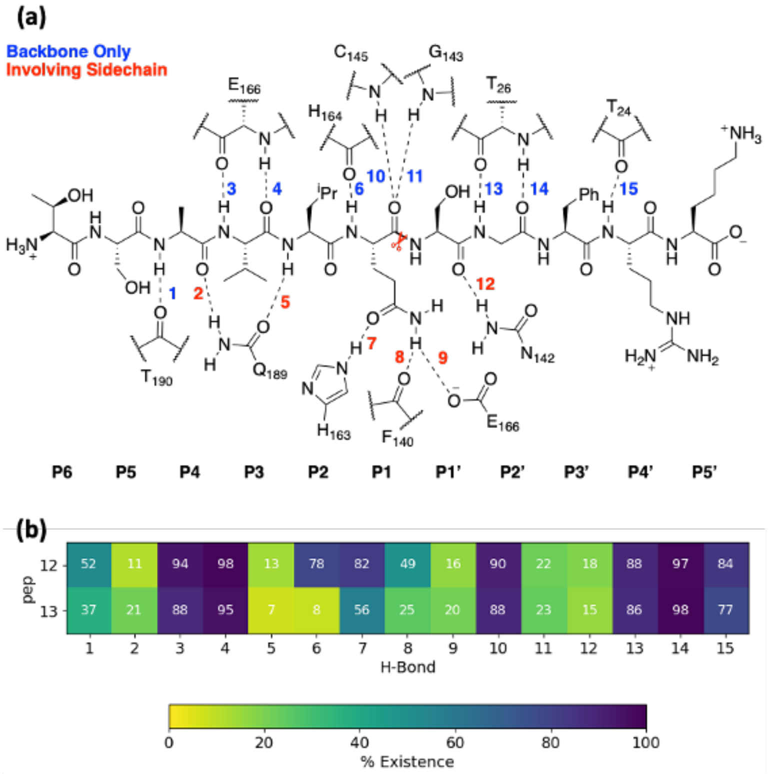

**Figure S3.10**: (a) Hydrogen bonds (HBs) analysed in the MD simulations of M$^{pro}$ complexed with p12 and p13, with s01 illustrated as an example. (b) An annotated heatmap displaying the percentage to which each HB is observed in the simulations of M$^{pro}$ in complex with p12 and p13 in explicit solvent. Frames were extracted every ns from 600 ns of cumulative MD conducted per system.

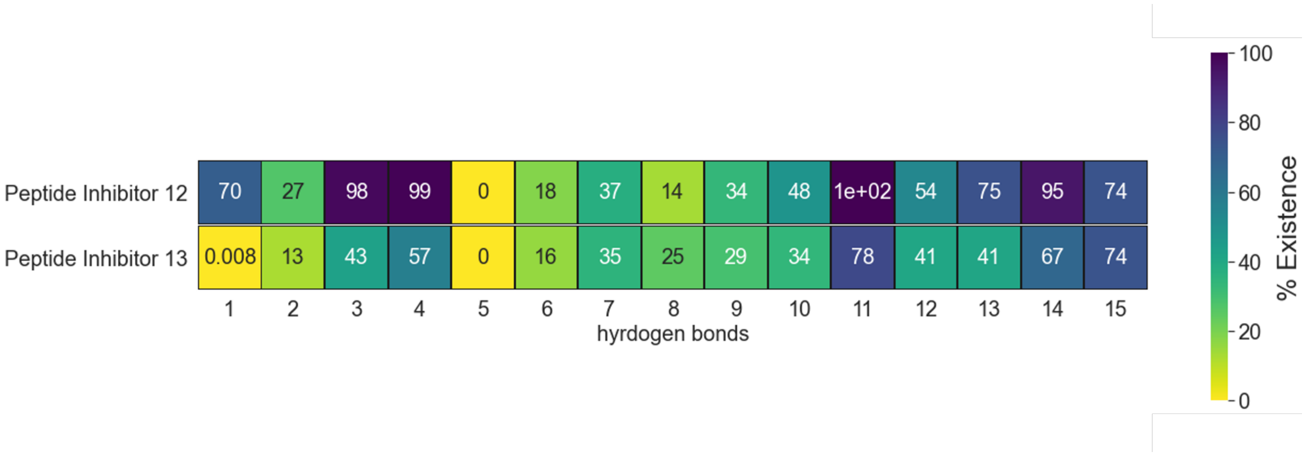

**Figure S3.11**: HBs present throughout the 5 × 500 ns MD simulations of each of the iMD-VR-docked designed peptide-M$^{pro}$ structures. Like substrates s01, s02 and s05 (**Figure S3.12**), the most well-maintained interactions in simulations of p12 are HBs 3, 4, 11, 13, and 14. On the other hand, p13 does not maintain these HBs as much. HB 5 is not observed in any of the simulations.

`

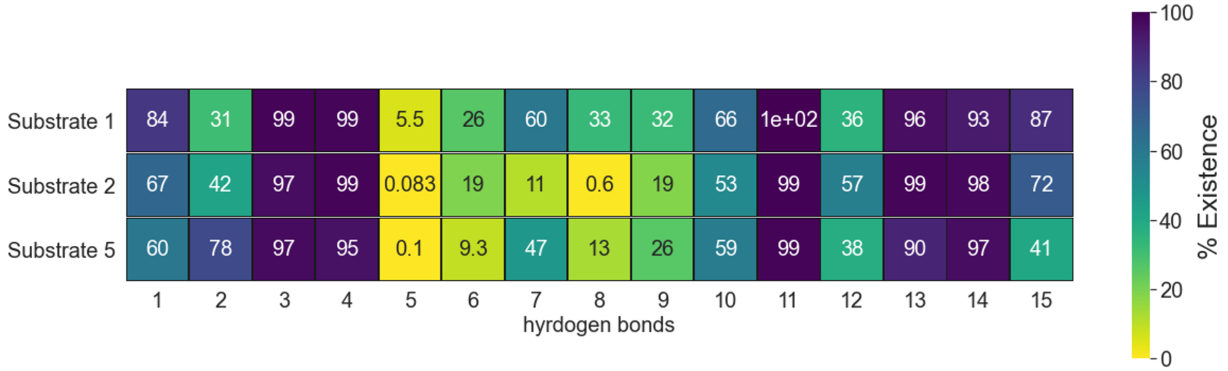

**Figure S3.12**: HBs present throughout the 3 × 200 ns MD simulations of each of the iMD-VR-docked substrate-M^pro structures (s01, s02, s05) from iMD-VR, in comparison with the designed peptide-M^pro complexes (p12 and p13, **Figure S3.11**). The most well-maintained HBs across all three substrates considered here are HBs 3, 4, 11, 13, and 14.

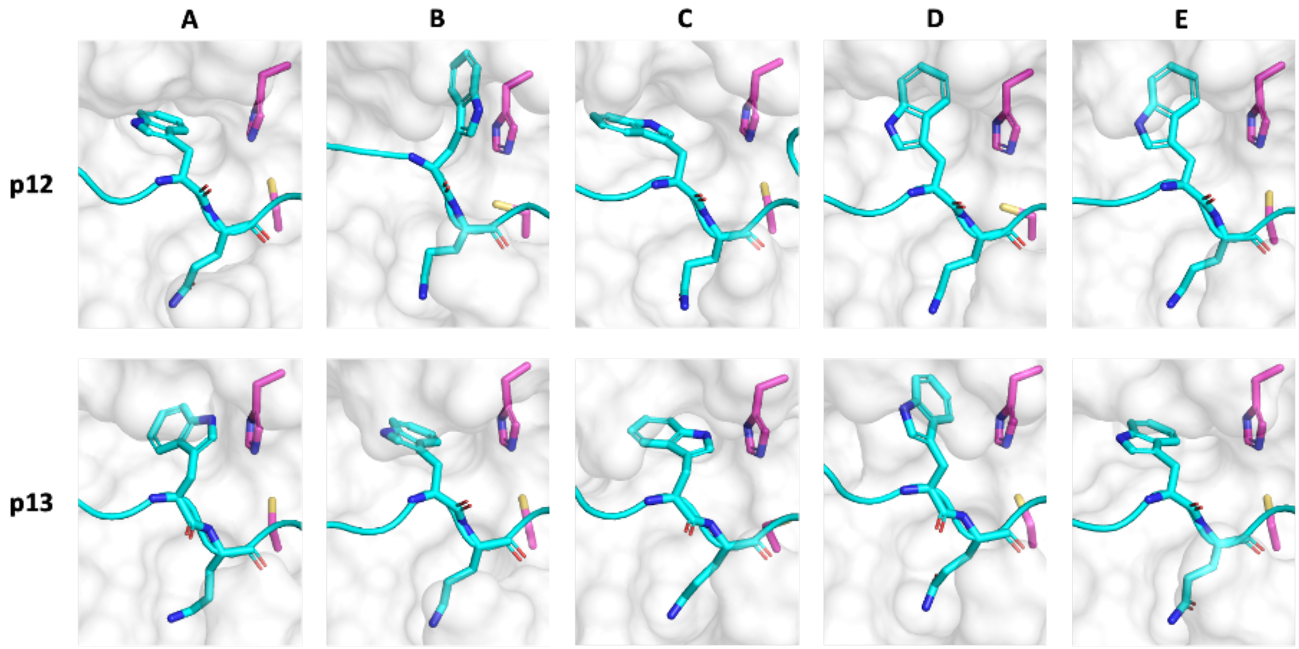

**Figure S3.13**: Conformations adopted by the P2 Trp side chain during MD simulations of M$^{pro}$ in complex with p12 (top row) and p13 (bottom row) in explicit solvent, as visualised with representative snapshots obtained from RMSD clustering. Frames extracted every ns were aligned using the backbone of surrounding M$^{pro}$ residues (His-41, Thr-45, Ser-46, Met-49, Pro-52, Tyr-54, Asn-142, Cys-145, His-164, Met-165, Glu-166, Phe-181, Val-186, Asp-187, Arg-188, Gln-189), and then clustered (gromos method, GROMACS v. 2019.2)[30] with a 2 Å cut-off on the RMSD of the P2 Trp side chain heavy atoms, resulting in 5 clusters for each of p12 and p13 (named A-E ranked by population; see **Figure S3.14**). M$^{pro}$ is presented as a white surface, with His-41 and Cys-145 shown in magenta. Only the P2 and P1 residues on the peptide are shown with the rest of the peptide backbone displayed as a ribbon (cyan). Hydrogens are omitted for clarity. In certain conformations (e.g. p12 conformation B; p13 conformations A and C) the P2 Trp side chain is in close proximity to the M$^{pro}$ residue His-41 which forms part of the catalytic dyad.

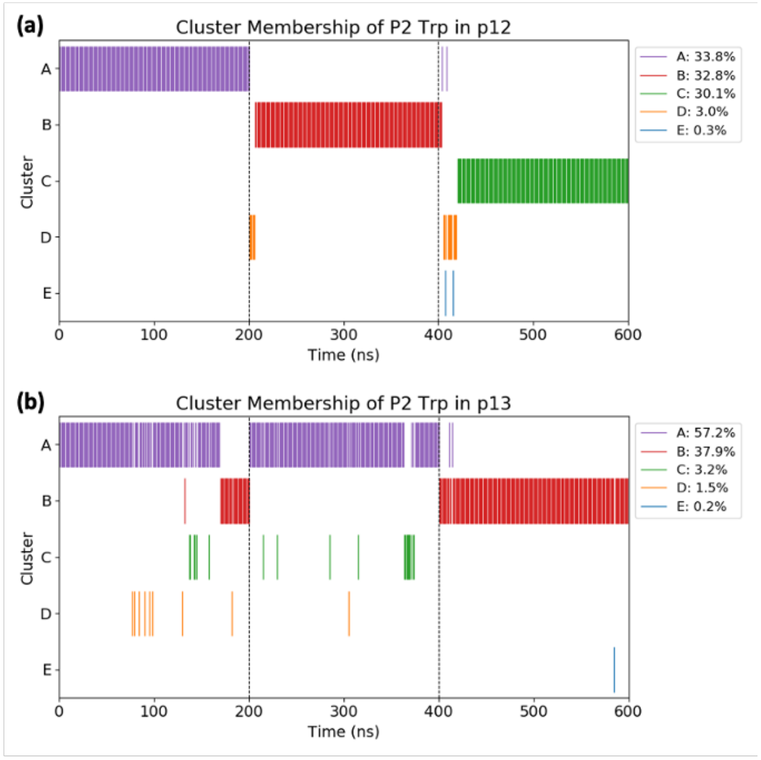

**Figure S3.14**: Evolution of the cluster membership (see **Figure S3.13** for representative structures) during the concatenated 3 × 200 ns MD simulations of M$^{pro}$ in complex with (a) p12 and (b) p13 in explicit solvent. The percentage population of each cluster for each system is displayed in the legend.

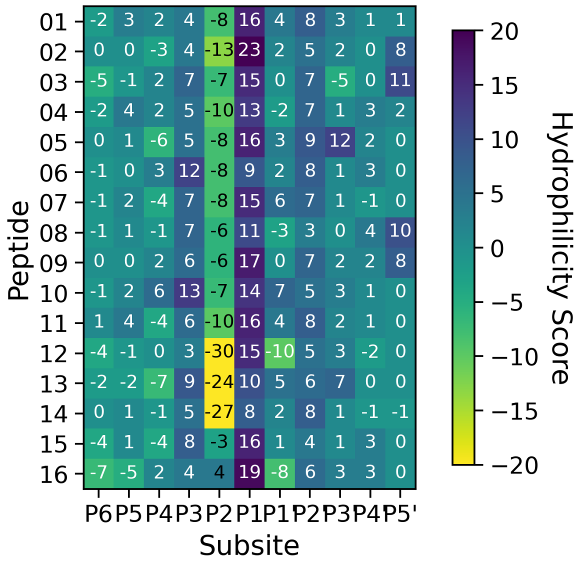

**Figure S3.15**: Hydrophilicity map of the eleven 11-mer substrate peptides (s01-s11) and the designed peptides (p12-p16). Hydrophilicity scores were calculated as a sum of all hydrophilic contacts subtracted by the sum of all hydrophobic contacts of each residue in the peptide. More positive scores correlate to hydrophilic sites and negative scores to hydrophobic pockets.

### Comparative peptide docking

**Table S3.3**: Details of the AutoDock CrankPep (ADCP) search space in Cartesian coordinates as identified by AutoSite (v. 1.1),[49] using the protein pdbqt file prepared with ADFRsuite from the corresponding PDB entry.

| Receptor | 2q6g (chain A) | 7bqy (dimer) | 7joy (dimer) |
|---|---|---|---|
| Centre (Å) | 95.708 0.477 3.342 | -10.000 0.189 -22.500 | 44.694 22.055 23.850 |
| Box length (Å) | 22.500 23.250 22.500 | 20.250 25.500 23.250 | 20.250 21.000 29.250 |
| Size (0.375 Å spacing) | 60 62 60 | 54 68 62 | 54 56 78 |
| Number of fill points | 76 | 82 | 97 |

**Table S3.4**: Assessment of the 10 highest ranked binding poses from ADCP redocking of the s01 sequence in the crystal structure of H41A SARS-CoV $M^{pro}$ (PDB 2q6g, chain A).[1] These poses are compared with the original position of the peptide by measuring the positional deviations of the 11 peptide Cα atoms. A pose is considered to pass the filter if the deviation is lower than 2 Å for at least three Cα atoms.

| Rank | Score (kcal/mol) | Cluster Size | Cα Deviation (Å) at Position | | | | | | | | | | | Filter | Cα RMSD (Å) |
|---|---|---|---|---|---|---|---|---|---|---|---|---|---|---|---|
| | | | 1 | 2 | 3 | 4 | 5 | 6 | 7 | 8 | 9 | 10 | 11 | | |
| 1 | -23.0 | 34 | 1.97 | 0.86 | 0.64 | 0.54 | 0.51 | 0.63 | 0.78 | 3.30 | 5.51 | 11.88 | 16.27 | Y | 6.42 |
| 2 | -21.5 | 9 | 2.67 | 1.27 | 0.96 | 1.08 | 0.81 | 0.63 | 1.53 | 5.36 | 5.96 | 13.13 | 15.44 | Y | 6.67 |
| 3 | -21.5 | 56 | 10.09 | 7.43 | 9.61 | 9.62 | 13.41 | 12.67 | 12.99 | 12.34 | 12.33 | 14.52 | 13.99 | N | 11.92 |
| 4 | -21.3 | 10 | 5.38 | 1.43 | 0.75 | 0.70 | 0.70 | 0.60 | 1.46 | 5.07 | 4.42 | 4.37 | 6.31 | Y | 3.56 |
| 5 | -21.0 | 14 | 2.06 | 0.56 | 0.29 | 0.50 | 0.49 | 0.68 | 1.48 | 4.07 | 5.95 | 12.99 | 16.06 | Y | 6.65 |
| 6 | -20.6 | 29 | 12.95 | 12.54 | 12.24 | 10.68 | 9.29 | 6.19 | 9.27 | 12.97 | 15.77 | 21.98 | 24.28 | N | 14.42 |
| 7 | -20.5 | 15 | 18.31 | 12.78 | 6.37 | 1.23 | 6.30 | 11.08 | 15.61 | 18.46 | 17.80 | 19.15 | 16.65 | N | 14.30 |
| 8 | -20.5 | 21 | 5.66 | 1.22 | 0.81 | 0.84 | 0.48 | 0.19 | 1.20 | 6.05 | 4.47 | 5.40 | 3.49 | Y | 3.50 |
| 9 | -20.5 | 22 | 16.08 | 12.80 | 10.71 | 6.98 | 10.03 | 9.77 | 10.03 | 9.31 | 9.76 | 9.68 | 9.88 | N | 10.69 |
| 10 | -20.3 | 25 | 14.76 | 11.41 | 9.30 | 6.81 | 7.17 | 5.02 | 5.52 | 5.98 | 6.16 | 11.39 | 16.88 | N | 9.89 |

`

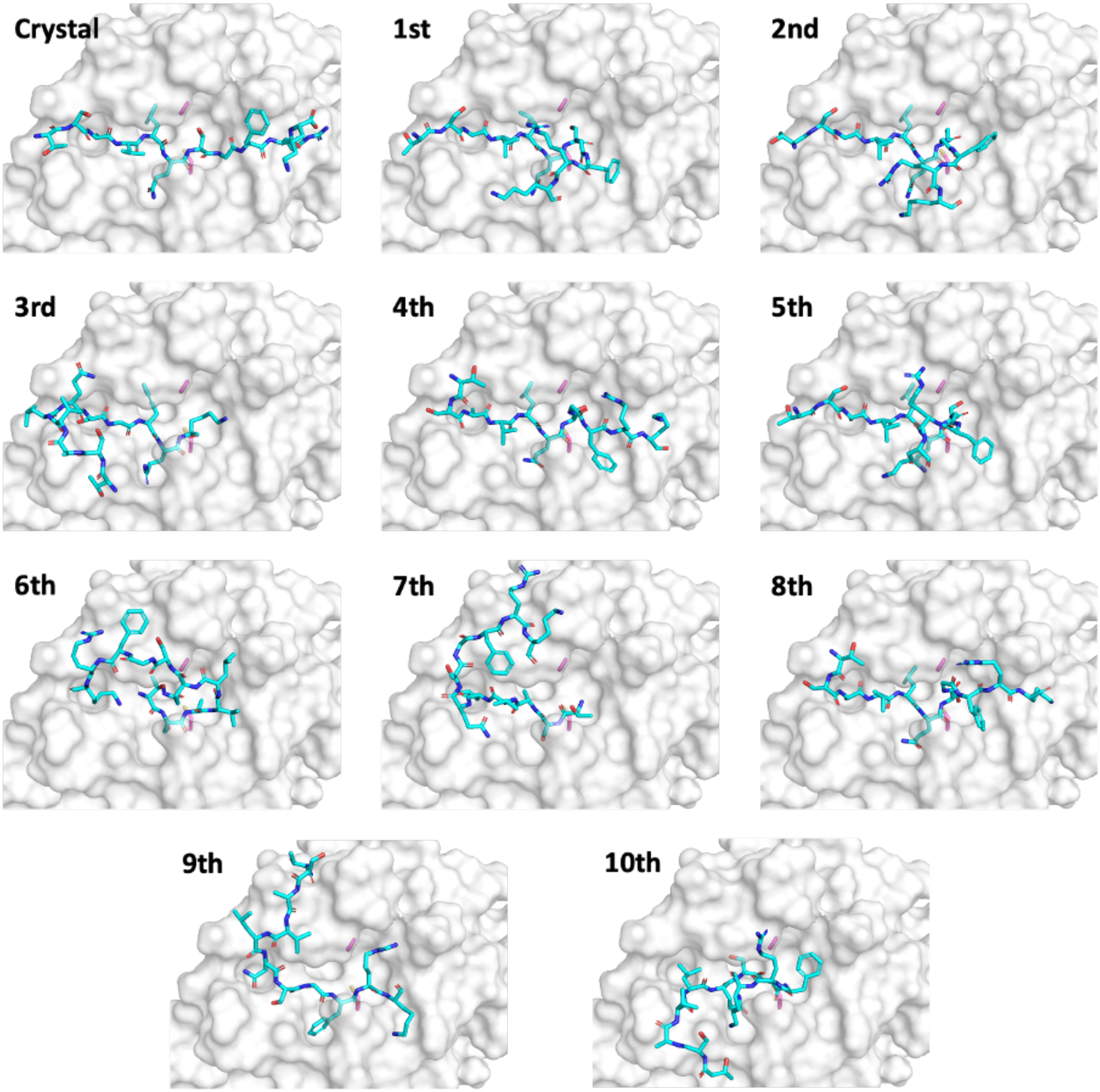

**Figure S3.16**: The 10 highest ranked binding poses from ADCP redocking of the s01 sequence (cyan) in the crystal structure of H41A SARS-CoV M$^{pro}$ (PDB 2q6g), compared to the original position in the crystal structure (top left).[1] Ala-41 (in place of His that is present in the catalytically competent protein) and Cys-145 are in magenta.

**Table S3.5**: Assessment of selected binding poses from ADCP docking of each substrate (s01-s11) and designed (p12, p13, p15, p16) sequence in the crystal structure of SARS-CoV-2 M$^{pro}$, originally in complex with the N3 inhibitor (PDB 7bqy).[47] For each peptide sequence, the 10 highest ranked solutions were evaluated by comparison to the M$^{pro}$-peptide complex resulting from comparative modelling followed by MM minimisation. For every sequence, the highest ranked solution that passed the filter of < 2 Å deviation in at least three C$\alpha$ atoms, or if none of the solutions passed the filter, the solution with the lowest C$\alpha$ RMSD, was presented below.

| Seq | Pose Rank | Score (kcal/mol) | Cluster Size | Cα Deviation (Å) at Position | | | | | | | | | | | Filter | Cα RMSD (Å) |
|---|---|---|---|---|---|---|---|---|---|---|---|---|---|---|---|---|
| | | | | 1 | 2 | 3 | 4 | 5 | 6 | 7 | 8 | 9 | 10 | 11 | | |
| 01 | 1 | -22 | 55 | 2.19 | 1.49 | 1.16 | 1.60 | 1.67 | 2.36 | 7.95 | 9.34 | 13.47 | 19.87 | 24.00 | Y | 10.96 |
| 02 | 6 | -19.9 | 19 | 7.32 | 3.87 | 1.60 | 2.25 | 3.23 | 2.85 | 3.99 | 3.59 | 4.21 | 4.56 | 7.58 | N | 4.47 |
| 03 | 7 | -20.5 | 27 | 5.83 | 3.10 | 1.31 | 1.52 | 1.37 | 1.38 | 7.22 | 10.34 | 13.72 | 20.06 | 23.77 | Y | 11.14 |
| 04 | 1 | -21 | 65 | 5.62 | 1.42 | 0.91 | 0.91 | 0.80 | 0.85 | 0.87 | 2.97 | 4.46 | 6.28 | 5.53 | Y | 3.52 |
| 05 | 7 | -21.1 | 10 | 5.24 | 3.29 | 1.03 | 1.76 | 0.99 | 0.40 | 0.58 | 0.87 | 3.11 | 5.93 | 8.52 | Y | 3.84 |
| 06 | 1 | -21 | 15 | 1.96 | 1.26 | 0.87 | 1.46 | 0.81 | 0.77 | 0.84 | 0.94 | 4.86 | 10.76 | 9.86 | Y | 4.75 |
| 07 | 2 | -19.7 | 25 | 0.75 | 0.94 | 1.25 | 2.32 | 1.17 | 0.84 | 0.24 | 0.64 | 5.15 | 10.78 | 10.89 | Y | 4.98 |
| 08 | 1 | -20.3 | 33 | 5.64 | 2.98 | 1.01 | 1.07 | 0.39 | 0.42 | 0.45 | 1.31 | 3.20 | 10.20 | 13.65 | Y | 5.61 |
| 09 | 2 | -20.3 | 27 | 7.94 | 1.84 | 1.42 | 1.71 | 2.82 | 2.60 | 7.45 | 9.81 | 12.79 | 18.27 | 20.35 | Y | 10.22 |
| 10 | 4 | -21 | 26 | 2.82 | 1.66 | 0.92 | 1.03 | 1.60 | 3.84 | 4.74 | 5.49 | 2.01 | 2.09 | 2.06 | Y | 2.94 |
| 11 | 1 | -22.5 | 55 | 2.45 | 2.07 | 1.29 | 1.84 | 1.94 | 2.11 | 7.87 | 11.78 | 14.77 | 21.15 | 25.07 | Y | 11.75 |
| 12 | 1 | -26 | 39 | 12.02 | 9.00 | 4.42 | 1.87 | 8.42 | 10.96 | 10.80 | 8.39 | 6.22 | 7.43 | 13.85 | N | 9.11 |
| 13 | 9 | -23.3 | 6 | 10.24 | 7.29 | 7.43 | 6.88 | 6.79 | 7.65 | 5.91 | 5.70 | 10.52 | 15.64 | 19.69 | N | 10.34 |
| 15 | 1 | -24 | 46 | 10.46 | 5.99 | 4.72 | 3.51 | 4.13 | 4.74 | 4.81 | 3.73 | 3.89 | 4.16 | 6.90 | N | 5.53 |
| 16 | 5 | -24 | 32 | 8.09 | 3.32 | 4.37 | 1.95 | 0.91 | 1.17 | 2.40 | 3.84 | 5.64 | 13.00 | 15.83 | Y | 7.22 |

`

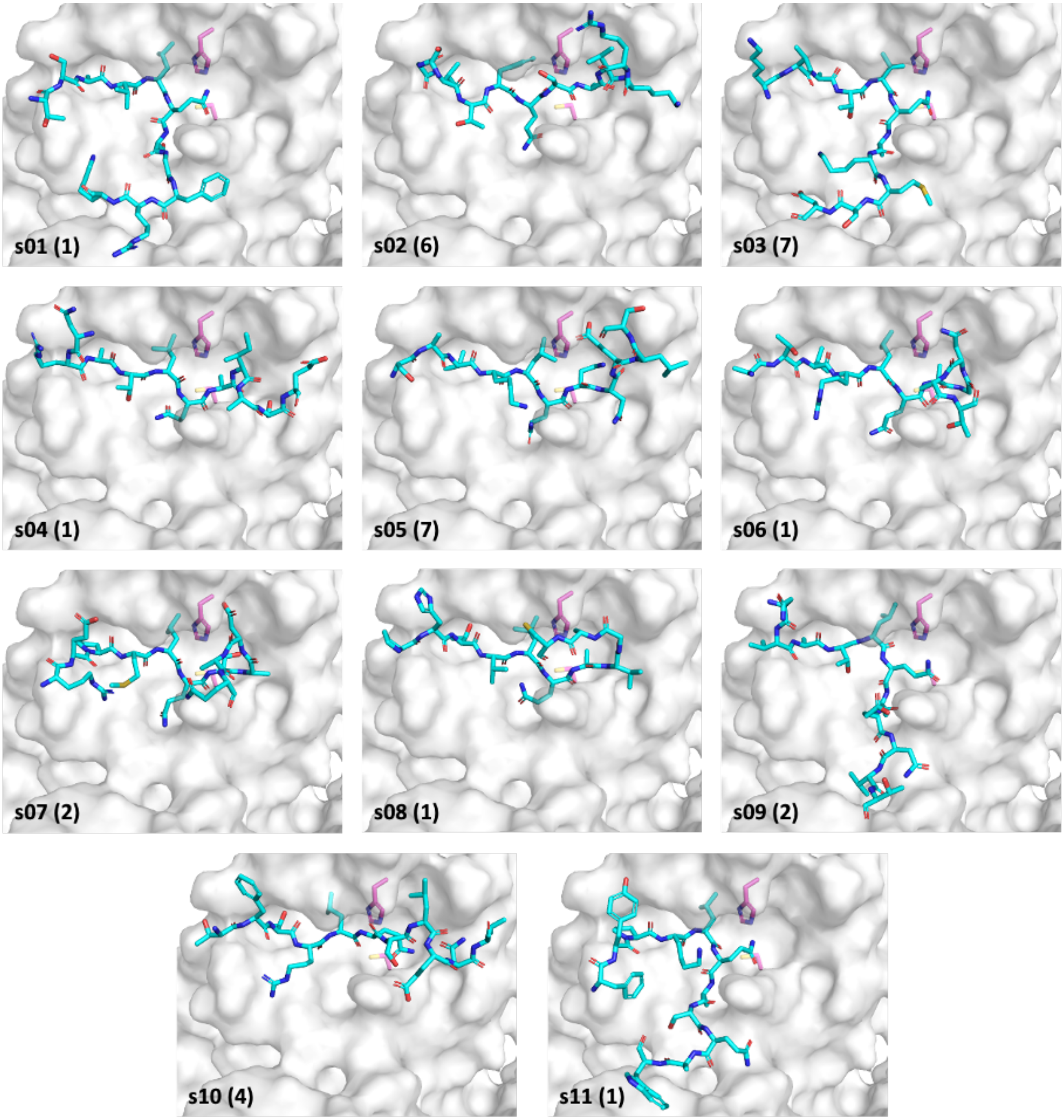

**Figure S3.17**: Selected binding poses (see **Table S3.5**; number in parentheses = pose rank) from ADCP docking of each native substrate (s01-s11) sequence in the crystal structure of SARS-CoV-2 M$^{pro}$, originally in complex with the N3 inhibitor (PDB 7bqy). The catalytic dyad His-41 and Cys-145 are in magenta. Docked structures with the P4 and P2 residues positioned correctly in their respective S4 and S2 pockets were consistently obtained. However, greater variation was observed in the positioning of residues from P1 onwards, likely due to the poorer definition of the S′ subsites.

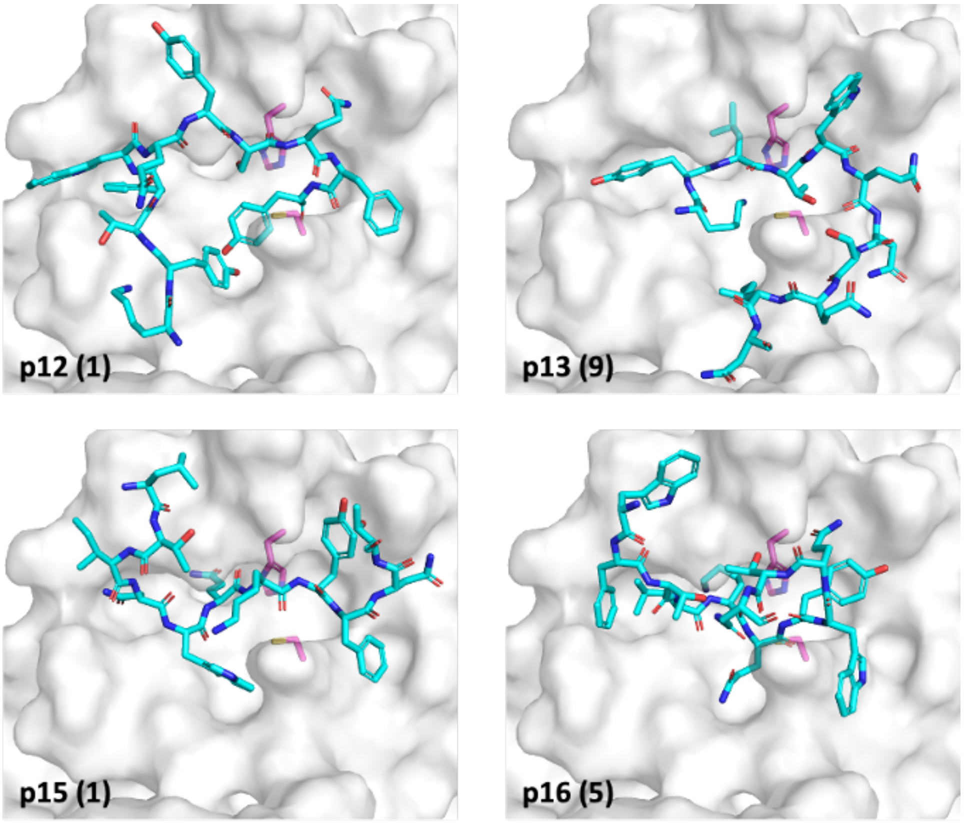

**Figure S3.18**: Selected binding poses (see **Table S3.5**; number in parentheses = pose rank) from ADCP docking of each designed (p12, p13, p15, p16) sequence in the crystal structure of SARS-CoV-2 M$^{pro}$, originally in complex with the N3 inhibitor (PDB 7bqy). The catalytic dyad His-41 and Cys-145 are in magenta.

**Table S3.6**: Assessment of selected binding poses from ADCP docking of each designed (p12, p13, p15, p16) sequence in the crystal structure of C145A SARS-CoV-2 M$^{pro}$, originally in complex with the s02 cleaved product (PDB 7joy).[48] For each peptide sequence, the 10 highest ranked solutions were evaluated by comparison to the M$^{pro}$-peptide complex resulting from comparative modelling followed by MM minimisation. The highest ranked solution that passed the filter of < 2 Å deviation in at least three Cα atoms, or if none of the solutions passed the filter, the pose with the lowest Cα RMSD, was presented below.

| Seq | Pose Rank | Score (kcal/mol) | Cluster Size | Cα Deviation (Å) at Position | | | | | | | | | | | Filter | Cα RMSD (Å) |
|---|---|---|---|---|---|---|---|---|---|---|---|---|---|---|---|---|
| | | | | 1 | 2 | 3 | 4 | 5 | 6 | 7 | 8 | 9 | 10 | 11 | | |
| 12 | 1 | -29.6 | 63 | 5.75 | 1.95 | 1.83 | 1.20 | 0.99 | 1.06 | 1.46 | 1.80 | 3.81 | 4.99 | 6.94 | Y | 3.53 |
| 13 | 4 | -23.3 | 35 | 8.06 | 4.05 | 4.76 | 4.22 | 3.83 | 1.74 | 3.62 | 3.22 | 4.36 | 5.04 | 6.80 | N | 4.80 |
| 15 | 2 | -25.7 | 29 | 4.23 | 5.33 | 4.02 | 4.61 | 4.58 | 3.87 | 3.76 | 4.37 | 3.89 | 7.05 | 5.78 | N | 4.78 |
| 16 | 1 | -27.7 | 69 | 4.57 | 1.60 | 1.85 | 0.79 | 0.65 | 0.85 | 1.06 | 1.62 | 3.94 | 5.64 | 6.84 | Y | 3.39 |

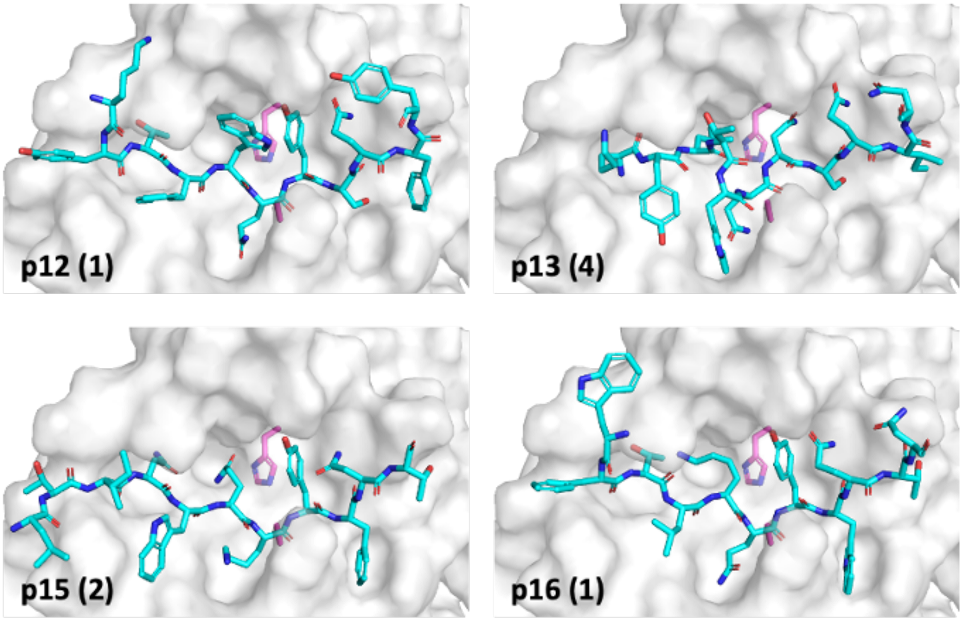

**Figure S3.19**: Selected binding poses (see **Table S3.6**; number in parentheses = pose rank) from ADCP docking of each designed (p12, p13, p15, p16) sequence in the crystal structure of C145A SARS-CoV-2 M$^{pro}$, originally in complex with the s02 product (PDB 7joy). The catalytic dyad His-41 and Ala-145 (in place of Cys-145 that is present in the catalytically competent protein) are in magenta.

### S4. Supplementary Results – Analysis of Results from Fragment Crystallography

#### Interaction analysis of the XChem fragments

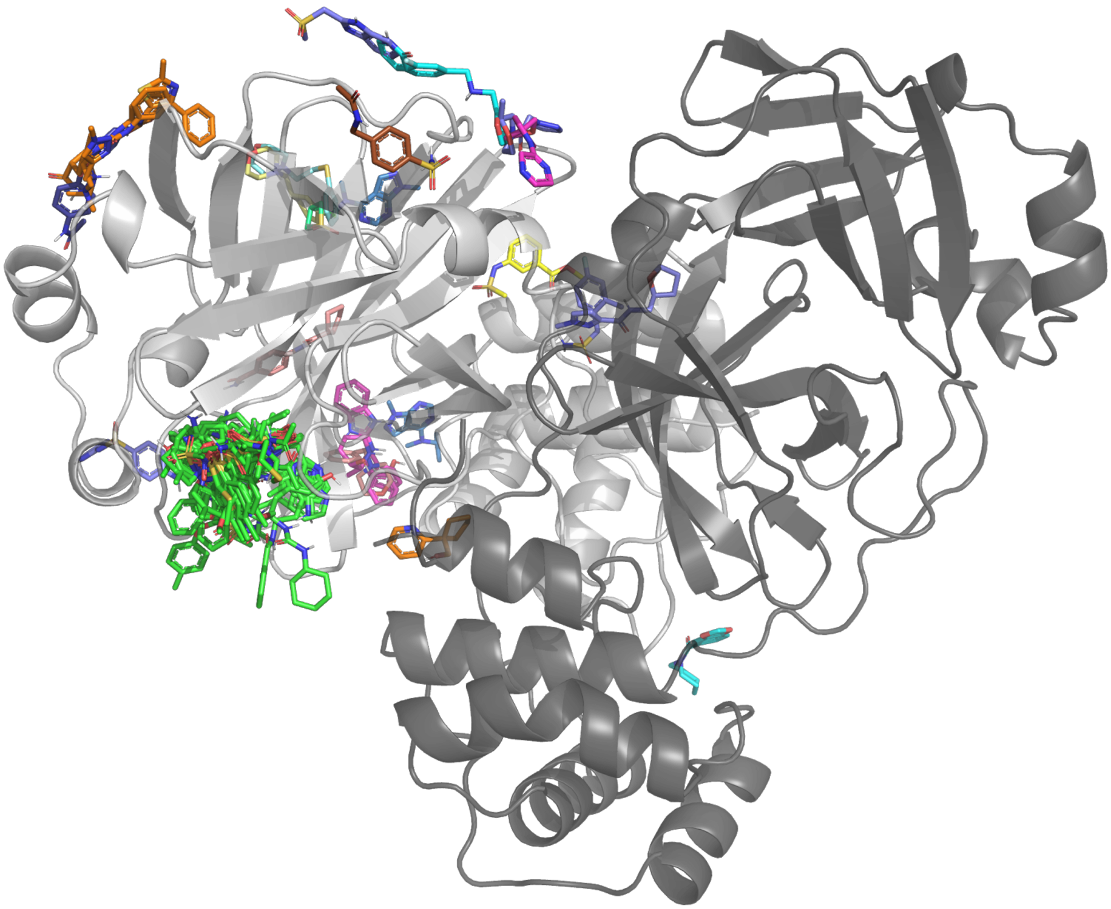

**Figure S4.1**: Views from crystal structures of all XChem fragments and their binding site on the M^pro^ dimer.[37] All binding sites are on chain A (white). As a representative structure the fragment x0830 co-crystal structure was used. There are 66 fragments that bind into the active site (green fragments) and 25 allosteric fragments (fragment 1101 binds in two different allosteric sites).

`

**Table S4.1**: Relationship between Tanimoto similarity threshold and cluster sizes for the XChem fragment crystal structures. Only active-site binders were considered.

| Tanimoto threshold | Number of clusters | Number of single molecule clusters | Average cluster size |
|---|---|---|---|
| 0.1 | 2 | 0 | 33.0 |
| 0.2 | 2 | 0 | 33.0 |
| 0.3 | 5 | 1 | 13.2 |
| 0.4 | 9 | 0 | 7.3 |
| 0.5 | 11 | 2 | 6.0 |
| 0.6 | 16 | 4 | 4.1 |
| 0.7 | 29 | 20 | 2.3 |
| 0.8 | 37 | 27 | 1.8 |
| 0.9 | 53 | 46 | 1.2 |

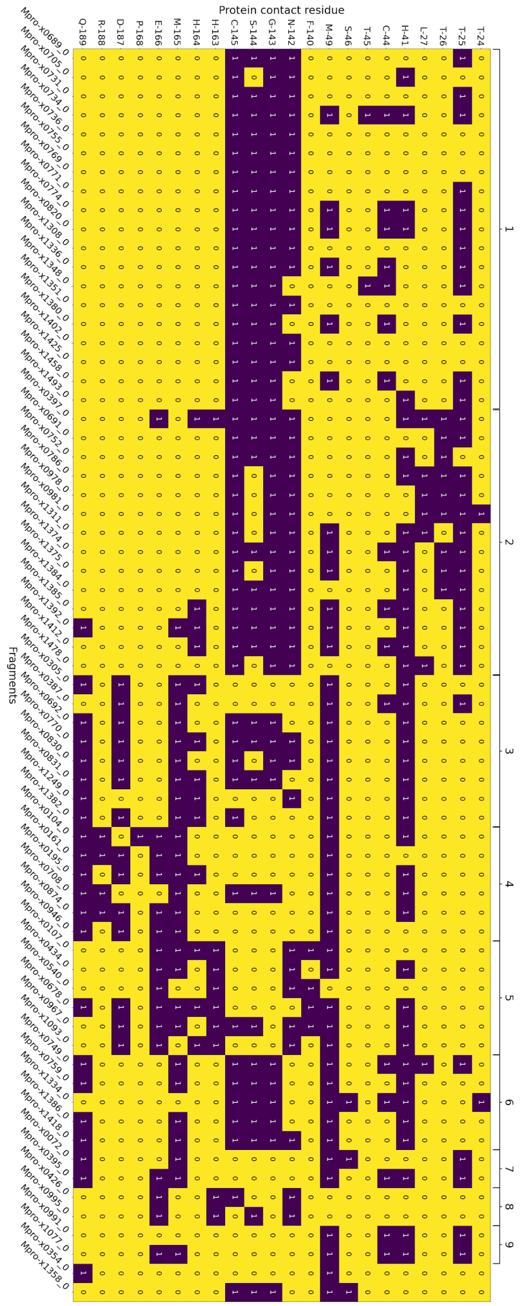

**Figure S4.2**: Contact matrix for the 66 active site XChem fragments sorted by their assigned cluster (threshold 0.5) based on the protein contacts which are indicated by “1”. Clusters 1 through 9 are indicated by brackets. Clusters with more than 1 molecule are marked by brackets (except x0354 and x1358).

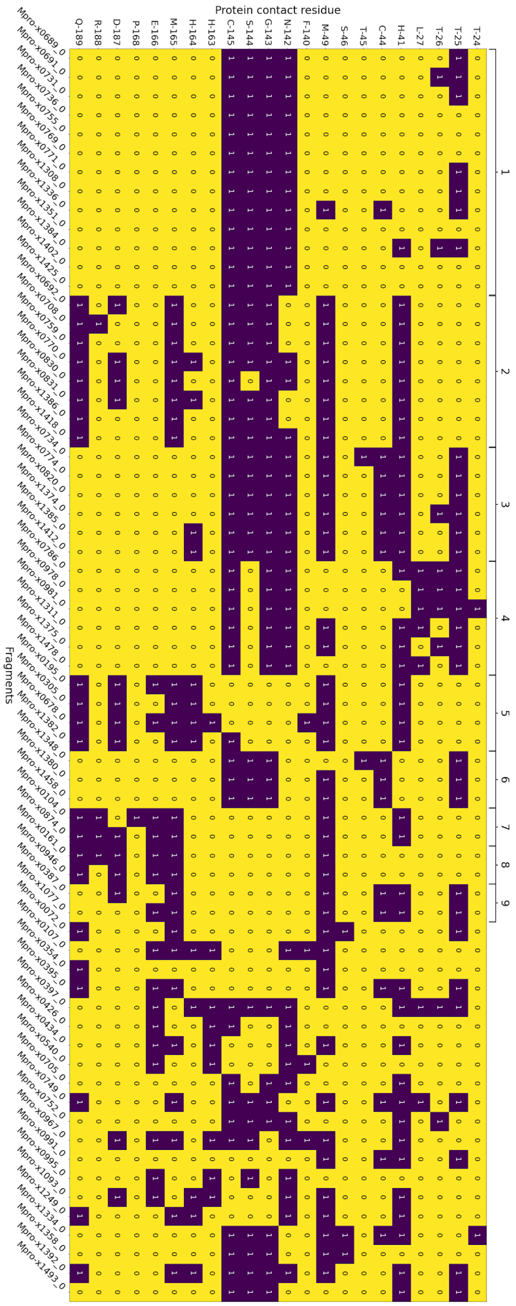

**Figure S4.3**: Contact matrix for the 66 active site XChem fragments sorted by their assigned cluster (threshold 0.7) based on the protein contacts which are indicated by "1". Clusters 1 through 9 are indicated by brackets. Only clusters with more than 1 molecule are marked by brackets.

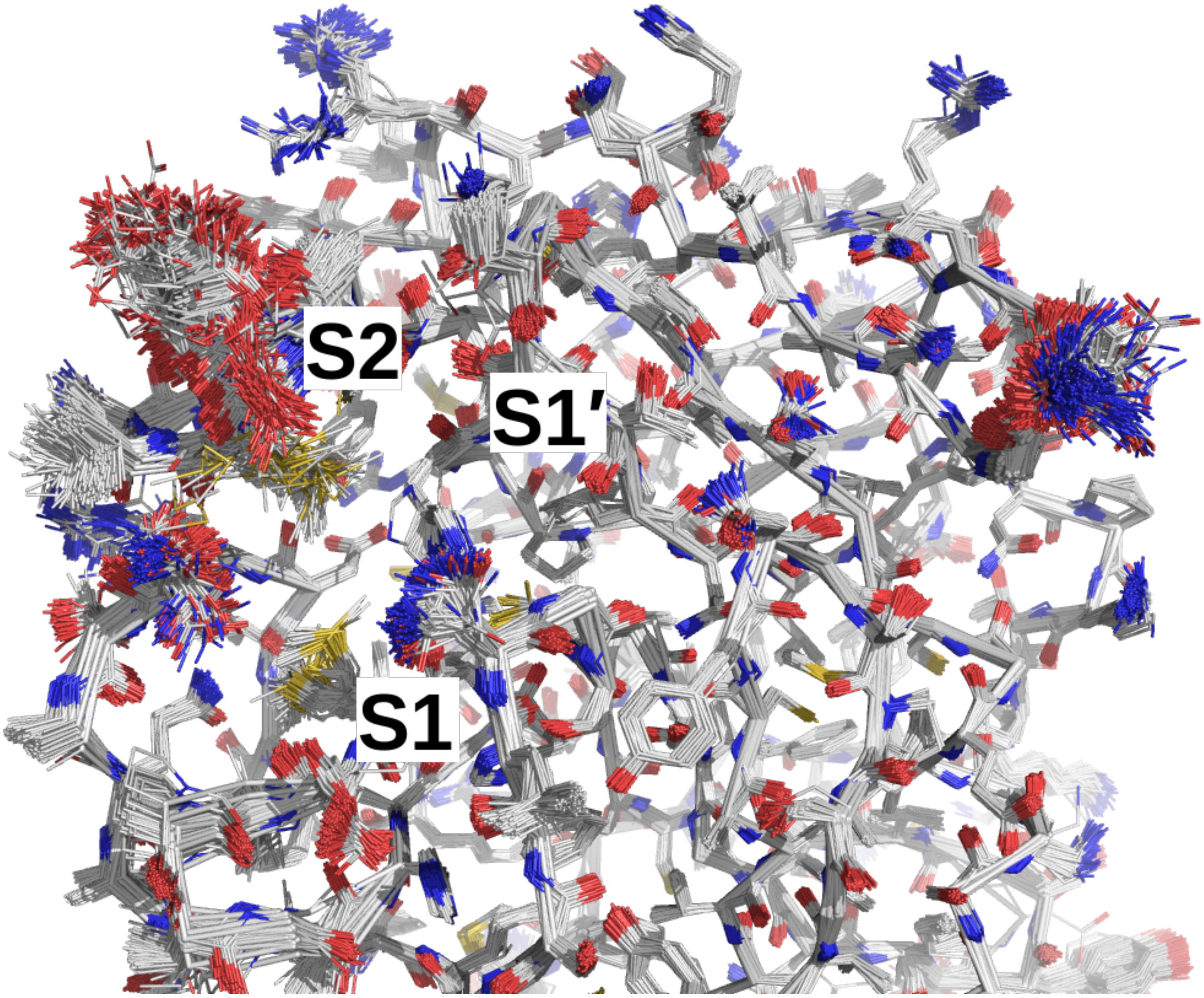

**Figure S4.4:** Overlay of views of the 333 M$^{pro}$ co-crystal structures published by Fragalysis.[52] Subsite S2 shows extremely large changes in conformation between structures while S1 is conserved.

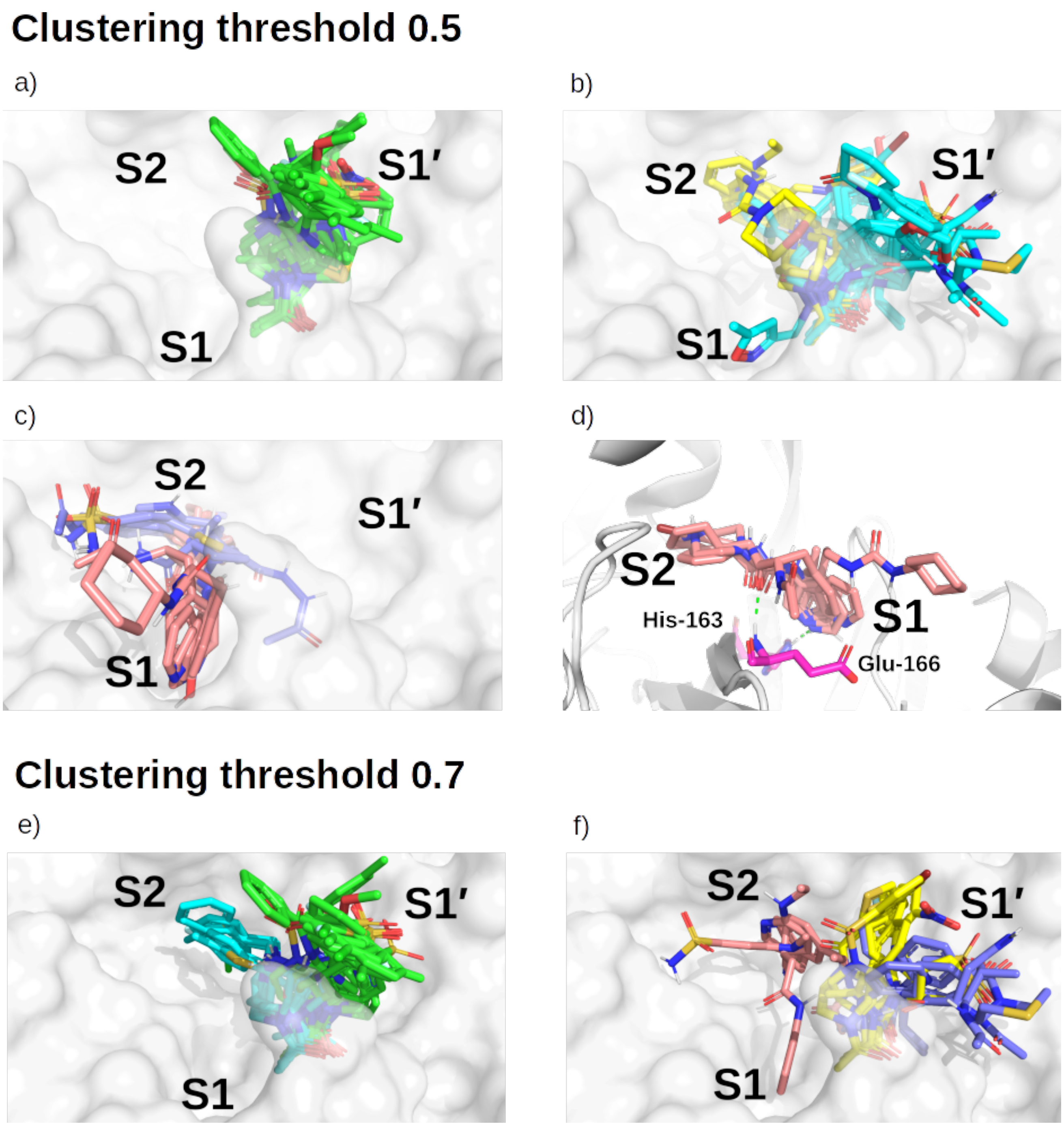

**Figure S4.5**: M$^{pro}$ crystal structure (x0830) in complex with the top 5 most populated clusters using a clustering threshold of 0.5: a) cluster 1 (green); b) clusters 2 (cyan) and 3 (yellow); c) clusters 4 (blue) and 5 (salmon). d) Close-up on the binding pose of cluster 5. Shown in green are the two key HBs between the fragment carbonyl oxygen and the backbone nitrogen of Glu-166 (HB 3 as identified in main text **Figure 3**), and between the His-163 Nε and the nitrogen heterocycle of the fragment (HB 6 as identified in main text **Figure 3**). Also shown are the top 5 most populated clusters using a threshold of 0.7: e) clusters 1 (green) and 2 (cyan); f) clusters 3 (yellow), 4 (blue) and 5 (salmon).

**Descriptor based on contact and long-range interactions**

To further understand how the XChem fragments interact with $M^{pro}$, we performed linear scaling DFT calculations, as done for the substrate/designed peptides. Both short-range and long-range interaction terms were analysed and the computed coupling strengths between fragments and enzyme residues were compared to those obtained for the natural substrates, with the aim to identify potentially interesting inhibitor candidates. We have made these results publicly available in the following links:
Contact Interaction:
https://maayanlab.cloud/clustergrammer/viz/603f6484d0867e01721c4820/contact_interactions_subs_XChem.tsv
Electrostatic Interaction:
https://maayanlab.cloud/clustergrammer/viz/603f624cd0867e01721c47e5/electrostatic_interactions_subs_XChem.tsv

A hierarchical visualisation of contact and long-range interactions enables identification of families of different inhibitors (**Figure S4.6** below; left and right panels respectively). Within this clustering approach, the fragments/inhibitors belonging to the same family as the native substrates (s01, s02, s05) are highlighted in red and the cluster 5 compounds identified with Arpeggio in bold red (**Section 4.1** of Main text), by considering the short-range DFT interaction as a descriptor. Remarkably, all cluster 5 compounds belong to the same family as the native substrates. Although this analysis is at a preliminary stage, we believe that this is potentially a powerful direction of investigation as it enables agnostic comparison between compounds of different size and nature based on first-principle considerations.

Using long-range electrostatic interactions as a descriptor, cluster 5 compounds split into two main subfamilies. One of the compounds (x0540) belongs to the same subgroup as the substrate peptides, while the remaining compounds are grouped into another subfamily with compound x0874. Compounds that show similar long-range interaction patterns but different contact interaction patterns as substrates s01/s02 are highlighted in purple.

This analysis can be interpreted in a twofold way: on one hand, it enables a criterion to single out some XChem compounds which provide interaction patterns similar to those exhibited by native substrates. We can thus generalize the cluster 5 group of compounds into a larger family, which also includes the compounds highlighted in red. On the other hand, when applying such a criterion to long-range interactions, other compounds (in purple) which share similarities with the native substrates can be identified. In the future, it will be interesting to further investigate such interaction pattern categorisation with the aim of identifying potent enzyme binders.

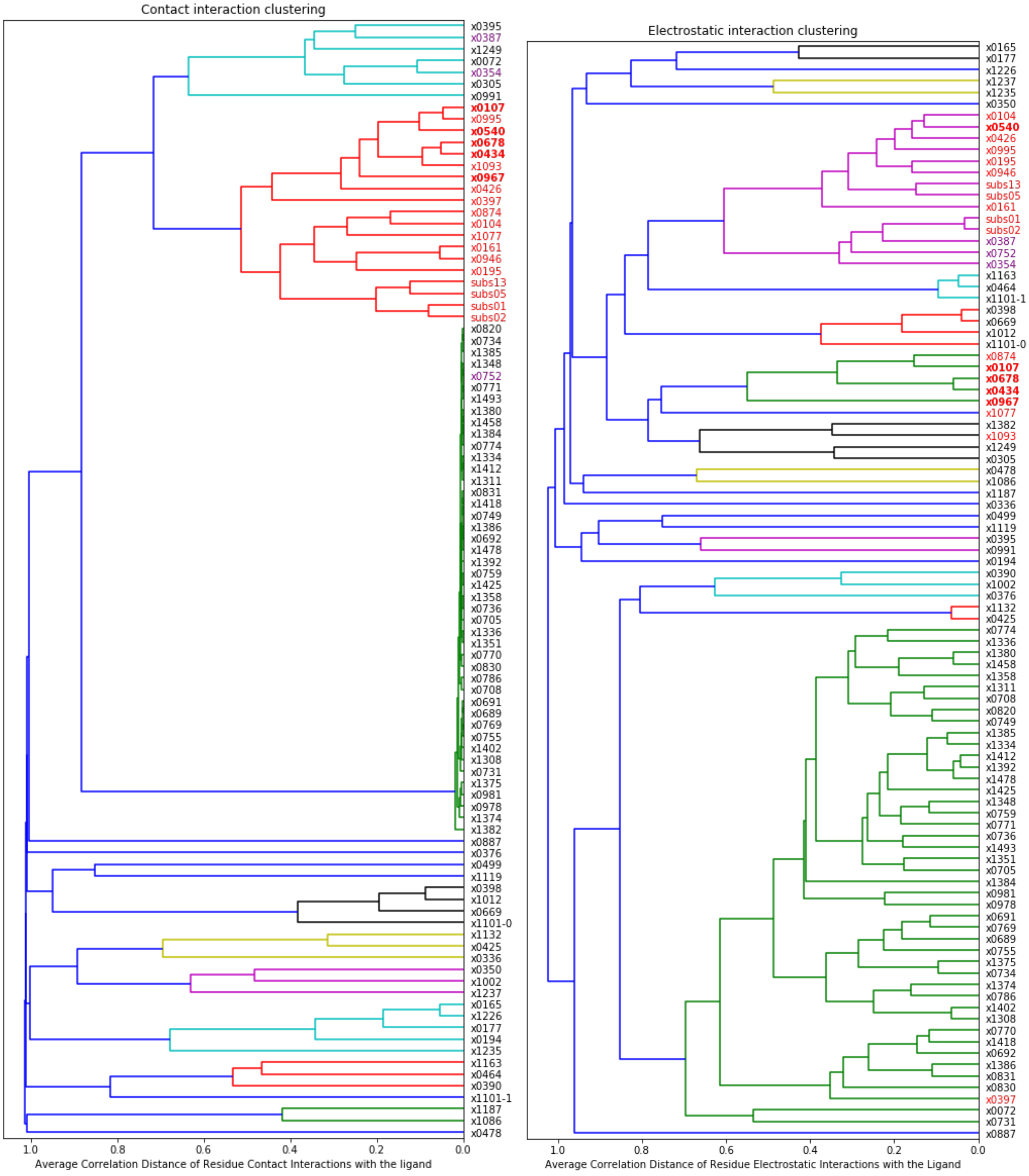

**Figure S4.6**: Dendrograms showing the hierarchical clustering of XChem fragment inhibitors, substrate peptides (s01, s02, s05), and designed peptide p13, considering descriptors defined on (left) short-range contact and (right) long-range DFT electrostatic interactions. The average correlation distance between interaction patterns is indicated on the x-axis. A family of compounds (red) which contains the peptides (s01, s02, s05, p13) as well as cluster 5 binders (bold) is identified using short-range interactions (left). Fragments that belong to the same family as s01 and s02 based on long-range (right) but not short-range interactions are highlighted in purple.

#### Interaction analysis of COVID Moonshot compounds

The kernel density estimation (KDE) and histogram of the distribution of the COVID Moonshot dataset for the fluorescence and RapidFire dataset are shown in **Figure S4.7**.[35]

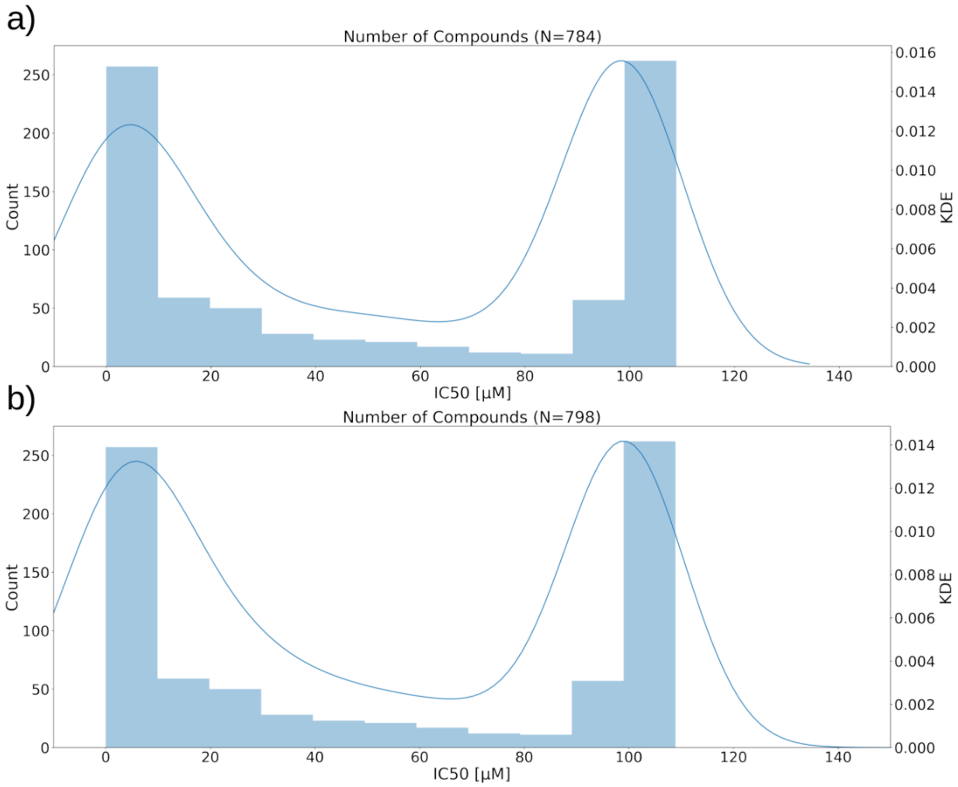

**Figure S4.7**: a) KDE and histogram for the RapidFire dataset. b) KDE and histogram for the fluorescence dataset.

`

**Covalent docking of COVID Moonshot compounds**

By selecting docked compounds that did not have significant modifications to the original fragment (less than 10 atoms difference to the MCS between fragment and docked compounds), the number of compounds regaining the binding pose of the fragment is further increased to 87 (54.7%) of the 159 compounds. Interestingly, the distribution of SuCOS over the data subsets resembles a bimodal distribution in all cases, with one peak around a SuCOS of 0.25 and a second peak between 0.7-0.8 depending on the subset (**Figure S4.8**). This suggests that the AD4 docking process performs well generally in identifying binding poses similar to the original fragment (especially in cases where few modifications were made), but fails completely in some cases, resulting in almost no overlap between the pose and the fragment and an extremely low SuCOS (<0.3). This observation is in line with the expected changes in binding mode of a molecule when subjected to large changes in structure and does not necessarily correspond to incorrect docked poses. As a result, when filtering for compounds where only minor changes were made to the inspiration fragment and the docking pose generated regained the same binding pose as the fragment, we increased the probability of gaining relevant poses. However, thorough validation of this hypothesis is yet to be done.

a)

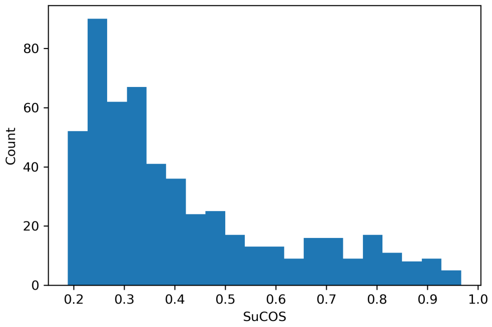

b)

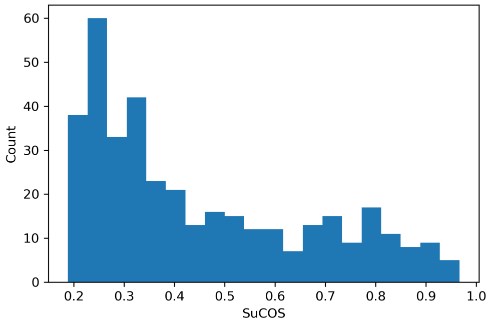

c)

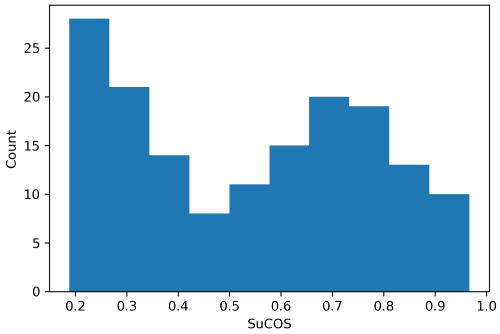

**Figure S4.8**: Distribution of SuCOS between the docked pose of the covalent Moonshot design and the original covalent fragment used as a basis for design. a) SuCOS scores of the 540 docked compounds; b) SuCOS scores of the 379 docked compounds with significant MCS overlap (i.e. more than 8 atoms MCS match) to the fragment; c) SuCOS scores of the 159 docked compounds with significant MCS overlap to the fragment and only small changes to its structure (<10 atoms difference between the compound and its MCS with the inspiration fragment).

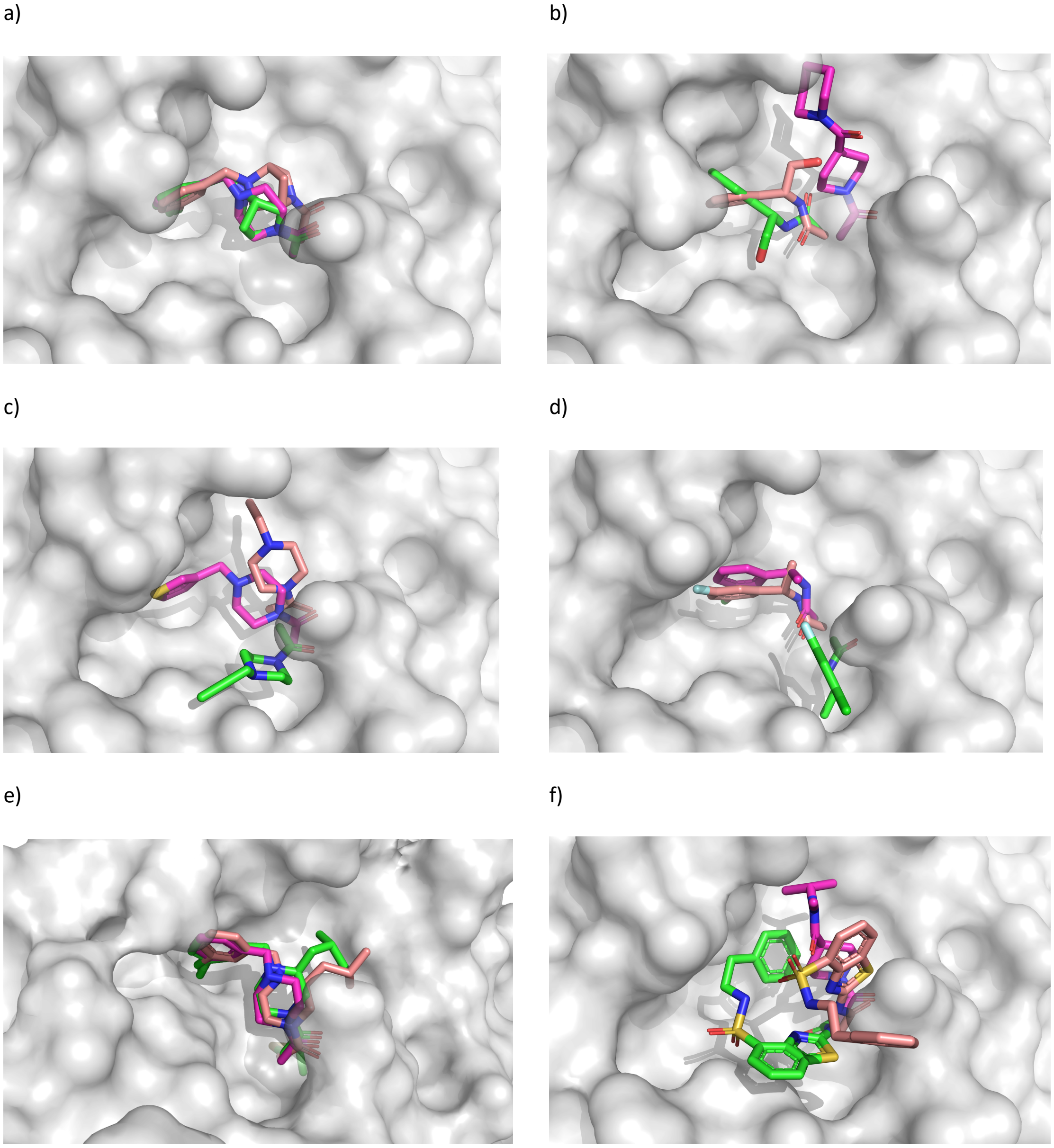

**Figure S4.9**: Overlay of the lowest energy pose in the highest populated cluster of the AD4 covalent docking procedure (green) with the original inspiration fragment (pink) and the crystal structure (salmon). For every dock, the M$^{pro}$ protein structure of the corresponding inspiration fragment co-crystal structure was used. a) Crystal structure of Moonshot design X3077 (salmon) with fragment X0770 (pink) and the docked pose of X3077 (green). b) Crystal structure of Moonshot designed compound X3324 (salmon) with fragment X1380 (pink) and the docked pose of X3324 (green). c) Crystal structure of Moonshot design X3325 (salmon) with fragment X1386 (pink) and the docked pose of X3325 (green). d) Crystal structure of Moonshot design X10172 (salmon) with fragment X1382 (pink) and the docked pose of X10172 (green). e) Crystal structure of Moonshot design X10306 (salmon) with fragment X0770 (pink) and the docked pose of X10306 (green). f) Crystal structure of Moonshot design X10899 (salmon) with fragment X1458 (pink) and the docked pose of X10899 (green).

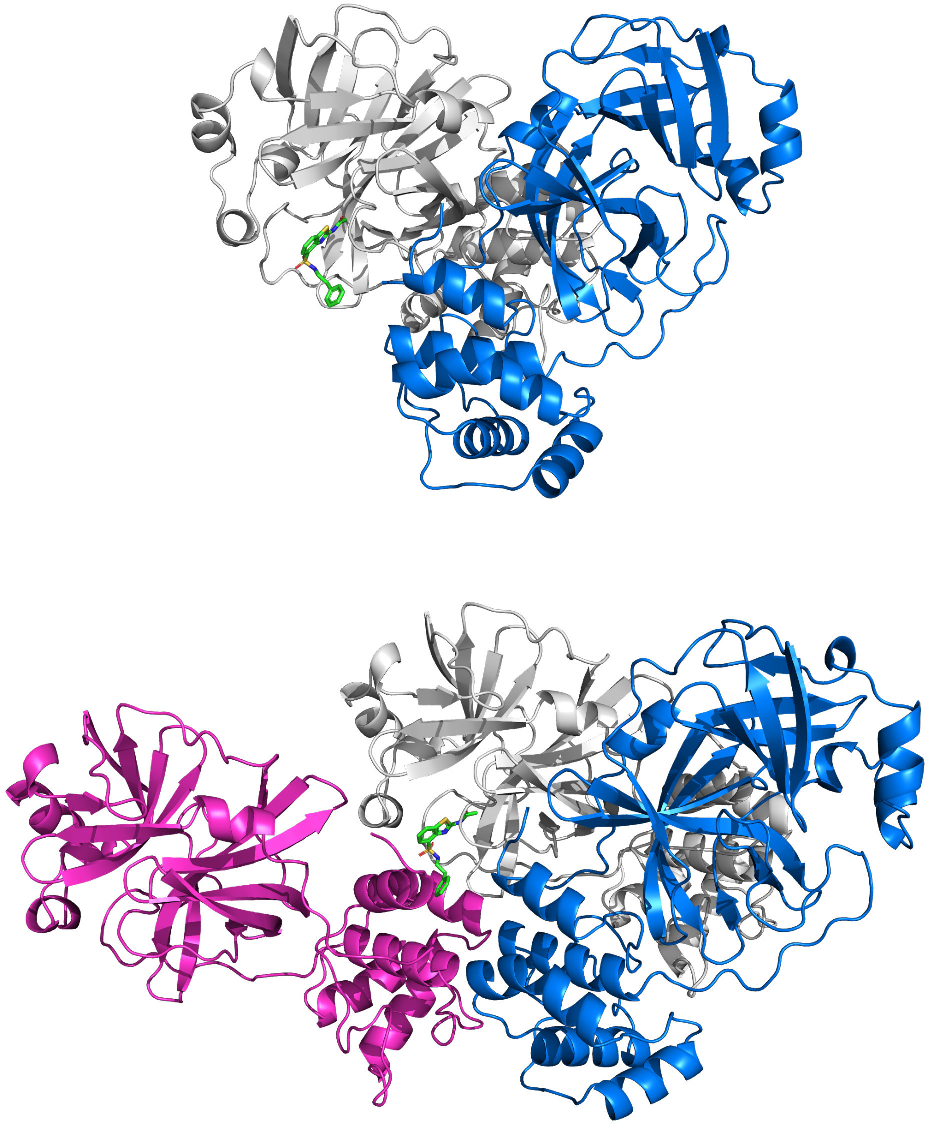

**Figure S4.10**: View from a crystal structure of X10899 (green) from the perspective of (top) the biologically relevant dimer and (bottom) one additional crystal packing symmetry mate. Chains A, B and C are coloured white, blue and pink respectively. X10899 is bound in the active site of chain A (white); its aromatic sidechain interacts with the symmetry-related chain C.

### Implications for future inhibitor design

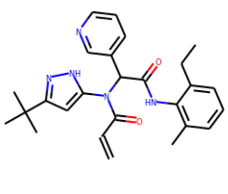

Mpro-x2705 IC50 [uM]: nan

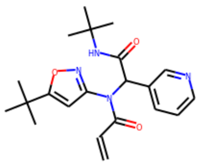

Mpro-x2540 IC50 [uM]: 99.0

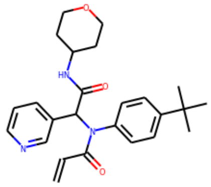

Mpro-x3117 IC50 [uM]: nan

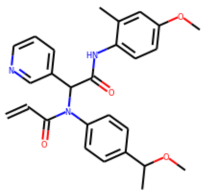

Mpro-x2776 IC50 [uM]: nan

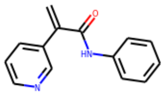

Mpro-x10022 IC50 [uM]: nan

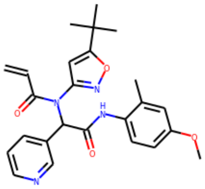

Mpro-x2694 IC50 [uM]: nan

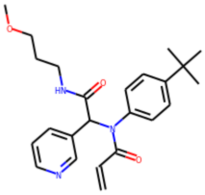

Mpro-x3113 IC50 [uM]: nan

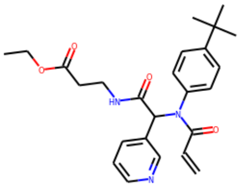

Mpro-x3110 IC50 [uM]: nan

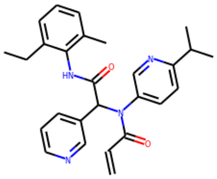

Mpro-x2703 IC50 [uM]: nan

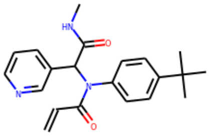

Mpro-x3124 IC50 [uM]: 35.71

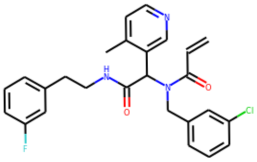

Mpro-x10082 IC50 [uM]: 99.5

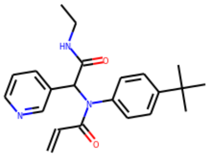

Mpro-x3359 IC50 [uM]: 45.49

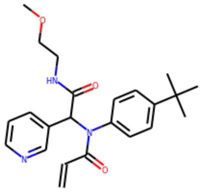

Mpro-x3115 IC50 [uM]: 55.78

**Figure S4.11**: Selection of all Moonshot compounds in cluster 5 that bind into the oxyanion hole. All compounds are covalent inhibitors, reacting with Cys-145 via the acrylamide warhead. $IC_{50}$ values are obtained from the postera.ai GitHub page.[36] nan = not a number, indicating that the compound has not been assayed.

`

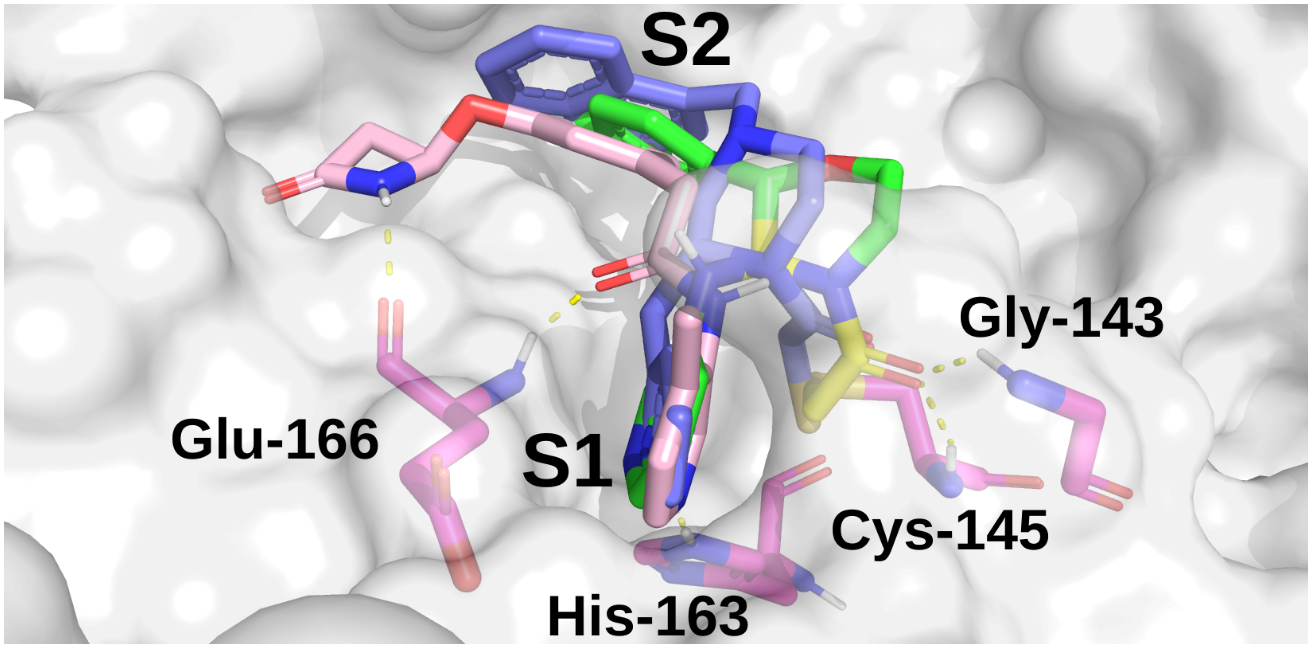

**Figure S4.12**: Overlay of the docked pose of FOC-CAS-e3a94da8-1 (green), the docked pose of MIH-UNI-e573136b-3 (blue), and a crystallographically observed binding mode of X10789 (salmon) with M$^{pro}$ (PDB: 5RER).[37] The proposed expansion of x10789 into the oxyanion hole is shown in yellow on compound FOC-CAS-e3a94da8-1.

### S5. Supplementary Information References

1. X. Xue, H. Yu, H. Yang, F. Xue, Z. Wu, W. Shen, J. Li, Z. Zhou, Y. Ding, Q. Zhao, X. C. Zhang, M. Liao, M. Bartlam and Z. Rao, *J. Virol.*, 2008, **82**, 2515-2527.
2. B. Webb and A. Sali, *Curr. Protoc. Bioinformatics*, 2016, **54**, 5.6.1-5.6.37.
3. M. H. M. Olsson, C. R. Søndergaard, M. Rostkowski and J. H. Jensen, *J. Chem. Theory Comput.*, 2011, **7**, 525-537.
4. Schrödinger LLC., Schrödinger Release 2020-4: Maestro New York, NY, 2020.
5. J. A. Maier, C. Martinez, K. Kasavajhala, L. Wickstrom, K. E. Hauser and C. Simmerling, *J. Chem. Theory Comput.*, 2015, **11**, 3696-3713.
6. W. L. Jorgensen, J. Chandrasekhar, J. D. Madura, R. W. Impey and M. L. Klein, *J. Chem. Phys.*, 1983, **79**, 926-935.
7. H. Grubmüller, V. Groll and P. Tavan, SOLVATE, Institute for Medical Optics, University of Munich, 1996-2013.
8. D. A. Case, I. Y. Ben-Shalom, S. R. Brozell, D. S. Cerutti, I. T.E. Cheatham, V. W. D. Cruzeiro, T. A. Darden, R. E. Duke, D. Ghoreishi, G. Giambasu, T. Giese, M. K. Gilson, H. Gohlke, A. W. Goetz, D. Greene, R. Harris, N. Homeyer, Y. Huang, S. Izadi, A. Kovalenko, R. Krasny, T. Kurtzman, T. S. Lee, S. LeGrand, P. Li, C. Lin, J. Liu, T. Luchko, R. Luo, V. Man, D. J. Mermelstein, K. M. Merz, Y. Miao, G. Monard, C. Nguyen, H. Nguyen, A. Onufriev, F. Pan, R. Qi, D. R. Roe, A. Roitberg, C. Sagui, S. Schott-Verdugo, J. Shen, C. L. Simmerling, J. Smith, J. Swails, R. C. Walker, J. Wang, H. Wei, L. Wilson, R. M. Wolf, X. Wu, L. Xiao, Y. Xiong, D. M. York and P. A. Kollman, AMBER 2019, University of California, San Francisco, 2019.
9. J.-P. Ryckaert, G. Ciccotti and H. J. C. Berendsen, *J. Comput. Phys.*, 1977, **23**, 327-341.
10. M. Gaus, Q. Cui and M. Elstner, *J. Chem. Theory Comput.*, 2011, **7**, 931-948.
11. J.-D. Chai and M. Head-Gordon, *Phys. Chem. Chem. Phys.*, 2008, **10**, 6615-6620.
12. P. C. Hariharan and J. A. Pople, *Theor. Chem. Acc.*, 1973, **28**, 213-222.
13. M. M. Francl, W. J. Pietro, W. J. Hehre, J. S. Binkley, M. S. Gordon, D. J. DeFrees and J. A. Pople, *J. Chem. Phys.*, 1982, **77**, 3654-3665.
14. M. J. Frisch, G. W. Trucks, H. B. Schlegel, G. E. Scuseria, M. A. Robb, J. R. Cheeseman, G. Scalmani, V. Barone, G. A. Petersson, H. Nakatsuji, X. Li, M. Caricato, A. V. Marenich, J. Bloino, B. G. Janesko, R. Gomperts, B. Mennucci, H. P. Hratchian, J. V. Ortiz, A. F. Izmaylov, J. L. Sonnenberg, Williams, F. Ding, F. Lipparini, F. Egidi, J. Goings, B. Peng, A. Petrone, T. Henderson, D. Ranasinghe, V. G. Zakrzewski, J. Gao, N. Rega, G. Zheng, W. Liang, M. Hada, M. Ehara, K. Toyota, R. Fukuda, J. Hasegawa, M. Ishida, T. Nakajima, Y. Honda, O. Kitao, H. Nakai, T. Vreven, K. Throssell, J. A. Montgomery Jr., J. E. Peralta, F. Ogliaro, M. J. Bearpark, J. J. Heyd, E. N. Brothers, K. N. Kudin, V. N. Staroverov, T. A. Keith, R. Kobayashi, J. Normand, K. Raghavachari, A. P. Rendell, J. C. Burant, S. S. Iyengar, J. Tomasi, M. Cossi, J. M. Millam, M. Klene, C. Adamo, R. Cammi, J. W. Ochterski, R. L. Martin, K. Morokuma, O. Farkas, J. B. Foresman and D. J. Fox, Gaussian, Wallingford, CT, 2016.
15. C. D. Owen, P. Lukacik, C. M. Strain-Damerell, A. Douangamath, A. J. Powell, D. Fearon, J. Brandao-Neto, A. D. Crawshaw, D. Aragao, M. Williams, R. Flaig, D. Hall, K. McAauley, D. I. Stuart, F. von Delft and M. A. Walsh, *PDB 6YB7*, 2020, DOI: 10.2210/pdb6yb7/pdb.
16. R. He, F. Dobie, M. Ballantine, A. Leeson, Y. Li, N. Bastien, T. Cutts, A. Andonov, J. Cao, T. F. Booth, F. A. Plummer, S. Tyler, L. Baker and X. Li, *Biochem. Biophys. Res. Commun.*, 2004, **316**, 476-483.
17. F. Wu, S. Zhao, B. Yu, Y.-M. Chen, W. Wang, Z.-G. Song, Y. Hu, Z.-W. Tao, J.-H. Tian, Y.-Y. Pei, M.-L. Yuan, Y.-L. Zhang, F.-H. Dai, Y. Liu, Q.-M. Wang, J.-J. Zheng, L. Xu, E. C. Holmes and Y.-Z. Zhang, *Nature*, 2020, **579**, 265-269.
18. R. C. Edgar, *Nucleic Acids Res.*, 2004, **32**, 1792-1797.
19. Schrödinger LLC., The PyMOL Molecular Graphics System, Version 2.3.0.
20. R. L. Dunbrack Jr. and F. E. Cohen, *Protein Sci.*, 1997, **6**, 1661-1681.
21. Molecular Operating Environment (MOE), (2019.01), Chemical Computing Group ULC, 1010 Sherbooke St. West, Suite #910, Montreal, QC, Canada, H3A 2R7, 2021.
22. W. D. Cornell, P. Cieplak, C. I. Bayly, I. R. Gould, K. M. Merz, D. M. Ferguson, D. C. Spellmeyer, T. Fox, J. W. Caldwell and P. A. Kollman, *J. Am. Chem. Soc.*, 1995, **117**, 5179-5197.
23. P. R. Gerber and K. Müller, *J. Comput. Aided Mol. Des.*, 1995, **9**, 251-268.
24. C. J. Williams, J. J. Headd, N. W. Moriarty, M. G. Prisant, L. L. Videau, L. N. Deis, V. Verma, D. A. Keedy, B. J. Hintze, V. B. Chen, S. Jain, S. M. Lewis, W. B. Arendall III, J. Snoeyink, P. D. Adams, S. C. Lovell, J. S. Richardson and D. C. Richardson, *Protein Sci.*, 2018, **27**, 293-315.
25. R. Anandakrishnan, B. Aguilar and A. V. Onufriev, *Nucleic Acids Res.*, 2012, **40**, W537-W541.
26. M. J. Abraham, T. Murtola, R. Schulz, S. Páll, J. C. Smith, B. Hess and E. Lindahl, *SoftwareX*, 2015, **1-2**, 19-25.
27. K. Lindorff-Larsen, S. Piana, K. Palmo, P. Maragakis, J. L. Klepeis, R. O. Dror and D. E. Shaw, *Proteins*, 2010, **78**, 1950-1958.
28. U. Essmann, L. Perera, M. L. Berkowitz, T. Darden, H. Lee and L. G. Pedersen, *J. Chem. Phys.*, 1995, **103**, 8577-8593.
29. B. Hess, H. Bekker, H. J. C. Berendsen and J. G. E. M. Fraaije, *J. Comput. Chem.*, 1997, **18**, 1463-1472.
30. X. Daura, K. Gademann, B. Jaun, D. Seebach, W. F. van Gunsteren and A. E. Mark, *Angew. Chem. Int. Ed.*, 1999, **38**, 236-240.
31. A. Onufriev, D. Bashford and D. A. Case, *Proteins*, 2004, **55**, 383-394.
32. P. Eastman, J. Swails, J. D. Chodera, R. T. McGibbon, Y. Zhao, K. A. Beauchamp, L.-P. Wang, A. C. Simmonett, M. P. Harrigan, C. D. Stern, R. P. Wiewiora, B. R. Brooks and V. S. Pande, *PLOS Comput. Biol.*, 2017, **13**, e1005659.
33. H. C. Jubb, A. P. Higueruelo, B. Ochoa-Montaño, W. R. Pitt, D. B. Ascher and T. L. Blundell, *J. Mol. Biol.*, 2017, **429**, 365-371.
34. H. C. Jubb, PDBTools GitHub: https://github.com/harryjubb/pdbtools, 2019.

35. J. Chodera, A. A. Lee, N. London and F. von Delft, *Nat. Chem.*, 2020, **12**, 581-581.
36. H. Achdout, A. Aimon, E. Bar-David, H. Barr, A. Ben-Shmuel, J. Bennett, M. L. Bobby, J. Brun, B. Sarma, M. Calmiano, A. Carbery, E. Cattermole, J. D. Chodera, A. Clyde, J. E. Coffland, G. Cohen, J. Cole, A. Contini, L. Cox, M. Cvitkovic, A. Dias, A. Douangamath, S. Duberstein, T. Dudgeon, L. Dunnett, P. K. Eastman, N. Erez, M. Fairhead, D. Fearon, O. Fedorov, M. Ferla, H. Foster, R. Foster, R. Gabizon, P. Gehrtz, C. Gileadi, C. Giroud, W. G. Glass, R. Glen, I. Glinert, M. Gorichko, T. Gorrie-Stone, E. J. Griffen, J. Heer, M. Hill, S. Horrell, M. F. D. Hurley, T. Israely, A. Jajack, E. Jnoff, T. John, A. L. Kantsadi, P. W. Kenny, J. L. Kiappes, L. Koekemoer, B. Kovar, T. Krojer, A. A. Lee, B. A. Lefker, H. Levy, N. London, P. Lukacik, H. B. Macdonald, B. MacLean, T. R. Malla, T. Matviiuk, W. McCorkindale, S. Melamed, O. Michurin, H. Mikolajek, A. Morris, G. M. Morris, M. J. Morwitzer, D. Moustakas, J. B. Neto, V. Oleinikovas, G. J. Overheul, D. Owen, R. Pai, J. Pan, N. Paran, B. Perry, M. Pingle, J. Pinjari, B. Politi, A. Powell, V. Psenak, R. Puni, V. L. Rangel, R. N. Reddi, S. P. Reid, E. Resnick, M. C. Robinson, R. P. Robinson, D. Rufa, C. Schofield, A. Shaikh, J. Shi, K. Shurrush, A. Sittner, R. Skyner, A. Smalley, M. D. Smilova, J. Spencer, C. Strain-Damerell, V. Swamy, H. Tamir, R. Tennant, A. Thompson, W. Thompson, S. Tomasio, A. Tumber, I. Vakonakis, R. P. van Rij, F. S. Varghese, M. Vaschetto, E. B. Vitner, V. Voelz, A. von Delft, F. von Delft, M. Walsh, W. Ward, C. Weatherall, S. Weiss, C. F. Wild, M. Wittmann, N. Wright, Y. Yahalom-Ronen, D. Zaidmann, H. Zidane and N. Zitzmann, *bioRxiv*, 2020, DOI: 10.1101/2020.10.29.339317, 2020.2010.2029.339317.
37. A. Douangamath, D. Fearon, P. Gehrtz, T. Krojer, P. Lukacik, C. D. Owen, E. Resnick, C. Strain-Damerell, A. Aimon, P. Ábrányi-Balogh, J. Brandão-Neto, A. Carbery, G. Davison, A. Dias, T. D. Downes, L. Dunnett, M. Fairhead, J. D. Firth, S. P. Jones, A. Keeley, G. M. Keserü, H. F. Klein, M. P. Martin, M. E. M. Noble, P. O'Brien, A. Powell, R. N. Reddi, R. Skyner, M. Snee, M. J. Waring, C. Wild, N. London, F. von Delft and M. A. Walsh, *Nat. Commun.*, 2020, **11**, 5047.
38. P. Jaccard, *New Phytol.*, 1912, **11**, 37-50.
39. H. Gohlke, C. Kiel and D. A. Case, *J. Mol. Biol.*, 2003, **330**, 891-913.
40. B. R. Miller, T. D. McGee, J. M. Swails, N. Homeyer, H. Gohlke and A. E. Roitberg, *J. Chem. Theory Comput.*, 2012, **8**, 3314-3321.
41. L. E. Ratcliff, W. Dawson, G. Fisicaro, D. Caliste, S. Mohr, A. Degomme, B. Videau, V. Cristiglio, M. Stella, M. D'Alessandro, S. Goedecker, T. Nakajima, T. Deutsch and L. Genovese, *J. Chem. Phys.*, 2020, **152**, 194110.
42. W. Dawson, S. Mohr, L. E. Ratcliff, T. Nakajima and L. Genovese, *J. Chem. Theory Comput.*, 2020, **16**, 2952-2964.
43. C. Bannwarth, E. Caldeweyher, S. Ehlert, A. Hansen, P. Pracht, J. Seibert, S. Spicher and S. Grimme, *WIREs Comput. Mol. Sci.*, 2021, **11**, e1493.
44. S. Spicher and S. Grimme, *Angew. Chem. Int. Ed.*, 2020, **59**, 15665-15673.
45. T. R. Malla, A. Tumber, T. John, L. Brewitz, C. Strain-Damerell, C. D. Owen, P. Lukacik, H. T. H. Chan, P. Maheswaran, E. Salah, F. Duarte, H. Yang, Z. Rao, M. A. Walsh and C. J. Schofield, *Chem. Commun.*, 2021, **57**, 1430-1433.
46. Y. Zhang and M. F. Sanner, *Bioinformatics*, 2019, **35**, 5121-5127.
47. Z. Jin, X. Du, Y. Xu, Y. Deng, M. Liu, Y. Zhao, B. Zhang, X. Li, L. Zhang, C. Peng, Y. Duan, J. Yu, L. Wang, K. Yang, F. Liu, R. Jiang, X. Yang, T. You, X. Liu, X. Yang, F. Bai, H. Liu, X. Liu, L. W. Guddat, W. Xu, G. Xiao, C. Qin, Z. Shi, H. Jiang, Z. Rao and H. Yang, *Nature*, 2020, **582**, 289-293.
48. J. Lee, L. J. Worrall, M. Vuckovic, F. I. Rosell, F. Gentile, A.-T. Ton, N. A. Caveney, F. Ban, A. Cherkasov, M. Paetzel and N. C. J. Strynadka, *Nat. Commun.*, 2020, **11**, 5877.
49. P. A. Ravindranath and M. F. Sanner, *Bioinformatics*, 2016, **32**, 3142-3149.
50. P. A. Ravindranath, S. Forli, D. S. Goodsell, A. J. Olson and M. F. Sanner, *PLOS Comput. Biol.*, 2015, **11**, e1004586.
51. W. Humphrey, A. Dalke and K. Schulten, *J. Molec. Graphics*, 1996, **14**, 33-38.
52. Diamond, Fragalysis, https://fragalysis.diamond.ac.uk/, (accessed 2020).
53. RDKit: Open-Source Cheminformatics Software, 2020.
54. G. M. Morris, R. Huey, W. Lindstrom, M. F. Sanner, R. K. Belew, D. S. Goodsell and A. J. Olson, *J. Comput. Chem.*, 2009, **30**, 2785-2791.
55. M. Wójcikowski, P. Zielenkiewicz and P. Siedlecki, *J. Cheminformatics*, 2015, **7**, 26.
56. S. Leung, M. Bodkin, F. von Delft, P. Brennan and G. Morris, *ChemRxiv*, 2019, DOI: 10.26434/chemrxiv.8100203.v1.
57. L. Zhang, D. Lin, X. Sun, U. Curth, C. Drosten, L. Sauerhering, S. Becker, K. Rox and R. Hilgenfeld, *Science*, 2020, **368**, 409-412.
58. D. W. Kneller, G. Phillips, H. M. O'Neill, R. Jedrzejczak, L. Stols, P. Langan, A. Joachimiak, L. Coates and A. Kovalevsky, *Nat. Commun.*, 2020, **11**, 3202.
59. Commissariat à l'Energie Atomique, Polaris(MD), http://biodev.cea.fr/polaris/index.html, 2018.
60. Y. Duan, C. Wu, S. Chowdhury, M. C. Lee, G. Xiong, W. Zhang, R. Yang, P. Cieplak, R. Luo, T. Lee, J. Caldwell, J. Wang and P. Kollman, *J. Comput. Chem.*, 2003, **24**, 1999-2012.
61. M. J. Field, M. Albe, C. Bret, F. Proust-De Martin and A. Thomas, *J. Comput. Chem.*, 2000, **21**, 1088-1100.
62. A. Krzemińska, P. Paneth, V. Moliner and K. Świderek, *J. Phys. Chem. B*, 2015, **119**, 917-927.
63. M. Wei, R. Wynn, G. Hollis, B. Liao, A. Margulis, B. G. Reid, R. Klabe, P. C. Liu, M. Becker-Pasha, M. Rupar, T. C. Burn, D. E. McCall and Y. Li, *J. Biomol. Screen.*, 2007, **12**, 220-228.
